# KOPTIC: A novel approach for *in silico* prediction of enzyme kinetics and regulation

**DOI:** 10.1101/807628

**Authors:** Wheaton L. Schroeder, Rajib Saha

**Author notes:** Wheaton L. Schroeder, University of Nebraska-Lincoln, Department of Chemical and Biomolecular Engineering, 209.1 Othmer Hall, Lincoln, NE 68588. Corresponding Author Rajib Saha, University of Nebraska-Lincoln, Department of Chemical and Biomolecular Engineering, 213 Othmer Hall, Lincoln, NE 68588.

## Abstract

Kinetic models of metabolism (kMMs) provide not only a more accurate method for designing novel biological systems but also characterization of system regulations; however, the multi-‘omics’ data required is prohibitive to their development and widespread use. Here, we introduce a new approach named **K**inetic **OPT**imization using **I**nteger **C**onditions (**KOPTIC**), which can circumvent the ‘omics’ data requirement and semi-automate kMM construction using *in silico* reaction flux data and metabolite concentration estimates derived from a metabolic network model to return plausible reaction mechanisms, regulations, and kinetic parameters (defined as ‘reactomics’) using an optimization-based approach. As a benchmark for the performance of KOPTIC, a previously published, four-tissue (leaf, root, seed, and stem) metabolic model of *Arabidopsis thaliana* was used, consisting of major primary carbon metabolism pathways, named p-ath780 (1015 reactions, 901 metabolites, and 780 genes). Data required for KOPTIC was derived from an Arabidopsis’ lifecycle of 61 days. Nine separate regulator restriction sets (allowing multiple solutions) defining KOPTIC runs hypothesized 3577 total regulatory interactions involving metabolic, allosteric, and transcriptional regulatory mechanisms (with nearly 40 verified by existing literature) with a median fit error of 13.44%. Flux rates of most KOPTIC fits were found to be significantly correlated with (93.6% with *p* < 0.05) and approximately 1:1 (*r* = 0.775, *p* ≪ 0.001) to the input time-series data. Thus, KOPTIC can hypothesize maps the regulatory landscape for a specific reaction, out of which the most relevant regulatory interaction(s) can be defined by the desired growth/stress conditions or the desired genetic interventions for use in the creation of kMMs.

---

The use of synthetic biology for the engineering of uni- and multi-cellular organisms to enhance desirable phenotypes in microbe, plant, and animal systems, is well established and is capable of affecting the lives of millions of individuals, such as in the case of artemisinin production in yeast or enhancing nutritional value of agricultural products [1][2]. Synthetic biology techniques have been applied to many plant systems such as tomatoes [3], rice [4], and maize [5] to produce enhanced phenotypes often with application to human nutrition [2], pest resistance [5], and resilience to abiotic stresses [6]. Many of these efforts focus on a genetic understanding and manipulation of the plant system (or plant tissue) in question, relying on intuitive interventions such as changes in regulation, insertion of new gene(s), and deletion of gene(s) from competing pathway(s) [2][5][6]. Alternatively, computational approaches based on stoichiometric genome-scale models (GSMs) of metabolism can be used to predict non-intuitive genetic interventions [7] by accounting for gene-protein-reaction (GPR) links, but also understand how a gene knockout, or a change in gene regulation, can affect the entire system through tools such as Flux Balance Analysis (FBA) [8], OptKnock [9], and OptForce [10]. Hence, these tools were reported to lead to enhanced mechanistic understanding for exploring the system-wide effects of genetic interventions especially in a microbial or a fungal system, such as *E. coli* [10], cyanobacteria [11], and yeast [12] as well as various plant species such as Arabidopsis [13][14], maize [15], sorghum [16], sugarcane [16], rapeseed [16], and rice [17].

Stoichiometric models are simpler (compared to kinetic models) steady-state representations of cellular metabolism and are widely used, since microbial cellular factories are often operated assuming a pseudo-steady state. This is a reasonable assumption since the time scale of metabolic reactions (fractions of seconds) is much quicker than other biological processes (such as transcription and translation which are on the order of minutes) [18]. Genetic interventions gained from stoichiometric modeling, while successful in many microbial applications, sometimes fail due to limitations of not incorporating reaction mechanisms, associated regulation, and enzyme metabolite concentrations [8][19]. Kinetic models of metabolism (kMMs) make up for the shortcomings of stoichiometric models at the expense of increased computational cost and ‘omics’ knowledge/data requirements. While kMMs should generate the same steady-state reaction fluxes as stoichiometric models, they are also able to model unsteady-state operation [18]. The ‘omics’ knowledge requirement includes transcriptomics, proteomics, and metabolomics to create accurate kMMs. If sufficient ‘omics’ data is available, deterministic or simulation-based kinetic modeling methods may be used, which while potentially accurate, form a system of stiff ordinary differential equations and require reasonable *in vivo*-relevant estimates for all kinetic parameters [7]. Another approach is Jacobian-based modeling, which makes local linear approximations of the kinetic system and can be used to calculate the time-scale of reactions and model stability from the eigenvalues of the Jacobian matrix. However, Jacobian-based modeling is more computationally complex than deterministic and simulation-based models and relies on some, if not all, *in vivo* kinetic parameters being known, in addition to knowledge of reaction mechanisms [7]. Since *in vivo* kinetic parameters are difficult to measure, Monte Carlo simulation-based modeling (particularly ensemble modeling), which estimates kinetic parameters, has become popular for the development of kMMs for prokaryotic organisms [7][18][19][20][21]. In this method, each reaction is decomposed to its elementary mass action steps, and then Monte Carlo simulation is used to produce a large number (ensemble) of kinetic parameter sets. These sets are pruned by training data sets until the best kinetic parameter estimate set is selected. Furthermore, no *in vivo* kinetic parameters are required *a priori* [19][20]. Despite this advantage, Monte Carlo methods are limited since the reaction mechanisms including the modes of regulations must be known [7] or *in vivo* mutant flux data must be available to verify hypothesized regulation [22].

It is this limitation (*a priori* knowledge or *in vivo* reaction flux data) which this current work seeks to address with an optimization-based tool capable of addressing the lack of *in vivo* knowledge of reaction mechanisms, regulations, and kinetic parameters, collectively hereafter defined as ‘reactomics’, which serve as a barrier to kMMs development for many species. This tool introduces a new approach for developing kMMs which is based on the use of stoichiometric models of metabolism, called <u>K</u>inetic <u>OPT</u>imization using <u>I</u>nteger <u>C</u>onditions, KOPTIC. As proof of validity of the underlying concept, KOPTIC is applied to a stoichiometric model of *Arabidopsis thaliana*, hereafter Arabidopsis, which was reconstructed in our recent study [23] as a model plant and a higher order biological system. Although Arabidopsis has the necessary ‘omics’ data to create a small core-metabolism kinetic model, this biological system is chosen because its metabolic regulation is well-studied, allowing ‘reactomic’ predictions made by KOPTIC to be verified. The KOPTIC approach, illustrated in Figure 1, uses Mixed Integer Non-Linear Programming (MINLP) and the data from the 61 time-points (described previously) to predict Arabidopsis ‘reactomics’. By circumventing the *in vivo* data requirements and automating kinetic model generation, KOPTIC can be used for rapid development of kMMs for poorly-studied organisms (those organisms with annotated genomes but little or no ‘reactomics’ data), thus broadening the usefulness of kMMs.

**Figure 1:**
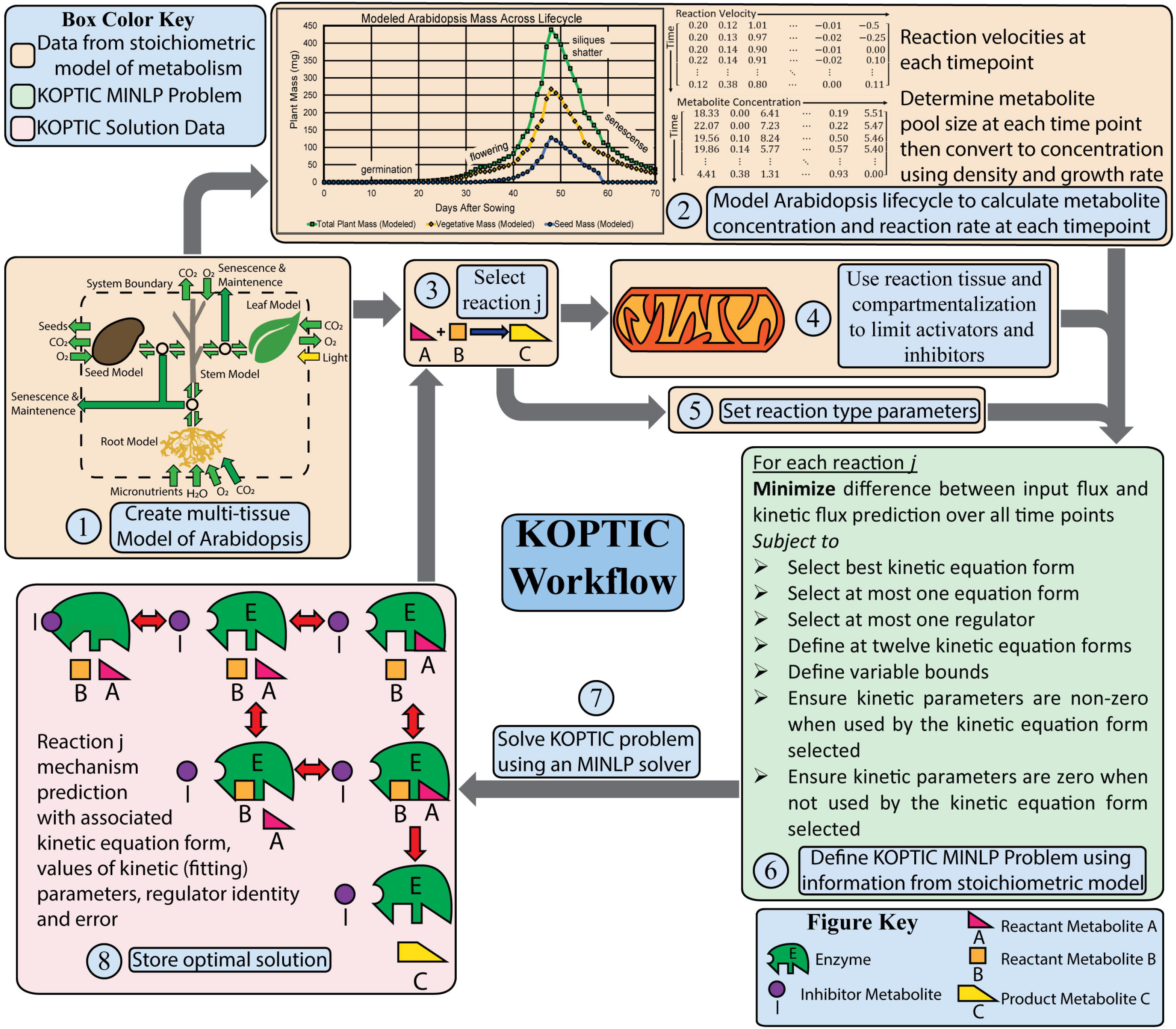
Workflow of the KOPTIC method. Much of this workflow is done by coding scripts. The brown boxes represent input data to KOPTIC, the green box represents the mixed integer non-linear programming (MINLP) optimization problem, and the pink boxes are the results obtained from solving the optimization problem. This workflow is repeated for each reaction (as KOPTIC solves on a per reaction basis). The collection of kinetic equations forms the basis a kinetic model of metabolism (kMM). Symbol definitions can be found in Supplemental File 1.

In the current work, a core stoichiometric metabolic model of Arabidopsis which was reconstructed in our recent study [23], consisting of major primary carbon metabolism pathways was used as the basis for the application of KOPTIC. This multi-tissue Arabidopsis stoichiometric model, referred to as p-ath780 has 1033 total (and 633 unique) reactions (R), 157 total (and 325 unique) metabolites (M), and 780 genes (G). The model p-ath780 consists of four tissue-level models of metabolism: leaf (R: 537, M: 479, and G: 703), root (R: 130, M: 126, and G: 250), seed (R: 428, M: 411, and G: 529), and stem (R: 160, M: 140, and G: 250) [23]. The tissues were linked and their respective environmental interactions described by a Flux Balance Analysis (FBA)-based [8][23] optimization framework [24][23]. These four tissues represent the core plant system with their essential metabolic roles: the root for nutrient uptake; the leaf for photosynthesis; the seed for metabolite storage and high metabolic investment; and the stem for metabolic transport, thus logically connecting these tissues. The optimization framework makes use of biologically relevant constraints on respiration, growth, photosynthesis, maintenance, senescence, and tissue ratios [25][26][27][28][29][30] in order to simulate flux values at each hour across the selected 61 day Arabidopsis lifecycle by using the p-ath780 model.

KOPTIC predicts ‘reactomics’ of each reaction using reaction type information from the stoichiometric model, such as specific number of substrates (single or dual) and reversibility (reversible or irreversible) and assumes three possible metabolite regulatory mechanisms for each reaction type: activation, inhibition, or no regulation. Kinetic equations derived for each reaction type combined with each metabolite regulatory mechanism, a total of 12 kinetic equation forms (see Supplemental File 1 for derivation of these equation forms), were then used by KOPTIC to fit each reaction from p-ath780 to a single kinetic equation form. This was done by minimizing the error between previously described time-point data and reaction flux predicted for that time point by a single kinetic equation form. The optimal solution for each reaction includes a ‘reactomic’ prediction as a mechanism, regulation, and kinetic parameters. To study various regulation mechanisms *in silico*, nine different regulatory restriction sets were devised and applied in nine separate KOPTIC runs. Each restriction set is a combination of one location and one identity restriction (see Table 1). These restrictions applied to metabolic regulators in separate KOPTIC runs allow for multiple ‘reactomic’ predictions for some reactions. Thus, the nine KOPTIC runs returned 3577 ‘reactomic’ predictions for the 594 reactions for which at least one solution was found. These solutions are hereafter referred as ‘fits’.

**Table 1:**
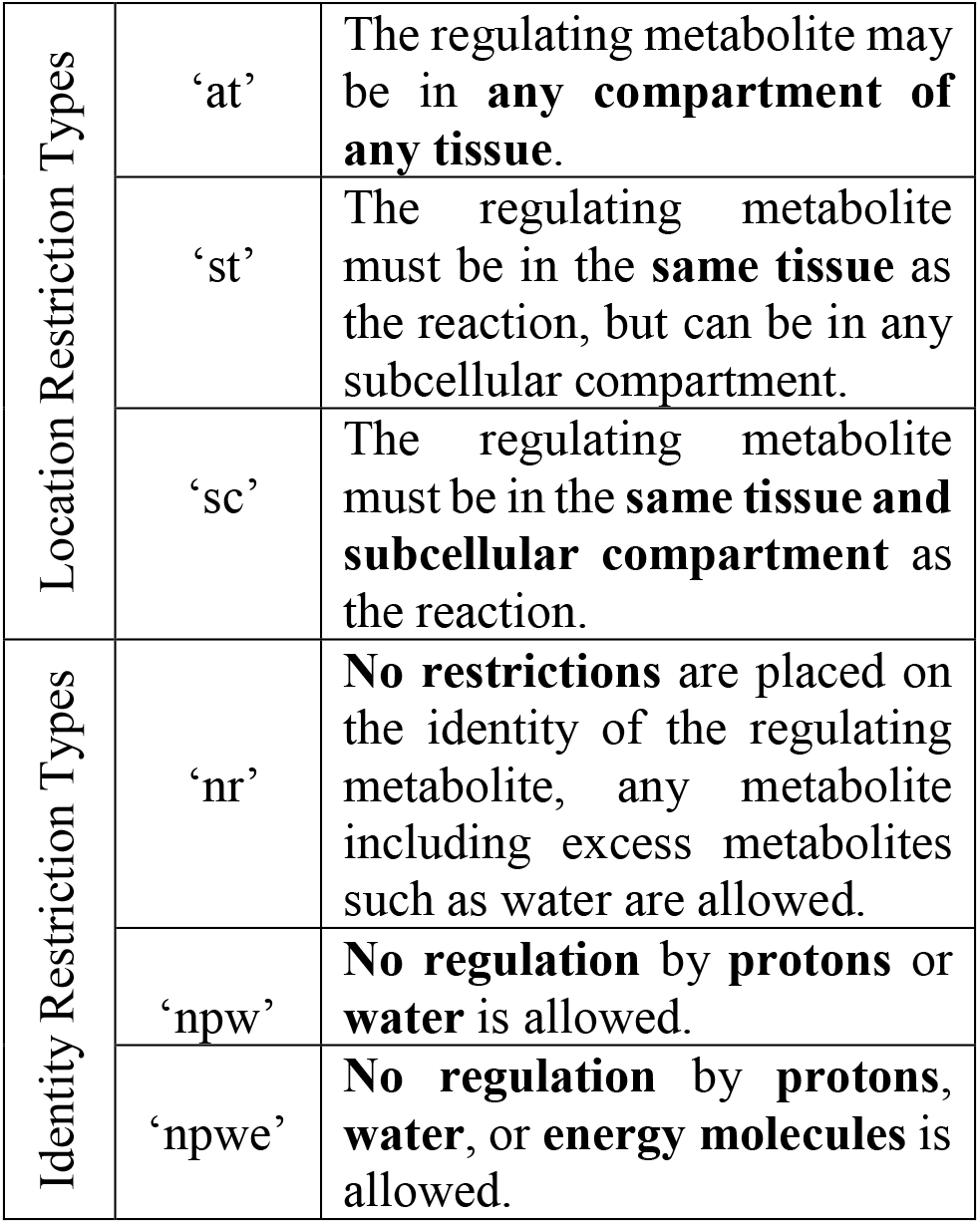
Restriction types used to create the nine KOPTIC restriction sets.

|  |  |  |
| --- | --- | --- |
| Location Restriction Types | ‘at’ | The regulating metabolite may be in **any compartment of any tissue**. |
|  | ‘st’ | The regulating metabolite must be in the **same tissue** as the reaction, but can be in any subcellular compartment. |
|  | ‘sc’ | The regulating metabolite must be in the **same tissue and subcellular compartment** as the reaction. |
| Identity Restriction Types | ‘nr’ | **No restrictions** are placed on the identity of the regulating metabolite, any metabolite including excess metabolites such as water are allowed. |
|  | ‘npw’ | **No regulation** by **protons** or **water** is allowed. |
|  | ‘npwe’ | **No regulation** by **protons, water, or energy molecules** is allowed. |

KOPTIC fits had a median error of 13.44% and particularly had low error when the regulating metabolite was limited to the same tissue as the reaction it acted upon (see Methods for details). To verify the qualitative accuracy of KOPTIC regulatory predictions, several predictions were compared to regulatory mechanisms reported in literature. We verified metabolic regulation predictions which include the ferredoxin/thioredoxin mechanism (fit errors ranging from near 0% to 37%), inhibition of ribose-5-phosphate isomerase by water-rich conditions (fit errors of 0.11% and 2.24%), and transcriptional regulation by nutrients such as sucrose, ammonia, and phosphate (fit errors ranging from 0.7% to 20.4%). These comparisons to experimental evidences demonstrate KOPTIC’s ability to predict correct metabolic regulations in response to abiotic stress and nutrient availability. In summary, this work shows how the KOPTIC approach can be used to semi-automatically (largely automated workflow), accurately (low fitting error), and correctly (correct regulatory mechanism) predict a variety of *in vivo* ‘reactomics’ through an *in silico* workflow that requires no foreknowledge of an organism’s *in vivo* ‘reactomics’ or ‘omics’ data.

## <u>K</u>inetic <u>OPT</u>imization using <u>I</u>nteger <u>C</u>onditions (KOPTIC)

KOPTIC’s first fit criteria for determining the ‘reactomics’ of each reaction was the reaction type as specified by the p-ath780 model based on the number of substrates (single- or dual-substrate) and reversibility of the reaction (irreversible or reversible). For each of these four reaction types, three possible metabolite regulatory mechanisms were assumed plausible: activation, competitive inhibition, or no regulation. Kinetic equations were then derived for each reaction type combined with each metabolite regulatory mechanism to yield 12 unique kinetic equation forms (see Supplemental File 1 for derivation of these equation forms). KOPTIC then uses MINLP optimization to attempt to fit each reaction from p-ath780 to a single kinetic equation form by minimizing the sum of squared error between the previously described time-point data and reaction flux predicted for that time-point by a single kinetic equation form of the 12 possible. The optimal solution includes ‘reactomic’ predictions as a reaction mechanism, regulation, and kinetic parameters are returned for each reaction for which at least one optimal solution was found. For each equation form, the *in silico* concentration of the regulator metabolite is multiplied by one or more *K*_*m*_(*j*) terms, which can take values ranging from 1*e*^−7^ to 1*e*^5^, such that the magnitude of a metabolite’s *in silico* concentration is of little or no importance in determining an optimal regulator for a given reaction. Instead, the pattern of a metabolite’s *in silico* concentration compared to a given reaction’s flux rate is of importance in determining whether a metabolite is an optimal regulator. More details on the formulation and creation of KOPTIC can be found in the Methods section and in Supplemental File 1.

To study various reaction mechanisms *in silico*, nine different regulatory restriction sets were devised and applied in nine separate KOPTIC runs. Each restriction set is a combination of one location and one identity restriction type (see Table 1). The location restriction types were same compartment (‘sc’), same tissue (‘st’), and any tissue (‘at’), while the identity restriction types were no restriction (‘nr’), no proton or water regulation (‘npw’), and no proton, water, or energy molecule regulation (‘npwe’). These restriction sets were applied to metabolic regulators in separate KOPTIC runs in order to allow multiple ‘reactomic’ prediction for some reactions, to explore how regulation changes by conditions, and to study multiple regulatory mechanisms for a single reaction. In order to make ‘reactomic’ predictions for as many model reactions as possible in a reasonable time, each of the nine separate KOPTIC runs (distinguished by its regulatory restriction set) had ten parallel instances, each starting with a reaction 10% of the way further through the model than the previous instance (so that each instance only predicts ‘reactomics’ for 10% of the model reactions for full coverage). The results of the ten parallel instances for each reaction set were concatenated into summaries of results for each of the nine reaction sets (Supplemental File 1) after a runtime of 168 hours (or 7 days) for each instance.

There were three KOPTIC results possible for each reaction: i) a ‘reactomics’ prediction, ii) no fit found, and iii) no fit attempted. The no fit found category occurs if the solver was unable to find a solution due to no solution space existing or the inability to find the solution space or heuristic termination with no suitable solution. The no fit attempted category is due to KOPTIC being unable to fit the reaction in question when the reaction has more than two reactants (53 reactions) or has no flux during the lifecycle of Arabidopsis (61 reactions serve as in-model documentation and are intentionally blocked). Therefore, KOPTIC could fit at most 891 reactions. From all the results obtained from all the restriction sets, there are 3577 unique kinetic equation fits for 594 of 891 total reactions (66.7%). To be defined as a unique kinetic equation fit, at least one of kinetic parameters, metabolic regulator, and kinetic mechanism needs to be unique. The complete set of these results are included in Supplemental File 2.

Figure 2B shows the average number of KOPTIC results (any output for a reaction) and number of ‘reactomic’ predictions for runs containing the same location or identity restriction type. As shown in Figure 2B, the ‘any tissues’ (‘at’) restriction type returned on average 100 fewer kinetic equation fits, even though it had approximately the same number of total reactions returned. This is likely because the binary solution space is significantly restricted by the latter two restriction types, specifically activator (Γ_ij_) and inhibitor (Ω_*ij*_) variables (see Supplemental File 1 for details). Binary variables Γ_*ij*_ and Ω_*ij*_ corresponding to regulators that are not allowed are fixed to 0 and treated as parameters, resulting in a quicker solution and more iterations before heuristic termination.

**Figure 2:**
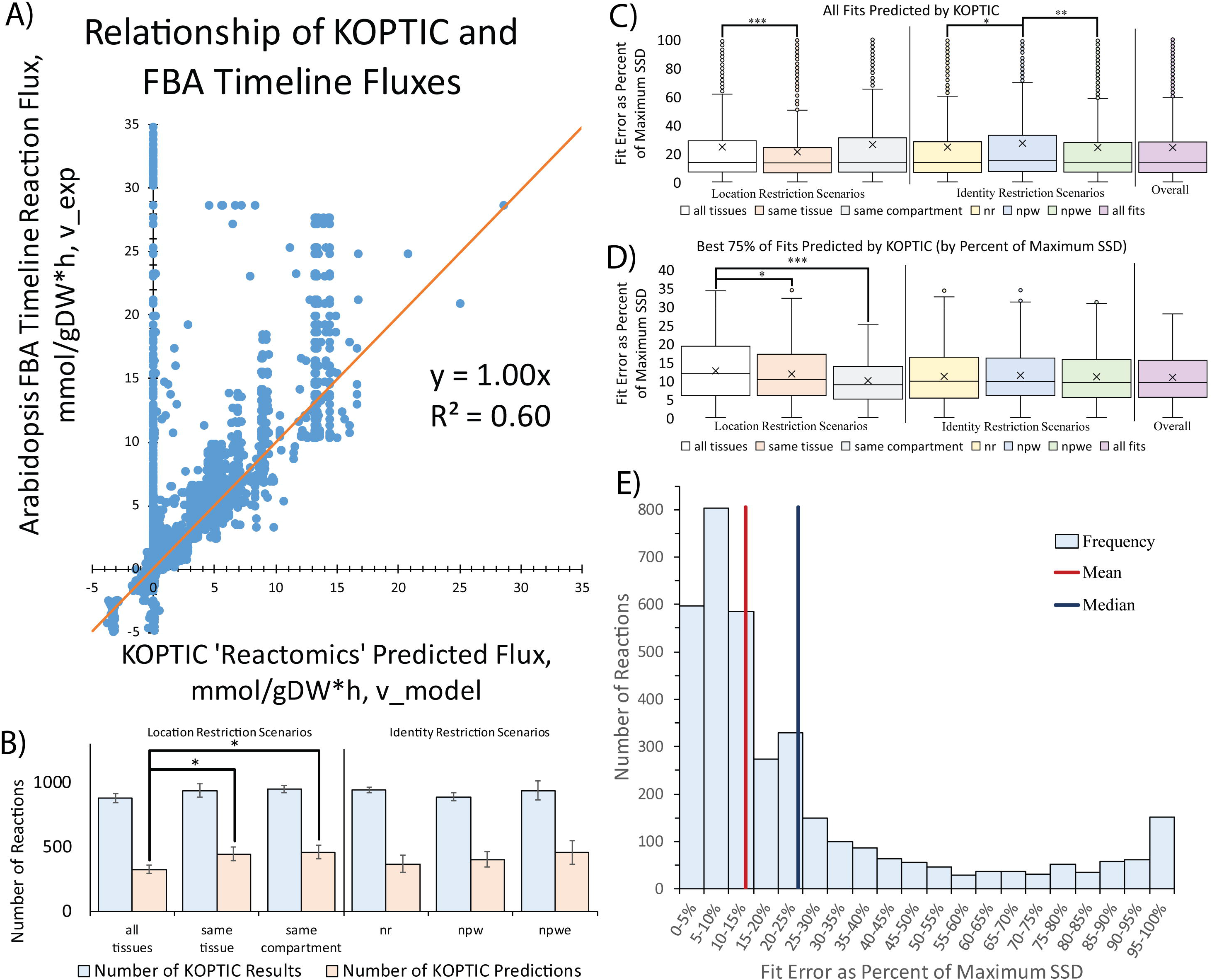
Statistical Analysis Graphs. A) Linear relationship between arabidopsis timeline fluxes and KOPTIC ‘reactomic’ flux predictions, including the squared Pearson’s correlation coefficient. B) Number of reactions returned by KOPTIC (number of reactions with any output) and number of fit kinetic equations returned by KOPTIC. Brackets and asterisks indicate statistically significant mean differences by the between-group ANOVA test. C) Shows the fit error of all KOPTIC predictions for each scenario type in terms of percent of maximum SSD. D) Shows the fit error of the best 75% of KOPTIC predictions, determined by percent of maximum SSD. E) Histogram of fit errors for all reactions fit by KOPTIC (counting multiple fits independently), along with the median and mean of all reactions fit. For A, B, and C, no comparison is made between location restriction scenarios (left) and identity restriction scenarios (right). * Represents p < 0.05, ** represents p < 0.01, *** represents p < 0.001.

Figure 2C shows the error of the fits returned by KOPTIC, which is the ratio of sum of squared differences of the kinetic mechanism fits to the maximum sum of squared differences (see Methods for finding how the sum of squared differences was utilized as an error measure). Full error statistics can be found in Supplemental File 2. The ‘same tissue’ (‘st’) restriction type was more accurate than the ‘any tissues’ (‘at’) restriction type, likely because of the increased number of fixed binary variables (as previously described, see Supplemental File 1). The ‘same compartment’ (‘sc’) restriction type had a standard deviation too high to show significant mean differences from either ‘at’ or ‘st’ restriction type. The ‘no proton or water’ (‘npw’) restriction type was the least accurate, and no significant difference was found between ‘no restriction’ (‘nr’) and ‘no proton, water, or energy molecule’ (‘npwe’) restriction types. Lower error for the ‘nr’ restriction type (compared to the ‘npw’ restriction type) might be due to capturing important abiotic stress regulations (e.g. osmotic and pH stress), while the lower error for the ‘npwe’ restriction type might be due to the restricted binary solution space (more fixed inhibitor and activator variables), allowing for more iterations. As many reaction fittings were heuristically terminated due to time, the accuracy of ‘npw’ was lower when compared to ‘npwe’ because the latter had more iterations in the time period allowed for solution.

The ‘sc’ restriction type had many reactions with very poor fits (more than 50 reactions with 90% fitting error or greater). Ignoring the poorest fits and considering the error of the best 75% of fits for each reaction type, shown in Figure 2D, the ‘sc’ restriction type had a significantly lower mean error than the ‘at’ restriction type, and had a lower standard deviation and a smaller interquartile range than any other restriction type. This suggests a bimodal distribution, with reactions being either well or poorly fit by the ‘sc’ restriction type. From Figures 2C and 2D, it is evident that the KOPTIC fitting error was positively skewed, with all 3577 KOPTIC predictions having a median error of 13.44% and a mean error of 24.10%, as shown in Figure 2E. Using Pearson’s correlation, it was found that the correlation between the flux rates predicted by KOPTIC ‘reactomics’ and the flux rate given by the Arabidopsis timeline was *r* = 0.775 (*p* ≪ 0.001). Additionally, 93.6% for KOPTIC ‘reactomic’ flux predictions had a significant correlation with their Arabidopsis timeline flux counterparts (e.g. same reaction, same timepoint, *p* ≤ 0.05). As noted in Figure 2A, the regression between Arabidopsis timeline fluxes (denoted *v*_*exp*_(*j*, *t*)) and KOPTIC ‘reactomic’ flux predictions (denoted *v*_*model*_(*j*, *t*)) was a straight line with a slope of 1 (e.g. generally *v*_*exp*_(*j*, *t*) = *v*_*model*_(*j*, *t*)).

To determine which types of reactions (low- or high-flux) were best fit by different restriction sets applied to KOPTIC, we determined the weighted mean sum of squared differences (as a measure of error) for each of the nine restriction sets and compared that value to the unweighted mean error. The weighted error used is *θ*_*SSD*_, and using this, we can say that low-flux reactions had better ‘reactomic’ predictions if *θ*_*SSD*_ > *μ*_*error*_, high-flux reactions had better ‘reactomic’ predictions if *θ*_*SSD*_ < *μ*_*error*_, and no significant difference in ‘reactomic’ predictions if *θ*_*SSD*_ ≈ *μ*_*error*_

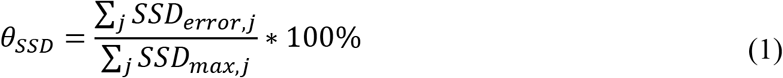

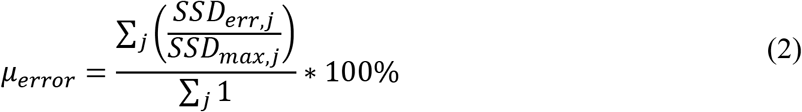

The location restriction type had a strong effect on what reactions were fit well by KOPTIC. For ‘at’ restriction type, low flux reactions were fit well and high flux reactions were fit poorly. This is elucidated by the values of *θ*_*SSD*_ for the three restriction sets including this restriction type being much higher, than the raw mean errors (*θ*_*SSD*_ = 86.44% and _error_ = 27.58% for ‘nr’/’at’, *θ*_*SSD*_ = 84.26% and *μ*_error_ = 26.52% for ‘npw’/’at’, and *θ*_*SSD*_ = 71.94% and μ_error_ = 26.68% for ‘npwe’/’at’). This conclusion also applied to the ‘nr’/’st’ restriction set which had better ‘reactomic’ prediction for low-flux reactions (*θ*_*SSD*_ = 55.62% *and μ*_*error*_ = 21.28%). There appeared to be no significant difference in goodness of ‘reactomic’ predictions for low- and high-flux reactions in the ‘npw’/’sc’ restriction set (*θ*_*SSD*_ = 26.30% and *μ*_*error*_ = 27.78%). High-flux reactions had better ‘reactomic’ predictions for the restriction sets ‘nr’/’sc’(*θ*_*SSD*_ = 2.70%; *μ*_*error*_ = 25.32%), ‘npw’/’st’ (*θ*_*SSD*_ = 12.78% and *μ*_*error*_ = 20.52%), ‘npwe’/’sc’ (*θ*_*SSD*_ = 11.12% and *μ*_*error*_ = 25.25%), ‘npwe’/’st’ (*θ*_*SSD*_ = 16.25% and *μ*_*error*_ = 21.19%).

## KOPTIC Predicted Regulations

### The Thioredoxin Regulatory Mechanism

The Thioredoxin (Trx) regulatory mechanism reversibly reduces disulfide bonds in target enzymes, changing the enzyme structure and increasing the level of activity of the desired enzyme. The first step in the mechanism is the reduction of thioredoxin by either NADPH or Ferredoxin. This is followed by the reduction of disulfide bond in the regulated enzyme through either a short-lived activation (Trx reduces the disulfide bond), or a longer-term activation (Trx reduces the disulfide bond by forming a complex with the target enzyme, see Figure 3A) [31][32][33][34][35][36]. This is a common and reversible mechanism of allosteric protein regulation in land plants and can help plants respond to oxidative stress [35][36]. Literature reports that in land plants, the ferredoxin as the initiator is generally limited to the chloroplast, and the NADPH as the initiator is generally identified in the cytosol and mitochondria [34][35]. However, Arabidopsis contains ferredoxin and ferredoxin reductase in mitochondria [37] as well as cytosolic ferredoxin [38], making the ferredoxin regulation mechanism plausible in mitochondrial, cytosolic or chloroplastic subcellular compartments.

**Figure 3:**
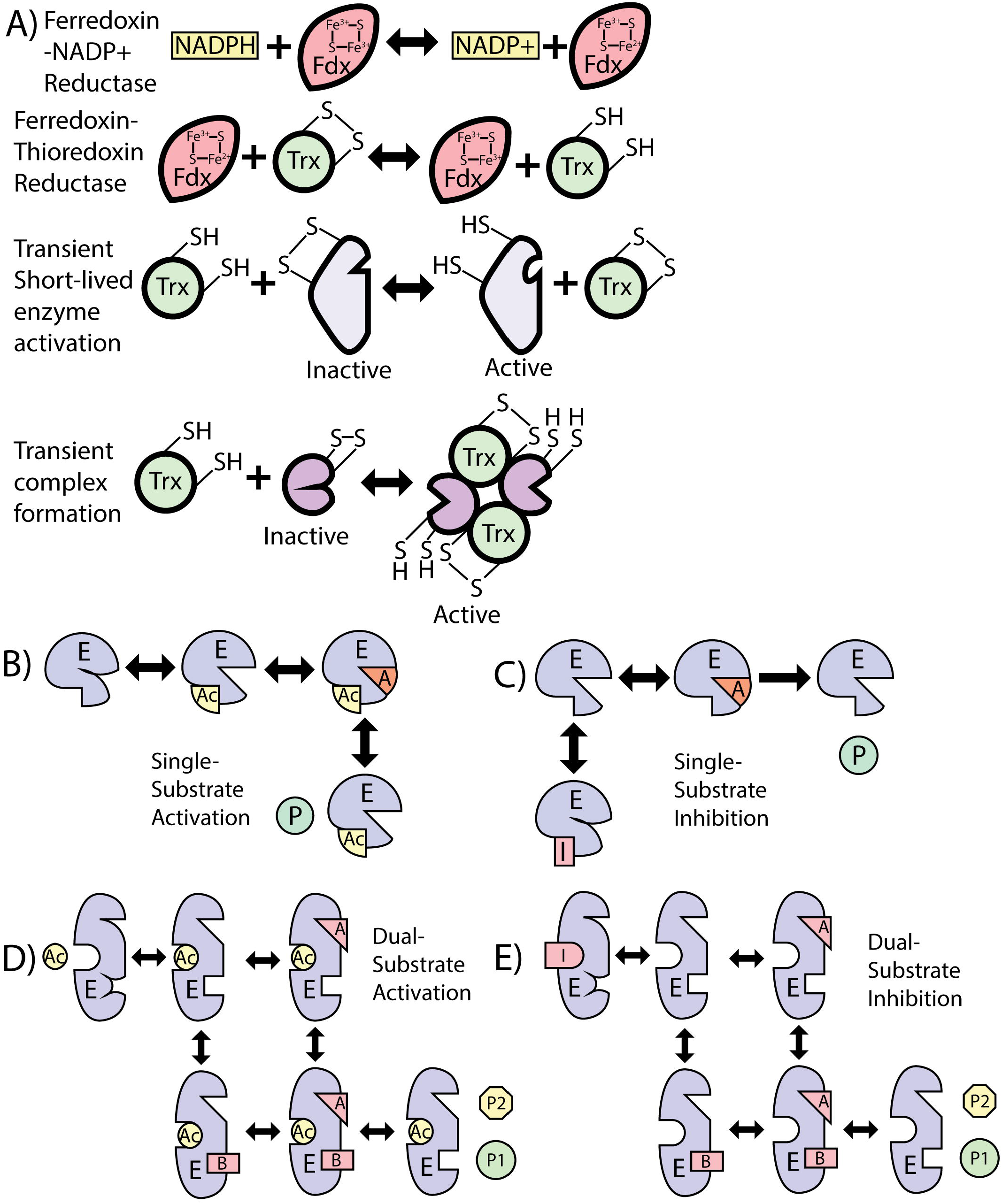
Kinetic Mechanisms. A) Mechanism of the thioredoxin enzyme regulation in *Arabidopsis*. The activation by reduced ferredoxin is reversible, and can activate the target enzyme by forming a complex with it or by reducing the disulfide bridges [31][35]. Figures B, C, D, and E are mechanisms used by KOPTIC for ‘reactomic’ predictions. B) A single-substrate irreversible enzyme reaction with activation. C) A reversible single-substrate irreversible enzyme reaction with inhibition. D) A dual-substrate reversible reaction with activation. E) A dual-substrate reversible reaction with inhibition

KOPTIC correctly predicted activation by reduced ferredoxin, inhibition by oxidized ferredoxin, activation by NADPH, and inhibition by NADP+ for several enzymes, of which selected predicted thioredoxin-mechanism regulation predictions are shown in Table 2 (complete list of predictions can be found in Supplemental File 2). KOPTIC’s kinetic equations use single-step regulation mechanisms (see Figure 3B, C, D, and E); therefore, the fit equations are simplifications of the actual mechanism, using single-step rather than multi-step regulation. All regulatory mechanisms were single substrate kinetics with activation (Figure 3B) or inhibition (Figure 3C) except for ATPase, which was modeled as irreversible dual-substrate kinetics with activation (Figure 3D). For instance, in Figure 3B we know that the activator (Ac) is reduced ferredoxin, which through ferredoxin-thioredoxin reductase forms reduced thioredoxin which in-turn activates dihydroxy-acid dehydratase (E) by reducing a disulfide bond. This allows the enzyme to act on 2,3-dihydroxy-3-methylbutanoate (A), to form 3-methyl-2-oxobutanoic acid (P). For this reaction, KOPTIC lumps the intermediate regulatory steps into a single step, but with low fit error (0.17%), giving confidence that the derived kinetic parameters returned by KOPTIC capture the net effects of the intermediate steps for this reaction. Other reactions were fit by low (<10%) or moderate to high error (27 to 37%), depending on the efficacy of the single model regulation step capturing the multi-step mechanism. It is likely that the low fit error cases are activated by the transient activation mechanism (Figure 3A). One enzyme with relatively high error (ATPase, 37%) has high error because Trx activates ATPase by forming an enzyme complex [39], resulting in significantly more complex reaction kinetics which are more difficult to fit with a single step. High fit error cases are likely mechanisms with complex activation. Generally, KOPTIC is more successful in simplifying regulation mechanisms to a single step when the regulation mechanism is less complex. Despite KOPTIC’s predictive success in the examples listed in Table 2, KOPTIC made some incorrect predictions. One is that NAD-glyceraldehyde-3-phosphatase (NAD-G3P) was predicted by KOPTIC to be inhibited by Ferredoxin^2+^, where literature data shows that the competing reaction, NADP-G3P, is instead activated by the Trx mechanism [35][40]. Additionally, aldehyde dehydrogenase was predicted by KOPTIC to be inhibited by Ferredoxin^2+^, when this enzyme was reported to be activated by the Trx mechanism [35].

**Table 2:** Selected KOPTIC predictions agreeing with literature data on the thioredoxin enzyme regulation mechanism.

| Tissue | Enzyme | Enzyme Compartment | Regulator | Regulation | Regulator Compartment | Error (% of maximum SSD) | Source(s) |
| --- | --- | --- | --- | --- | --- | --- | --- |
| Stem | Pyruvate dehydrogenase E2 component | Cytosol | Ferredoxin<sup>3+</sup> | Inhibition | Cytosol | 2.28x10<sup>-5</sup>% | [35] |
| Leaf | Dihydroxy-acid dehydratase | Cytosol | Ferredoxin<sup>2+</sup> | Activation | Cytosol | 0.17% | [35] |
| Leaf | Ketol-acid reductoisomerase | Cytosol | Ferredoxin<sup>2+</sup> | Activation | Cytosol | 0.17% | [35] |
| Stem | Dihydrolipoamide Dehydrogenase | Cytosol | Ferredoxin<sup>3+</sup> | Inhibition | Cytosol | 3.51% | [35] |
| Seed | Glucose-6-phosphate isomerase | Plastid | NADPH | Activation | Cytosol | 3.66% | [35] |
| Stem | UDP-glucose phosphorylase | Cytosol | Ferredoxin<sup>3+</sup> | Inhibition | Cytosol | 5.92% | [35] |
| Stem | Phosphogluco-mutase | Cytosol | Ferredoxin<sup>3+</sup> | Inhibition | Cytosol | 5.96% | [35] |
| Stem | Glucose-6-phosphate isomerase | Cytosol | NADP<sup>+</sup> | Inhibition | Cytosol | 7.74% | [35] |
| Seed | Pyruvate decarboxylase | Cytosol | NADPH | Activation | Cytosol | 27.02% | [35] |
| Seed | Triosephosphate iosmerase | Cytosol | NADPH | Activation | Cytosol | 30.63% | [35] |
| Seed | ATPase | Mitochondria | Ferredoxin<sup>2+</sup> | Activation | Mitochondria | 37.00% | [34][54][40] |

### Inhibition of R5PI by High Water Availability

Plant cells are able to respond to drought conditions via signaling enzymes which regulate the expression or activity of other enzymes in response. According to literature, osmotic stress (drought) activates sucrose nonfermenting-1-related protein kinase 2 (SnRK2, gene *at1G78290*) [41] which phosphorylates chloroplastic R5PI (gene *at3G04790*), increasing R5PI’s activity [42]. KOPTIC predicted that leaf chloroplastic ribose 5-phosphate isomerase (R5PI) was inhibited by extracellular water *in silico* concentration, with a fitting error of 2.24%. The predicted mechanism is shown in Figure 3C. This inhibition form was detected by KOPTIC because osmotic-stress signaling is likely the rate-limiting step in the signaling pathway which increases activity of R5PI. This is because SnRK2 is activated two minutes after the onset of osmotic stress and reached maximal activity level within 0.5 to 2 hours after the onset of osmotic stress [41]. This same R5PI gene is also located in non-green plastids (as part of the pentose phosphate pathway) [43]. KOPTIC predicted that stem plastid water inhibited leaf plastid R5PI with a fitting error of 0.11% of the maximum SSD, suggesting some cross-tissue drought signaling. The mechanism of this reaction is also shown in Figure 3C.

### Transcriptional Regulation by (CN)-Signaling

A plant cell has mechanisms for sensing carbon, nitrogen, and phosphate as signaling molecules, which allows cells to respond appropriately by increasing or decreasing gene transcription [44][45][46]. KOPTIC was able to capture microarray-verified transcriptional regulation [46] by sucrose, ammonia, and phosphate. Of a total of 11 predictions, a select set of 9 predictions are summarized in Table 3 (the full set can be found in Supplemental File 3). For these predictions, all kinetic equation fits returned by KOPTIC were either single-substrate kinetics with activation (Figure 3B, product-producing step is reversible or irreversible) or inhibition (Figure 3C, product-producing step is reversible or irreversible), with the exception of 6-phosphofructokinase which used dual-substrate kinetics with inhibition (shown in Figure 3E). The signaling pathway, transcription, and translation were “black-boxed” by the binding of the inhibitor (I) or the binding of the activator (Ac) step in the KOPTIC fit mechanisms, resulting in moderate error (6 to 20%) of fitting. As previously mentioned, it appears that KOPTIC is better at fitting less complex regulation mechanisms, therefore higher errors likely correspond to more complex transcriptional regulation.

**Table 3:** Selected KOPTIC regulatory predictions corresponding to transcriptional regulation of enzymes by nitrogen or carbon signaling.

| Tissue | Enzyme (Gene) | Enzyme Compartment | Regulator | Regulation | Regulator Compartment | Error (% of maximum SSD) | Source(s) |
| --- | --- | --- | --- | --- | --- | --- | --- |
| Seed | Ribose-5-phosphate isomerase | Cytosol | Phosphate | Inhibition | Cytosol | 0.70% | [55] |
| Seed | 1,4-alpha-glucan branching enzyme | Plastid | Phosphate | Inhibition | Plastid | 6.06% | [55] |
| Stem | Glucose-6-phosphate isomerase (*at4G24620*) | Cytosol | Sucrose | Activation | Cytosol | 6.61% | [46] |
| Seed | Fructose-bisphosphate aldose (*at4G26530*) | Plastid | Sucrose | Inhibition | Extracellular | 7.92% | [46] |
| Leaf | Fructose-1,6-bisphosphatase | Chloroplast | Phosphate | Inhibition | Chloroplast | 10.94% | [55] |
| Seed | Trehalose 6-phosphate phosphatase (*at4G22590*) | Cytosol | Sucrose | Activation | Cytosol | 12.34% | [46] |
| Stem | Phosphogulco-mutase (*at1G23190*) | Cytosol | Sucrose | Activation | Extracellular | 15.47% | [46] |
| Seed | 6-phosphofructokinase | Plastid | Phosphate | Inhibition | Cytosol | 15.92% | [55] |
| Root | 2,3-bisphosphoglycerate-independent phosphoglycerate mutase (*at1G09780*) | Cytosol | Sucrose | Activation | Cytosol | 20.40% | [46] |

### The TCA Cycle

KOPTIC predicted some correct regulation predictions (with low and high error), some close to correct predictions, and some unverifiable or incorrect predictions for the TCA cycle. Examples of correct predictions are outlined in Table 4. All of these reactions had predicted inhibitions mechanisms, shown in Figures 3C and 3E. Some predictions were made close to literature reported regulations, such as leaf succinate dehydrogenase was predicted to be inhibited by isocitrate (13% error, mechanism in Figures 3C) when succinyl-CoA ligase, the previous step in the TCA cycle, is inhibited by isocitrate [47]. Additionally, leaf aconitase was predicted to be inhibited by malate (11% error, Figure 3C), where the enzyme is known to be inhibited by the structurally similar oxalomalate [47] (as the latter metabolite not present in any tissue model in this work). Incorrect and/or currently unverifiable (due to no published *in vivo* evidence) regulations often predicted fumarate as a regulator for a variety of mitochondrial enzymes including aconitase, isocitrate dehydrogenase, and malate dehydrogenase (error ranges from 15 to 39%, mechanisms shown in Figures 3B and Figues 3C).

**Table 4:** Correct predictions of citric acid cycle regulation enzyme regulation made by KOPTIC.

| Tissue | Enzyme (Gene) | Enzyme Compartment | Regulator | Regulation | Regulator Compartment | Error (% of maximum SSD) | Source(s) |
| --- | --- | --- | --- | --- | --- | --- | --- |
| Stem | 2-Oxoglutarate dehydrogenase | Inner Mitochondria | NADH | Inhibition | Inner Mitochondria | 5.14% | [56] |
| Seed | Succinate dehydrogenase | Inner Mitochondria | Oxaloacetate | Inhibition | Outer Mitochondria | 24.89% | [47] |
| Seed | Fumarase | Inner Mitochondria | Pyruvate | Inhibition | Inner Mitochondria | 37.79% | [56] |
| Root | Isocitrate dehydrogenase | Inner Mitochondria | ATP | Inhibition | Outer Mitochondria | 38.63% | [47] |
| Leaf | Fumarase | Inner Mitochondria | Pyruvate | Inhibition | Inner Mitochondria | 78.72% | [47] |

## Discussion

In this work, a four core metabolic models of Arabidopsis tissues (leaf, root, seed, and stem) [23] was used to as a base stoichiometric model to which KOPTIC was applied. This model linked all four tissue models in a comprehensive Flux Balance Analysis (FBA), multi-level optimization framework, which allowed interactions inside and between of the plant tissues [23]. This framework then calculated the reaction flux vectors and also estimated *in silico* metabolite concentration (based on metabolite pool sizes) at 1464 time points, each separated by one hour, in the Arabidopsis lifecycle to simulate changes in reaction fluxes at various time points, of which 61 time points, each separated by 24 hours, were selected to apply KOPTIC to due to computational limitations. We applied our KOPTIC approach to the 61 Arabidopsis time points from p-ath780. KOPTIC found optimal fit solutions for 594 of a possible 891 (66.7%) reactions. A relatively low median error of fits (13.44%) suggests that KOPTIC is a viable method for predicting ‘reactomics’ from accurate stoichiometric models for *in silico* study of reaction kinetics and mechanisms, as well as for the development of kMMs. KOPTIC is a particularly promising method when the model builders have little experience with creating kMMs or when there is little regulatory information available, such as for understudied metabolic systems, as KOPTIC offers an *in silico* workflow for semi-automating the creation of kMMs that enable the discovery and study of regulatory mechanisms.

From the error analysis performed, we can see that the ‘sc’ restriction type can produce lower error (see Figures 2C) for many reactions and can produce superior fits for high-flux reactions (*θ*_*SSD*_ < *μ*_*error*_), while producing higher error for others. We hypothesize that when the ‘sc’ restriction type has low error it is accurately capturing some regulation with a regulatory metabolite acting directly on the enzyme; however, not all enzymes are directly regulated by a metabolite, resulting in a number of high-error predictions. Conversely, the ‘nr’ restriction type produces superior ‘reactomics’ for low-flux reactions (*θ*_*SSD*_ < *μ*_*error*_). The ‘st’ restriction type produces a balanced approach to predicting reactomics, generally favoring neither low-nor high-flux reactions.

From the predicted regulation case studies discussed here, it was shown that the KOPTIC predictions can correctly predict abiotic stress responses (such as drought), multi-step allosteric regulation mechanisms (such as the Trx mechanism), and transcriptional regulation (such as (CN)-signaling). Generally, the less complex the regulatory mechanism predicted was, the higher the was the accuracy of the KOPTIC ‘reactomic’ fit. KOPTIC currently predicted TCA cycle regulation with mixed accuracy. When close-to-true or incorrect regulatory mechanisms were predicted by KOPTIC, they were often reasonable. For instance, leaf succinate dehydrogenase was predicted to be inhibited by isocitrate, but literature showed that succinyl-CoA ligase is inhibited by isocitrate [47] instead. It is reasonable that the inhibition of the immediately upstream reaction would result in a lower flux for the reaction catalyzed by succinate dehydrogenase. Furthermore, it is reasonable for KOPTIC to conclude that malate inhibits aconitase in the absence of oxalomalate in the model [47] as these metabolites are structurally similar.

Despite these successes, there is room for improvement in KOPTIC, for instance, the TCA cycle had many incorrect predictions or correct predictions with high error. A common incorrect (or unverifiable) prediction was predicting the regulation of TCA cycle reactions by fumarate. We hypothesize the fumarate was a common prediction because in the mitochondria of the tissue models, only TCA and oxidative phosphorylation pathways occurred. Therefore, for the ‘sc’ restriction type, a reaction in these pathways must be regulated by a metabolite in these pathways. Fumarate might have been optimal because it is the metabolite in the TCA cycle before malate and oxaloacetate. Both of these metabolites can be transported into or out of the mitochondria. Therefore, the *in silico* concentration of fumarate would be a better indicator of the rate of flux through the TCA cycle and would yield ‘reactomic’ predictions with lower error.

Furthermore, some reaction regulation predictions did not make sense and/or were incorrect. These were often due to limitations of the solver options used, in that the solver terminated when the absolute solution gap was less than 1*e*^−5^. When the reactions have small fluxes throughout the lifetime of the plant (∑_*j*_ *SSD*_*max,j*_ < 0.01), the solution method would often reach the termination criteria either in preprocessing or in the first few iterations, accepting one of the first potential regulators found. Simply reducing the absolute optimality criteria would result in the problem being mostly addressed for low flux reactions. However, this exacerbates the solution time for high flux reactions as the optimality criteria will be met only at very low error (≪ 1%) which will take considerable time to converge, significantly increasing KOPTIC run time as termination will rely on a time-based heuristic. Instead, a scaling factor will be applied to the KOPTIC objective function in future versions for heuristic termination at a fixed SSD error percentage. This will ideally not only fix the problem of high error associated with low flux reactions but also increase KOPTIC solution speed for high flux reactions and will allow user-defined error thresholds.

In addition, we will seek to sophisticate KOPTIC in order to increase its predictive capabilities and the number of reactions fit, as well as the optimize goodness of fit. One promising direction is to solve first by restricting possible regulators to the same compartment, then widen the location restriction on the regulator if a poor fit is achieved. Additionally, we will consider the likelihood of a metabolite being a regulator and use that information to point KOPTIC toward a more reasonable regulator earlier in the solution process. Further, we will expand the set of kinetic equations from which KOPTIC has to choose in order to make ‘reactomic’ predictions for reactions with more than two substrates. Moreover, effectively fixing *v*_*max*_ in the Michealis-Menten equation, we assume constant (or near constant) enzyme level. This assumption may be driving fit error, and therefore in future iterations of KOPTIC we will allow the *in silico* enzyme concentration to vary across time or condition. In addition, we will seek to decrease the computational cost of KOPTIC so that more data may be used and that KOPTIC solutions might be more quickly achieved.

KOPTIC will be used in future to develop condition-specific kinetic models of metabolism. By analyzing the ‘reactomic’ predictions for each reaction, we can choose to accept, reject, or seek validation for each. The set of ‘best reactomics’ (as defined by the model curator) for each reaction can be concatenated into a kinetic model of metabolism. The ‘best reactomics’ may be defined by literature validation of reaction mechanism or kinetic parameters. Alternatively, the ‘best reactomics’ may be defined as those corresponding to the restriction set which is most relevant for an organism, the desired growth conditions, or the desired genetic inetrventions. For instance, the ‘nr’ restriction type would be preferable when studying metabolic response to drought or pH stress conditions.

## Methods

### Development and Use of the P-ath780 Model

The p-ath780 model was developed in detail in our recent study [23]. In summary, this Arabidopsis model was developed in order to address the limitations of current stoichiometric models of metabolism which only take a single “snapshot” of organism metabolism which may not be suitable for organisms whose growth cannot be held at some steady state condition (such as multi-cellular organisms). By taking a series of “snapshots” of organisms metabolism across its lifecycle, using a Flux Balance Analysis (FBA) based approach, a more accurate and holistic picture of organism metabolism can be obtained. As with the KOPTIC tool, Arabidopsis was chosen for this work as a model organism [13]. The p-ath780 model focuses on the core-carbon metabolism of Arabidopsis and models seven distinct growth stages across 61 days of growth, taking “snapshots” of metabolism at one-hour intervals. The p-ath780 model agreed well with published literature data including mass yield, maintenance costs, senescence costs, and whole-plant growth checkpoints [23]. In this current work, the reaction flux rates at each “snapshot” was used in part as data input to KOPTIC as the target reaction rate fluxes of the fit kinetic equations. Specifically, due to the computational cost of the KOPTIC method at present, only 61 of the “snapshots” were used, one representing each day of the Arabidopsis lifecycle.

### Calculation of *in silico* Metabolite Concentration

In order to estimate *in silico* metabolite concentration, in a specific tissue, we first calculated the metabolic pool size [48] for each of the metabolites from the corresponding tissue models. *In silico* metabolite concentration represents an estimate of the concentration of a given metabolite in a given tissue or compartment, based on the summation of flux of reactions through that metabolite that is converted to concentration unit. The conversion was done using tissue growth rate (as a dilution factor) and tissue density (as a volume estimate from *in silico* plant mass). This follows from the assumptions that the flux through a metabolite will be greater in a metabolite with higher *in vivo* concentration, and that this estimate can be used in place of an *in vivo* concentration measurements in reaction kinetics. We further assumed that each sub-cellular compartment grows at the same rate as the tissue, that metabolite concentration is uniform in a subcellular compartment, and that each subcellular compartment is of the same density as the tissue. While these assumptions are oversimplifications of an *in vivo* system, they were necessary for *in silico* representation as quantitative *in vivo* data necessary to drop these assumptions is not available.

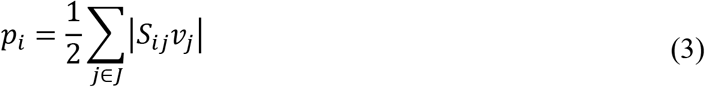

Equation (2) provides an estimate of the availability of the metabolite in a given tissue system in units of *mmol/gDW/* * *h*. Here, *p_i_* is the metabolite pool size of metabolite *i*,*S*_*ij*_ is the stoichiometric coefficient of metabolite *i* in reaction *j*, and *v_j_ is* the flux of reaction *j* in which *i* participates as a reactant or product. We converted this pool size value to an estimate of *in silico* metabolite concentration by using *in silico* biomass growth rate of the specific tissue (*v*_*biomass,tissue*_) and the tissue density (*ρ*_*tissue*_) [25][26][49][50][51][52]. To this end, the following conversion was used:

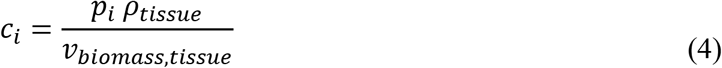

This conversion provided the estimate for all *in silico* metabolite concentration estimates used by KOPTIC.

### Development of KOPTIC

The KOPTIC method development, logic, derivation, symbol definition, and equations used can be found in Supplemental File 1, and the KOPTIC workflow is shown in Figure 1. In summary, we developed KOPTIC to study and predict kinetics of any biological system and to eventually develop kinetic models based on computational (such as FBA) or experimental (such as MFA) datasets. As previously discussed, KOPTIC uses twelve kinetic equation forms, from four reaction types with three possible types of regulation each, to find an optimal fit of the experimental data by one of these equation forms. KOPTIC returns ‘reactomic’ data of kinetic equation, kinetic parameters, and regulatory information. This is accomplished through an objective function that minimizes the sum of squared differences between the flux of reaction *j* as derived from the kinetic model *v*_*model*_(*j*, *t*), and the corresponding known (i.e., MFA) or calculated (i.e., FBA) reaction flux input into KOPTIC, assigned to parameter set *v*_exp_(j, t). Variable *v*_*model*_(*j*, *t*) and parameter *v*_exp_(j, t) are calculated for each time point or condition *t* in the set of time points or conditions *T*. KOPTIC is parallelizable in that the optimization formulation is solved for each input reaction independently (as solving all reactions at the same time is impractical as of yet due to computational time and cost), with the following objective function:

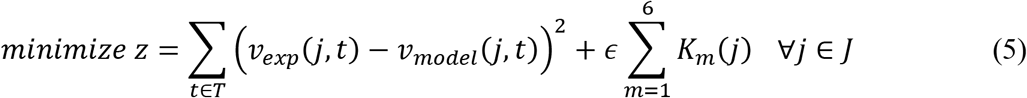

Where *K*_*m*_, *m* = [1,6], *m* ∈ ℤ is the set of kinetic parameters which are optimized to improve the fit of *v*_*model*_(*j*, *t*). There are at most six *K*_*m*_ parameters used to improve the fit of *v*_*model*_(*j*, *t*) (see Supplementary File 2). The modeled flux is defined as:

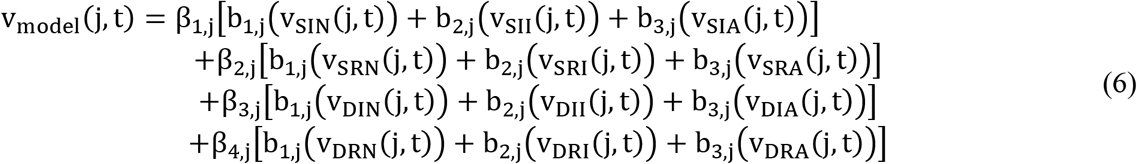

Where *β*_*u,j*_ are binary parameters defined by the stoichiometric model, in this case p-ath780, and restricted to β_1,j_ + β_2,j_ + β_3,j_ + β_4,j_ = 1. Parameter β_1,j_ = 1 corresponds to a single-substrate irreversible (SI) reaction, β_2,j_ = 1 corresponds to a single-substrate reversible reaction, β_3,j_ = 1 corresponds to a dual-substrate irreversible (DI) reaction, and β_4,j_ = 1 corresponds to a dual-substrate reversible (DR) reaction. Parameters *β* were set as parameters, rather than being combined with *b* variables in a single variable, to reduce the number of binary variables used in the formulation, which decreases solution time. Binary variables b_*y*,*j*_ are defined by optimization, as KOPTIC chooses the optimal regulatory mechanism. As with *β*_*u,j*_ parameters, b_1,j_ + b_1,j_ + b_1,j_ = 1, limiting KOPTIC to selecting a single regulatory mechanism. While often enzymes have multiple regulators, only a single regulator is allowed in the current formulation because of the form of the 12 kinetic equations derived (a new equation must be derived for each additional regulator). A single regulator equation form, with restriction sets being used to identify multiple possible regulators acting independently, allows identification of multiple regulators of a single enzyme. Variable *b*_1,*j*_ = 1 corresponds to no (N) regulation, *b*_2,*j*_ = 1 corresponds to inhibition (I) regulation, and *b*_3,*j*_ = 1 corresponds to activation (A) regulation. This forces v_model_(j, t) to equal exactly one of the kinetic forms. For simplicity, reactions with more than two substrates were not included due to the complexity of the kinetic equation forms and other regulation scenarios were not considered.

To understand how the ‘reactomics’ are predicted, consider if the optimal ‘reactomics’ of a reaction is single-substrate irreversible kinetics with no regulation (SIN, *β*_1,*j*_ = *b*_1,*j*_ = 1), then v_model_(j, t) is defined as below.

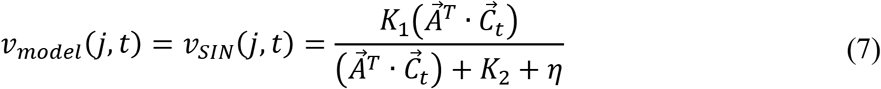

Where 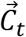 is a concentration vector from the Arabidopsis lifecycle FBA or from the MFA measurements, 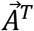 is a vector of unit magnitude which points at the substrate, *K*_1_ (akin to *v*_*max*_, the maximum reaction flux in the Michaelis-Menten equation) and *K*_2_ (akin to *K*_*M*_, the Michealis-Menten constant) are fitting parameters, and η is a very small number (here *η* = 1e^−7^) used to prevent errors when 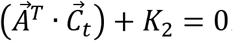. The objective function term involving K_m_(j) is used to prevent non-unique solutions resulting from multiple sets of K_m_(j) values yielding the same sum of squared differences. This term has minimal effect on the optimal solution in that *ϵ* is an arbitrary small number *ϵ* = 1e^−7^. Further constraints applied to the optimization problem include twelve constraints to define each of the twelve kinetic equation forms, six constraints to ensure that *M* ≥ *K*_*m*_(*j*) ≥ *η*, where *M* = 1*e*^5^, when used in the optimal kinetic equation form, four constraints to fix *K*_*m*_(*j*) = 0 when not used in the optimal kinetic equation form, and three constraints to ensure that a single kinetic equation form is selected and that only one metabolite is chosen as the optimal regulator. Because of the large range of possible values which *K*_*m*_(*j*) may take (spanning 12 order of magnitude), and also regulation forms having terms in which the *in silico* concentration of the regulator metabolite is modulated by one or more *K*_*m*_(*j*) values, the magnitude of the *in silico* concentration of any metabolite relative to that of the reaction is largely immaterial. The pattern of *in silico* concentration of the metabolite to the pattern of reaction flux is more important in determining an optimal metabolic regulator.

### KOPTIC Workflow

We used the 61 time point FBA-derived reaction fluxes and *in silico* metabolite concentration estimates from p-ath780 as input data for KOPTIC, as shown in Figure The KOPTIC formulation and symbols used in Figure 1 is discussed in the previous section and full details can be found in Supplemental File 1. Each KOPTIC run is restricted by one of the nine restriction sets (each set is a unique combination of identity and location restriction type, see Table 1) in order to identify multiple feasible combinations of regulating a metabolite and its location. This is advantageous as from the results we can choose the most plausible or best fitting kinetic equation form (as explained earlier). Each of the nine runs had 10 parallel instances starting at staggered model reactions to increase solution speed. This staggering is necessary as KOPTIC does not find solutions for most reactions in the model in a seven-day timeframe. Therefore, we can take these parallel instances and concatenate the results to get full coverage of the model (so that KOPTIC returned something for every reaction). The KOPTIC formulation was solved using BARON, an MINLP solver on the Generic Algebraic Modeling System (GAMS) [53], and each reaction solution yielded ‘reactomic’ predictions and model error (as a percentage of maximum SSD). We allowed 168 hours of runtime for each parallel instance, and when finished we concatenated the results of the appropriate instances into the results for each run.

### Error of Kinetic Fits by KOPTIC

Errors in kinetic equation fittings made by KOPTIC were described as a percentage of the maximum sum of squared differences. Equations used to describe error are shown below, where *T* is the set of 61 time points in the Arabidopsis lifecycle:

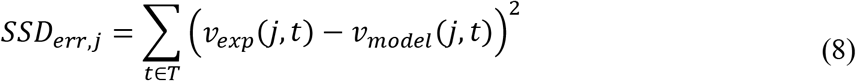

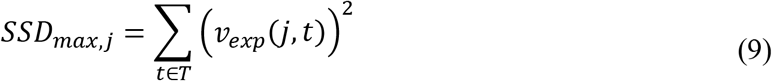

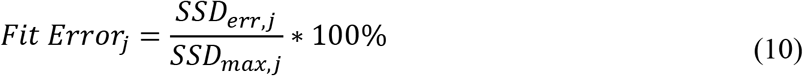

This *Fit Error*_*j*_ was used in the statistical analysis of this work to determine how well KOPTIC fit the 61 timepoint data given with the predicted ‘reactomics’.

### Statistical Analysis of Error

All statistical tests were performed using a between-group ANOVA analysis with a significance cutoff of *α* = 0.05. See Supplementary Text S3 for test statistic values and p-values of the statistical tests done.

## Acknowledgement

This work was completed utilizing the Holland Computing Center of the University of Nebraska, which receives support from the Nebraska Research Initiative. The authors gratefully acknowledge funding from UNL Faculty Startup Grant 21-1106-4038.

## Author Contributions

Experiments were conceived by R.S. and W.L.S. W.L.S. performed the experiments and analyzed the data. R.S. and W.L.S. contributed analysis tools. R.S. and W.L.S. wrote the manuscript.

**Supplementary Information** is linked to the online version of the paper.

