## Supplemental File 1 for "KOPTIC: A novel approach for *in silico* prediction of enzyme kinetics and regulation"

**KOPTIC Formulation**

Written by: Wheaton L. Schroeder and Rajib Saha

### **1 Premise**

The basic premise is that the best of twelve available kinetic equations is selected for a kinetic model in order to minimize the discrepancy between fluxomic data and model output. The resulting output from KOPTIC gives the optimal metabolite regulator (which best explains the experimental or FBA data), the kinetic (fitting) parameters, and the best fitting kinetic mechanism.

#### **1.1 Optimization formulation**

Variables and parameters used in the formulation of KOPTIC are given in the table below.

**Table 1:** Definitions of symbols used in the preliminary formulation of KOPTIC as outlined by Table 1. Variables here are unknown quantities which are solved for by the optimization problem. Parameters are known quantities which are fixed before a solution is attempted.

| **Variables (solved for by optimization)** | | |
| --- | --- | --- |
| Symbol | Variable Type | Definition |
| $v_{model}(j,t)$ | Continuous | Modeled flux rate of reaction $j$ in condition or timepoint $t$. |
| $v_{h}\left( j,t \right)$ | Continuous | The flux found by optimization through kinetic equation type $h$. |
| $K_{1,2,3,4,5,6}(j)$ | Continuous positive | Kinetic (fitting) parameters for the optimal equation form. |
| $b_{1,2,3}(j)$ | Binary | Solved for optimal type of regulation |
| $\vec{\Omega}_{j}^{T}$ | Binary | $\Omega_{ij}^{T}=1$ if $i$ is optimal inhibitor of reaction $j$, $\Omega_{ij}^{T}=0$ otherwise |
| $\vec{\Gamma}_{j}^{T}$ | Binary | $\Gamma_{ij}^{T}=1$ if $i$ is optimal activator for reaction j, $\Gamma_{ij}^{T}=0$ otherwise |
| **Parameters (set by ‘omics and/or GSM)** | | |
| Symbol | Parameter Type | Definition |
| $I$ | Set | Set of metabolites in the GSM |
| $J$ | Set | Set of reactions in the GSM |
| $T$ | Set | Set of tested conditions or time points |
| $v_{\exp}(j,t)$ | Continuous | Experimental or FBA flux rate through reaction $j$ in condition or timepoint $t$ |
| $\beta_{1,2,3,4}(j)$ | Binary | Stores overarching type of reaction which may be determined by the GSM |
| $\vec{C}_{t}$ | Continuous | Experimental concentration or FBA metabolite pool size of in condition or timepoint $t$, stored as a 1-D array with dimension of set $i$ |
| $\vec{A}$ | Binary | Binary array indicating substrate A, with a single entry. $A_{i}=1$ if it is the first entry in $S_{ij}$ which is positive, $A_{i}=0$ otherwise. |
| $\vec{B}$ | Binary | Binary array indicating substrate B, with a single entry. $B_{i}=1$ if it is the second entry in $S_{ij}$ which is positive, $B_{i}=0$ otherwise. |
| $\vec{\vec{P}}$ | Binary | A version of the $S_{ij}$ matrix where negative entries are replaced by zero and positive entries are replaced by a one. Products matrix. |
| $\epsilon$ | Fixed | An arbitrary very small number. |

The current formulation of KOPTIC is given in the table below, with both conceptual and mathematical formulations.

**Table 3:** Preliminary formulation of KOPTIC used to obtain the reported preliminary KOPTIC results. See table 2 for definitions of symbols used in the equations here presented. The first column is the mathematical formulation of the equations, the second column is the equation number, and the third column is a brief conceptual description of the equation.

| **Mathematical Formulation (Preliminary)** | **Eq.** | **Conceptual Formulation** |
| --- | --- | --- |
| $minimize z=\sum_{t\in T} \left\{ \left( v_{\exp}\left( j,t \right)-v_{model}\left( j,t \right) \right)^{2}+\epsilon\sum_{m=1}^{6} K_{m}\left( j \right) \right\}$  $\forall j\in J$ | (1) | For each reaction:  Minimize sum of squared difference of experimental (or FBA) and fit flux rate |
| $s.t.$ |  | Subject to: |
| $v_{\mathrm{model}}\left( j,t \right)=\beta_{1,j}\left[ b_{1,j}\left( v_{\mathrm{SIN}}\left( j,t \right) \right)+b_{2,j}\left( v_{\mathrm{SII}}\left( j,t \right) \right)+b_{3,j}\left( v_{\mathrm{SIA}}\left( j,t \right) \right) \right]$  $+\beta_{2,j}\left[ b_{1,j}\left( v_{\mathrm{SRN}}\left( j,t \right) \right)+b_{2,j}\left( v_{\mathrm{SRI}}\left( j,t \right) \right)+b_{3,j}\left( v_{\mathrm{SRA}}\left( j,t \right) \right) \right]$  $+\beta_{3,j}\left[ b_{1,j}\left( v_{\mathrm{DIN}}\left( j,t \right) \right)+b_{2,j}\left( v_{\mathrm{DII}}\left( j,t \right) \right)+b_{3,j}\left( v_{\mathrm{DIA}}\left( j,t \right) \right) \right]$  $+\beta_{4,j}\left[ b_{1,j}\left( v_{\mathrm{DRN}}\left( j,t \right) \right)+b_{2,j}\left( v_{\mathrm{DRI}}\left( j,t \right) \right)+b_{3,j}\left( v_{\mathrm{DRA}}\left( j,t \right) \right) \right]$ | (2) | Select the best of the below kinetic equation forms to describe the experimental results. |
| $\sum_{m=1}^{4} \beta_{m}(j)\leq1,\forall\beta\in\left\{ 0,1 \right\}, \forall j\in J$ | (3) | Must have either 1 or 2 substrates, be reversible or irreversible or not be fit |
| $\sum_{n=1}^{3} b_{n}(j)\leq1, \forall b\in\left\{ 0,1 \right\},\forall j\in J$ | (4) | Select at most one equation as best describing the experimental data (no fit is found) |
| $\sum_{i} \Gamma_{ij}^{T}\leq1, \forall j\in J$ | (5) | At most one optimal activator |
| $\sum_{i} \Omega_{ij}^{T}\leq1, \forall j\in J$ | (6) | At most one optimal inhibitor |
| $v_{\mathrm{SIN}}(t,j)=\frac{K_{1}\left( \vec{A}^{T}\cdot\vec{C}_{t} \right)}{\left( \vec{A}^{T}\cdot\vec{C}_{t} \right)+K_{2}+\epsilon}$ | (7) | Single-substrate irreversible non-regulated (SIN) kinetics |
| $v_{\mathrm{SII}}(t,j)=\frac{K_{1}\left( \vec{A}^{T}\cdot\vec{C}_{t} \right)}{\left( \vec{A}^{T}\cdot\vec{C}_{t} \right)+K_{2}\left( 1+\frac{\vec{\Omega}_{j}^{T}\cdot\vec{C}_{t}}{K_{3}+\epsilon} \right)+\epsilon}$ | (8) | Single-substrate irreversible inhibited (SII) kinetics |
| $v_{\mathrm{SIA}}(t,j)=\frac{K_{1}\left( \vec{A}^{T}\cdot\vec{C}_{t} \right)\left( \vec{\Gamma}_{j}^{T}\cdot\vec{C}_{t} \right)}{1+K_{2}\left( \vec{\Gamma}_{j}^{T}\cdot\vec{C}_{t} \right)+K_{3}\left( \vec{A}^{T}\cdot\vec{C}_{t} \right)\left( \vec{\Gamma}_{j}^{T}\cdot\vec{C}_{t} \right)}$ | (9) | Single-substrate irreversible activated (SIA) kinetics |
| $v_{\mathrm{SRN}}(t,j)=v=\frac{K_{1}\left[ A \right]-K_{2}\left[ P_{1} \right]\ldots\left[ P_{N} \right]}{1+K_{3}\left[ A \right]+K_{4}\left[ P_{1} \right]\ldots\left[ P_{N} \right]}$ | (10) | Single-substrate reversible non-regulated (SRN) kinetics |
| $v_{\mathrm{SRI}}(t,j)=\frac{K_{1}\left( \vec{A}^{T}\cdot\vec{C}_{t} \right)-K_{2}\left[ \left( \vec{C}_{t}\cdot\vec{\vec{P}} \right)\cdot\vec{1} \right]}{1+K_{3}\left( \vec{\Omega}_{j}^{T}\cdot\vec{C}_{t} \right)+K_{4}\left( \vec{A}^{T}\cdot\vec{C}_{t} \right)+K_{5}\left[ \left( \vec{C}_{t}\cdot\vec{\vec{P}} \right)\cdot\vec{1} \right]}$ | (11) | Single-substrate reversible inhibited (SRI) kinetics |
| $v_{\mathrm{SRA}}\left( t,j \right)=$  $\frac{K_{1}\left( \vec{A}^{T}\cdot\vec{C}_{t} \right)\left( \vec{\Gamma}^{T}\cdot\vec{C}_{t} \right)-K_{2}\left[ \left( \vec{C}_{t}\cdot\vec{\vec{P}} \right)\cdot\vec{1} \right]\left( \vec{\Gamma}_{j}^{T}\cdot\vec{C}_{t} \right)}{1+K_{3}\vec{\Gamma}_{j}^{T}\cdot\vec{C}_{t}+K_{4}\left( \vec{A}^{T}\cdot\vec{C}_{t} \right)\left( \vec{\Gamma}_{j}^{T}\cdot\vec{C}_{t} \right)+K_{5}\left[ \left( \vec{C}_{t}\cdot\vec{\vec{P}} \right)\cdot\vec{1} \right]\left( \vec{\Gamma}_{j}^{T}\cdot\vec{C}_{t} \right)}$ | (12) | Single-substrate reversible activated (SRA) kinetics |
| $v_{\mathrm{DIN}}\left( t,j \right)=$  $\frac{K_{1}}{\left( \frac{K_{2}}{\left( \vec{A}^{T}\cdot\vec{C}_{t} \right)\left( \vec{B}^{T}\cdot\vec{C}_{t} \right)+\epsilon}+\frac{K_{3}}{\left( \vec{B}^{T}\cdot\vec{C}_{t} \right)+\epsilon}+\frac{K_{4}}{\left( \vec{A}^{T}\cdot\vec{C}_{t} \right)+\epsilon}+1 \right)}$ | (13) | Dual-substrate irreversible non-regulated (DIN) kinetic |
| $v_{\mathrm{DII}}\left( t,j \right)=$  $\frac{K_{1}\left( \vec{A}^{T}\cdot\vec{C}_{t} \right)\left( \vec{B}^{T}\cdot\vec{C}_{t} \right)}{1+K_{2}\left( \vec{A}^{T}\cdot\vec{C}_{t} \right)+K_{3}\left( \vec{B}^{T}\cdot\vec{C}_{t} \right)+K_{4}\left( \vec{\Omega}_{j}^{T}\cdot\vec{C}_{t} \right)+K_{5}\left( \vec{A}^{T}\cdot\vec{C}_{t} \right)\left( \vec{B}^{T}\cdot\vec{C}_{t} \right)}$ | (14) | Dual-substrate irreversible inhibited (DII) kinetics |
| $v_{\mathrm{DIA}}\left( t,j \right)=$  $\frac{K_{1}\left( \vec{\Gamma}_{j}^{T}\vec{C}_{t} \right)\left( \vec{B}^{T}\vec{C}_{t} \right)\left( \vec{A}^{T}\vec{C}_{t} \right)}{1+K_{2}\left( \vec{\Gamma}_{j}^{T}\vec{C}_{t} \right)\left( 1+K_{3}\left( \vec{A}^{T}\vec{C}_{t} \right)+K_{4}\left( \vec{B}^{T}\vec{C}_{t} \right)+K_{5}\left( \vec{A}^{T}\vec{C}_{t} \right)\left( \vec{B}^{T}\vec{C}_{t} \right) \right)}$ | (15) | Dual-substrate irreversible activated (DIA) kinetics |
| $v_{\mathrm{DRN}}(t,j)=\frac{K_{1}\left( \vec{A}^{T}\cdot\vec{C}_{t} \right)\left( \vec{B}^{T}\cdot\vec{C}_{t} \right)-K_{2}\left[ \left( \vec{C}_{t}\cdot\vec{\vec{P}} \right)\cdot\vec{1} \right]}{1+K_{3}\left( \vec{A}^{T}\cdot\vec{C}_{t} \right)+K_{4}\left( \vec{B}^{T}\cdot\vec{C}_{t} \right)+K_{5}\left( \vec{A}^{T}\cdot\vec{C}_{t} \right)\left( \vec{B}^{T}\cdot\vec{C}_{t} \right)}$ | (16) | Dual-substrate reversible non-regulated (DRN) kinetics |
| $v_{\mathrm{DRI}}(t,j)=$  $\frac{K_{1}\left( \vec{A}^{T}\cdot\vec{C}_{t} \right)\left( \vec{B}^{T}\cdot\vec{C}_{t} \right)-K_{2}\left[ \left( \vec{C}_{t}\cdot\vec{\vec{P}} \right)\cdot\vec{1} \right]}{1+K_{3}\left( \vec{A}^{T}\vec{C}_{t} \right)+K_{4}\left( \vec{\Omega}_{j}^{T}\cdot\vec{C}_{t} \right)+K_{5}\left( \vec{B}^{T}\cdot\vec{C}_{t} \right)+K_{6}\left( \vec{A}^{T}\cdot\vec{C}_{t} \right)\left( \vec{B}^{T}\cdot\vec{C}_{t} \right)}$ | (17) | Dual-substrate reversible inhibited (DRI) kinetics |
| $v_{\mathrm{DRA}}\left( t,j \right)=$  $\frac{K_{1}\left( \vec{\Gamma}_{j}^{T}\cdot\vec{C} \right)\left( \vec{B}^{T}\cdot\vec{C} \right)\left( \vec{A}^{T}\cdot\vec{C} \right)-K_{2}\left( \vec{\Gamma}_{j}^{T}\cdot\vec{C} \right)\left[ \left( \vec{C}_{t}\cdot\vec{\vec{P}} \right)\cdot\vec{1} \right]}{1+K_{3}\left( \vec{\Gamma}_{j}^{T}\cdot\vec{C}_{t} \right)\left( 1+K_{4}\left( \vec{A}^{T}\cdot\vec{C}_{t} \right)+K_{5}\left( \vec{B}^{T}\cdot\vec{C}_{t} \right)+K_{6}\left( \vec{B}^{T}\cdot\vec{C}_{t} \right)\left( \vec{A}^{T}\cdot\vec{C}_{t} \right) \right)}$ | (18) | Dual-substrate reversible activated (DRA) kinetics |
| $v_{TIN}=\frac{K_{1}\left( \vec{A}^{T}\cdot\vec{C} \right)\left( \vec{B}^{T}\cdot\vec{C} \right)\left( {\vec{\mathrm{subC}}}^{T}\cdot\vec{C} \right)}{1+K_{2}\left( \vec{A}^{T}\cdot\vec{C} \right)+K_{3}\left( \vec{B}^{T}\cdot\vec{C} \right)+K_{4}\left( {\vec{\mathrm{subC}}}^{T}\cdot\vec{C} \right)+K_{5}\left( \vec{A}^{T}\cdot\vec{C} \right)\left( \vec{B}^{T}\cdot\vec{C} \right)+K_{6}\left( \vec{A}^{T}\cdot\vec{C} \right)\left( {\vec{\mathrm{subC}}}^{T}\cdot\vec{C} \right)+K_{7}\left( \vec{B}^{T}\cdot\vec{C} \right)\left( {\vec{\mathrm{subC}}}^{T}\cdot\vec{C} \right)+K_{8}\left( \vec{A}^{T}\cdot\vec{C} \right)\left( \vec{B}^{T}\cdot\vec{C} \right)\left( {\vec{\mathrm{subC}}}^{T}\cdot\vec{C} \right)}$ | (19) | Triple-substrate irreversible non-regulated (TIN) kinetics |
| $v_{TII}=\frac{K_{1}\left( \vec{A}^{T}\cdot\vec{C} \right)\left( \vec{B}^{T}\cdot\vec{C} \right)\left( {\vec{\mathrm{subC}}}^{T}\cdot\vec{C} \right)}{1+K_{2}\left( \vec{A}^{T}\cdot\vec{C} \right)+K_{3}\left( \vec{B}^{T}\cdot\vec{C} \right)+K_{4}\left( {\vec{\mathrm{subC}}}^{T}\cdot\vec{C} \right)+K_{5}\left( \vec{\Omega}_{j}^{T}\cdot\vec{C}_{t} \right)+K_{6}\left( \vec{A}^{T}\cdot\vec{C} \right)\left( \vec{B}^{T}\cdot\vec{C} \right)+K_{7}\left( \vec{A}^{T}\cdot\vec{C} \right)\left[ C \right]+K_{8}\left( \vec{B}^{T}\cdot\vec{C} \right)\left( {\vec{\mathrm{subC}}}^{T}\cdot\vec{C} \right)+K_{9}\left( \vec{A}^{T}\cdot\vec{C} \right)\left( \vec{B}^{T}\cdot\vec{C} \right)\left( {\vec{\mathrm{subC}}}^{T}\cdot\vec{C} \right)}$ | (20) | Triple-substrate irreversible inhibited (TII) kinetics |
| $v_{TIA}=$  $\frac{K_{1}\left( \vec{\Gamma}^{T}\cdot\vec{C} \right)\left( \vec{A}^{T}\cdot\vec{C} \right)\left( \vec{B}^{T}\cdot\vec{C} \right)\left( {\vec{\mathrm{subC}}}^{T}\cdot\vec{C} \right)}{1+K_{2}\left[ \alpha\right]+K_{3}\left( \vec{\Gamma}^{T}\cdot\vec{C} \right)\left( \vec{A}^{T}\cdot\vec{C} \right)+K_{4}\left[ \alpha\right]\left( \vec{B}^{T}\cdot\vec{C} \right)+K_{5}\left( \vec{\Gamma}^{T}\cdot\vec{C} \right)\left( {\vec{\mathrm{subC}}}^{T}\cdot\vec{C} \right)+K_{6}\left( \vec{\Gamma}^{T}\cdot\vec{C} \right)\left( \vec{A}^{T}\cdot\vec{C} \right)\left( \vec{B}^{T}\cdot\vec{C} \right)+K_{7}\left( \vec{\Gamma}^{T}\cdot\vec{C} \right)\left( \vec{A}^{T}\cdot\vec{C} \right)\left( {\vec{\mathrm{subC}}}^{T}\cdot\vec{C} \right)+K_{8}\left( \vec{\Gamma}^{T}\cdot\vec{C} \right)\left( \vec{B}^{T}\cdot\vec{C} \right)\left( {\vec{\mathrm{subC}}}^{T}\cdot\vec{C} \right)+K_{9}\left( \vec{\Gamma}^{T}\cdot\vec{C} \right)\left( \vec{A}^{T}\cdot\vec{C} \right)\left( \vec{B}^{T}\cdot\vec{C} \right)\left( {\vec{\mathrm{subC}}}^{T}\cdot\vec{C} \right)}$ | (21) | Triple-substrate irreversible activated (TIA) kinetics |
| $v_{TRN}=\frac{K_{1}\left( \vec{A}^{T}\cdot\vec{C} \right)\left( \vec{B}^{T}\cdot\vec{C} \right)\left( {\vec{\mathrm{subC}}}^{T}\cdot\vec{C} \right)-K_{2}\left[ \left( \vec{C}_{t}\cdot\vec{\vec{P}} \right)\cdot\vec{1} \right]}{1+K_{3}\left( \vec{A}^{T}\cdot\vec{C} \right)+K_{4}\left( \vec{B}^{T}\cdot\vec{C} \right)+K_{5}\left( {\vec{\mathrm{subC}}}^{T}\cdot\vec{C} \right)+K_{6}\left( \vec{A}^{T}\cdot\vec{C} \right)\left( \vec{B}^{T}\cdot\vec{C} \right)+K_{7}\left( \vec{A}^{T}\cdot\vec{C} \right)\left( {\vec{\mathrm{subC}}}^{T}\cdot\vec{C} \right)+K_{8}\left( \vec{B}^{T}\cdot\vec{C} \right)\left( {\vec{\mathrm{subC}}}^{T}\cdot\vec{C} \right)+K_{9}\left( \vec{A}^{T}\cdot\vec{C} \right)\left( \vec{B}^{T}\cdot\vec{C} \right)\left( {\vec{\mathrm{subC}}}^{T}\cdot\vec{C} \right)}$ | (22) | Triple-substrate reversible non-regulated (TRN) kinetics |
| $v_{TRI}=\frac{K_{1}\left( \vec{A}^{T}\cdot\vec{C} \right)\left( \vec{B}^{T}\cdot\vec{C} \right)\left( {\vec{\mathrm{subC}}}^{T}\cdot\vec{C} \right)-K_{2}\left[ \left( \vec{C}_{t}\cdot\vec{\vec{P}} \right)\cdot\vec{1} \right]}{1+K_{3}\left( \vec{A}^{T}\cdot\vec{C} \right)+K_{4}\left( \vec{B}^{T}\cdot\vec{C} \right)+K_{5}\left( {\vec{\mathrm{subC}}}^{T}\cdot\vec{C} \right)+K_{6}\left( \vec{\Omega}_{j}^{T}\cdot\vec{C}_{t} \right)+K_{7}\left( \vec{A}^{T}\cdot\vec{C} \right)\left( \vec{B}^{T}\cdot\vec{C} \right)+K_{8}\left( \vec{A}^{T}\cdot\vec{C} \right)\left( {\vec{\mathrm{subC}}}^{T}\cdot\vec{C} \right)+K_{9}\left( \vec{B}^{T}\cdot\vec{C} \right)\left[ C \right]+K_{10}\left( \vec{A}^{T}\cdot\vec{C} \right)\left( \vec{B}^{T}\cdot\vec{C} \right)\left( {\vec{\mathrm{subC}}}^{T}\cdot\vec{C} \right)}$ | (23) | Triple-substrate reversible inhibited (TRI) kinetics |
| $v_{TRA}=$  $\frac{K_{1}\left( \vec{\Gamma}^{T}\cdot\vec{C} \right)\left( \vec{A}^{T}\cdot\vec{C} \right)\left( \vec{B}^{T}\cdot\vec{C} \right)\left( {\vec{\mathrm{subC}}}^{T}\cdot\vec{C} \right)-K_{2}\left( \vec{\Gamma}^{T}\cdot\vec{C} \right)\left[ \left( \vec{C}_{t}\cdot\vec{\vec{P}} \right)\cdot\vec{1} \right]}{1+K_{3}\left( \vec{\Gamma}^{T}\cdot\vec{C} \right)+K_{4}\left( \vec{\Gamma}^{T}\cdot\vec{C} \right)\left( \vec{A}^{T}\cdot\vec{C} \right)+K_{5}\left[ \alpha\right]\left( \vec{B}^{T}\cdot\vec{C} \right)+K_{6}\left( \vec{\Gamma}^{T}\cdot\vec{C} \right)\left( {\vec{\mathrm{subC}}}^{T}\cdot\vec{C} \right)+K_{7}\left( \vec{\Gamma}^{T}\cdot\vec{C} \right)\left( \vec{A}^{T}\cdot\vec{C} \right)\left( \vec{B}^{T}\cdot\vec{C} \right)+K_{8}\left( \vec{\Gamma}^{T}\cdot\vec{C} \right)\left( \vec{A}^{T}\cdot\vec{C} \right)\left( {\vec{\mathrm{subC}}}^{T}\cdot\vec{C} \right)+K_{9}\left( \vec{\Gamma}^{T}\cdot\vec{C} \right)\left( \vec{B}^{T}\cdot\vec{C} \right)\left( {\vec{\mathrm{subC}}}^{T}\cdot\vec{C} \right)+K_{10}\left( \vec{\Gamma}^{T}\cdot\vec{C} \right)\left( \vec{A}^{T}\cdot\vec{C} \right)\left( \vec{B}^{T}\cdot\vec{C} \right)\left( {\vec{\mathrm{subC}}}^{T}\cdot\vec{C} \right)}$ | (24) | Triple-substrate reversible activated (TRA) kinetics |
| $0\leq K_{m}\left( j \right)\leq10000, m\in\left[ 1,2,3,4,5,6 \right],\forall j\in J$ | (25) | Given kinetic parameter bounds |
| $\left\vert v_{h}\left( t,j \right) \right\vert\leq10000, h\in\left[ SRN,SRI,SRA,DRN,DRI,DRA \right],\forall t\in T,$  $\forall j\in J$ | (26) | Given reaction rates bounds |
| $0\leq v_{h}\left( t,j \right)\leq10000, h\in\left[ SIN,SII,SIA,DIN,DII,DIA \right], \forall t\in T,$  $\forall j\in J$ | (27) | Given reaction rates bounds |
| $K_{1}\left( j \right)\geq\epsilon, \forall j\in J$ | (28) | Ensures $K_{1}(j)$ is strictly positive |
| $K_{2}\left( j \right)\geq\epsilon, \forall j\in J$ | (29) | Ensures $K_{2}(j)$ is strictly positive |
| $K_{3}\left( j \right)\geq\left( 1-\beta_{1}\left( j \right)*b_{1}\left( j \right) \right)*2*\epsilon-\epsilon, \forall j\in J$ | (30) | Ensure $K_{3}(j)$ is strictly positive when required for kinetic equation. |
| $K_{3}\left( j \right)=\left( 1-\beta_{1}\left( j \right)*b_{1}\left( j \right) \right)K_{3}\left( j \right), \forall j\in J$ | (31) | Ensure $K_{3}(j)$ is zero when not required for kinetic equation. |
| $K_{4}\left( j \right)\geq\left( 1-\beta_{1}\left( j \right) \right)*2*\epsilon-\epsilon, \forall j\in J$ | (32) | Ensure $K_{4}(j)$ is strictly positive when required for kinetic equation. |
| $K_{4}\left( j \right)=\left( 1-\beta_{1}\left( j \right) \right)K_{4}\left( j \right), \forall j\in J$ | (33) | Ensure $K_{4}(j)$ is zero when not required for kinetic equation. |
| $K_{5}\left( j \right)\geq2\epsilon\beta_{2}\left( j \right)*\left( 1-b_{1}\left( j \right) \right)+2\epsilon\beta_{3}\left( j \right)*\left( 1-b_{1}\left( j \right) \right)+2\epsilon\beta_{4}-\epsilon$  $\forall j\in J$ | (34) | Ensure $K_{5}(j)$ is strictly positive when required for kinetic equation. |
| $K_{5}\left( j \right)=\beta_{2}\left( j \right)K_{4}\left( j \right)*\left( 1-b_{1}\left( j \right) \right)+\beta_{3}\left( j \right)K_{4}\left( j \right)*\left( 1-b_{1}\left( j \right) \right)$  $+\beta_{4}\left( j \right)K_{4}\left( j \right)$  $\forall j\in J$ | (35) | Ensure $K_{5}(j)$ is zero when not required for kinetic equation. |
| $K_{6}\left( j \right)\geq2\left( \beta_{4}\left( j \right)\left( 1-b_{1}\left( j \right) \right)+\beta_{5}+\beta_{6} \right)\epsilon-\epsilon,\forall j\in J$ | (36) | Ensure $K_{6}(j)$ is strictly positive when required for kinetic equation. |
| $K_{6}\left( j \right)=K_{6}\left( j \right)(\beta_{4}\left( j \right)\left( 1-b_{1}\left( j \right) \right)+\beta_{5}+\beta_{6}),\forall j\in J$ | (37) | Ensure $K_{6}(j)$ is zero when not required for kinetic equation. |
| $K_{7}=\left( \beta_{5}+\beta_{6} \right)K_{7}$ | (38) |  |
| $K_{7}\geq2\left( \beta_{5}+\beta_{6} \right)\epsilon-\epsilon$ | (39) |  |
| $K_{8}=\left( \beta_{5}+\beta_{6} \right)K_{8}$ | (40) |  |
| $K_{8}\geq2\left( \beta_{5}+\beta_{6} \right)\epsilon-\epsilon$ | (41) |  |
| $K_{9}=\left( \beta_{5}\left( j \right)\left( 1-b_{1}\left( j \right) \right)+\beta_{6} \right)K_{9}$ | (42) |  |
| $K_{9}\geq2\left( \beta_{5}\left( j \right)\left( 1-b_{1}\left( j \right) \right)+\beta_{6} \right)\epsilon-\epsilon$ | (43) |  |
| $K_{10}=\left( \beta_{6}\left( 1-b_{1}\left( j \right) \right) \right)K_{10}$ | (44) |  |
| $K_{10}\geq2\left( \beta_{6}\left( 1-b_{1}\left( j \right) \right) \right)\epsilon-\epsilon$ | (45) |  |

In the immediately preceding table 3, equation (1) is the objective function, equation (2) is the modeled kinetic flux rate subject to the different kinetic equation forms given in equations (7) through (18). Equations (3) through (6) guarantee a single optimal kinetic equation form and a single optimal regulator. The bounds of the problem are set in equations (19) through (21). Equations (22) through (24), (26), (28), and (31) ensure that a non-zero optimal value is found for all kinetic parameters used in the optimal kinetic equation form by simplifying to $K_{m}\left( j \right)\geq\epsilon$ when used and to $K_{m}\left( j \right)\geq-\epsilon$ when unused. Equations (25), (27), (29), and (31) are used to ensure that kinetic parameters are fixed at zero that are not used in the optimal kinetic formulation by simplifying to $K_{m}\left( j \right)=0$ when unused and $K_{m}\left( j \right)=K_{m}\left( j \right)$ when used. The formulation above is looped for each reaction, solving for the regulation and kinetics of one reaction at a time and fixes all variables not related to the current reaction being examined as both a time saving measure and a measure to decrease computational intensity.

#### **1.2 Advantages**

The concept of KOPTIC and its current formulation has several advantages over traditional kinetic modeling methods.

1. No *a priori* knowledge of regulatory mechanisms in the system is required in order to build a kinetic model that uses regulatory mechanisms. The KOPTIC method is therefore the best choice at present for building kinetic models of metabolism for under-studied systems.
2. Further, experimental flux data is not necessary, FBA results can be easily used by KOPTIC for an *in silico* approach to develop kinetic models of metabolism.
3. Less manual labor is required to create a kMM, this method does not require the writing of kinetic mechanisms or elementary rate laws for each reaction. Once KOPTIC is implemented in some platform for the system, the computer takes over the process, freeing up the researcher to complete other tasks.
4. The formulation is modular, and allows users to substitute their own kinetic equations once they are converted to the correct form to fit KOPTIC.
5. This method makes use of stoichiometric models of metabolism, of which there are many published and which is a common starting point for computational approaches to metabolism.
6. This work demonstrates that KOPTIC is readily applicable to multi-tissue systems for use in creating kinetic models of metabolism for whole systems.
7. Since KOPTIC works on a per-reaction basis, KOPTIC is parallelizable. Several parallel instances of KOPTIC can to develop the kinetic model of metabolism much faster than a single instance.

#### **1.3 Limitation**

This current formulation has several limitations, which this research group intends to address in future.

1. Each metabolic inhibitor is equally likely unless it is specifically prohibited by the user. This is clearly not biologically true because some metabolites are more likely to be regulators than others.
2. There is only at present twelve equations forms, each of which allows only a single activator or inhibitor. This limitation can be overcome by deriving and adding/substituting new equation types, or by using integer cuts (and other limitations) to get several possible regulation schemes and the most likely scheme (as judged by the user) can be selected.
3. Reactions with more than two substrates cannot be fit by the current formulation.
4. Enzyme regulation of other enzymes is very common in biological systems, but the current formulation cannot account for this.

### **2 Kinetics Formulations and Derivations**

In the following formulations the notation that is used is:

$A$ and $B$ are substrates

$\alpha$ is an activator (another enzyme or substrate)

$I$ is an inhibitor (another enzyme or substrate)

$E$ is an enzyme

$E_{t}$ is total enzyme concentration (free enzyme+all enzyme complexes)

$V_{max}$ or $V_{max,f}$ is the maximum flux rate (forward) from FVA

$V_{max,b}$ is the maximum reverse (backwards) flux rate from FVA (only applies to reversible reactions)

As per convention an upper case $K$ is the final (bolded) kinetics equation represents an empirical reaction kinetics parameters. As also per convention, during the derivation a lower case $k$ represents a kinetic rate parameter (or some combination of these), where an upper case $K$ represents an equilibrium constant.

#### **2.1 Irreversible Single Substrate Kinetics**

If $\boldsymbol{\beta}_{\boldsymbol{1,j}}\boldsymbol{=1}$ then irreversible single substrate irreversible reaction kinetics. Derivation or statement of the equations related to single substrate irreversible enzyme kinetics.

**2.1.1 Irreversible Michaelis-Menten Kinetics**

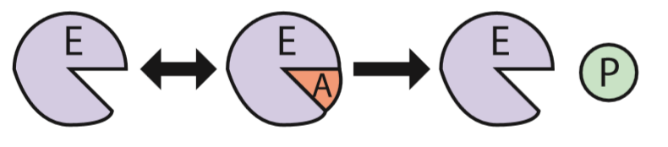
The irreversible Michaelis-Menten Kinetics elementary steps is:

$$A+E\rightleftharpoons EA$$

$$EA\to P+E$$

Giving the kinetics equation of (from *Elements of Chemical Reaction Engineering* by Fogler)

$$\boldsymbol{v=}\frac{\boldsymbol{k}_{\boldsymbol{cat}}\left[ \boldsymbol{E}_{\boldsymbol{t}} \right]\boldsymbol{[A]}}{\left[ \boldsymbol{A} \right]\boldsymbol{+}\boldsymbol{K}_{\boldsymbol{M}}}$$

##### **2.1.2 Irreversible Single-Substrate Allosteric Activation Kinetics**

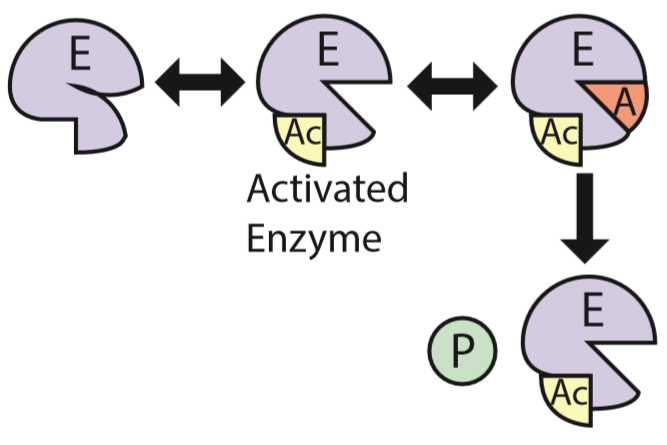
Allosteric activation of the enzyme. Allosteric activation required for substrate binding. The elementary steps are:

$$\alpha$$

$$E+\alpha\rightleftharpoons E\alpha,k_{1}\to,k_{1}^{'}\leftarrow$$

$$E\alpha+A\rightleftharpoons E\alpha A, k_{2}\to,k_{2}^{'}\leftarrow$$

$$E\alpha A\to E\alpha+P,k_{3}\to$$

This results in elementary step reaction rates of:

$$r_{E\alpha}=k_{1}\left[ E \right]\left[ \alpha\right]-k_{1}^{'}\left[ E\alpha\right]-k_{2}\left[ E\alpha\right]\left[ A \right]+k_{2}^{'}[E\alpha A]+k_{3}[E\alpha A]$$

$$r_{E\alpha A}=k_{2}\left[ E\alpha\right]\left[ A \right]-k_{2}^{'}\left[ E\alpha A \right]-k_{3}\left[ E\alpha A \right]$$

$$v=k_{3}\left[ E\alpha A \right]$$

Applying the Pseudo-Steady-State Hypothesis to enzyme complexes gives:

$$r_{E\alpha}=0=k_{1}\left[ E \right]\left[ \alpha\right]-k_{1}^{'}\left[ E\alpha\right]-k_{2}\left[ E\alpha\right]\left[ A \right]+k_{2}^{'}[E\alpha A]+k_{3}[E\alpha A]$$

$$0=k_{1}\left[ E \right]\left[ \alpha\right]-\left( k_{1}^{'}+k_{2}\left[ A \right] \right)\left[ E\alpha\right]+\left( k_{2}^{'}+k_{3} \right)[E\alpha A]$$

$$\left( k_{1}^{'}+k_{2}\left[ A \right] \right)\left[ E\alpha\right]=k_{1}\left[ E \right]\left[ \alpha\right]+\left( k_{2}^{'}+k_{3} \right)[E\alpha A]$$

$$\left[ E\alpha\right]=\frac{k_{1}\left[ E \right]\left[ \alpha\right]}{k_{1}^{'}+k_{2}\left[ A \right]}+\frac{k_{2}^{'}+k_{3}}{k_{1}^{'}+k_{2}\left[ A \right]}[E\alpha A]$$

$$r_{E\alpha A}=0=k_{2}\left[ E\alpha\right]\left[ A \right]-k_{2}^{'}\left[ E\alpha A \right]-k_{3}\left[ E\alpha A \right]$$

$$0=k_{2}\left[ E\alpha\right]\left[ A \right]-\left( k_{2}^{'}+k_{3} \right)\left[ E\alpha A \right]$$

$$\left( k_{2}^{'}+k_{3} \right)\left[ E\alpha A \right]=k_{2}\left[ E\alpha\right]\left[ A \right]$$

$$\left[ E\alpha A \right]=\frac{k_{2}\left[ A \right]}{k_{2}^{'}+k_{3}}\left[ E\alpha\right]$$

$$\left[ E\alpha\right]=\frac{k_{1}\left[ E \right]\left[ \alpha\right]}{k_{1}^{'}+k_{2}\left[ A \right]}+\frac{k_{2}^{'}+k_{3}}{k_{1}^{'}+k_{2}\left[ A \right]}\frac{k_{2}\left[ A \right]}{k_{2}^{'}+k_{3}}\left[ E\alpha\right]$$

$$\left[ E\alpha\right]=\frac{k_{1}\left[ E \right]\left[ \alpha\right]}{k_{1}^{'}+k_{2}\left[ A \right]}+\frac{k_{2}\left[ A \right]}{k_{1}^{'}+k_{2}\left[ A \right]}\left[ E\alpha\right]$$

$$\left( 1-\frac{k_{2}\left[ A \right]}{k_{1}^{'}+k_{2}\left[ A \right]} \right)\left[ E\alpha\right]=\frac{k_{1}\left[ E \right]\left[ \alpha\right]}{k_{1}^{'}+k_{2}\left[ A \right]}$$

$$\left( \frac{k_{1}^{'}+k_{2}\left[ A \right]-k_{2}\left[ A \right]}{k_{1}^{'}+k_{2}\left[ A \right]} \right)\left[ E\alpha\right]=\frac{k_{1}\left[ E \right]\left[ \alpha\right]}{k_{1}^{'}+k_{2}\left[ A \right]}$$

$$\left( \frac{k_{1}^{'}}{k_{1}^{'}+k_{2}\left[ A \right]} \right)\left[ E\alpha\right]=\frac{k_{1}\left[ E \right]\left[ \alpha\right]}{k_{1}^{'}+k_{2}\left[ A \right]}$$

$$\left[ E\alpha\right]=\frac{k_{1}\left[ E \right]\left[ \alpha\right]}{k_{1}^{'}+k_{2}\left[ A \right]}*\frac{k_{1}^{'}+k_{2}\left[ A \right]}{k_{1}^{'}}=\frac{k_{1}}{k_{1}^{'}}\left[ E \right]\left[ \alpha\right]$$

$$\left[ E\alpha A \right]=\frac{k_{2}\left[ A \right]}{k_{2}^{'}+k_{3}}\left[ E\alpha\right]=\frac{k_{2}\left[ A \right]}{k_{2}^{'}+k_{3}}\frac{k_{1}}{k_{1}^{'}}\left[ E \right]\left[ \alpha\right]$$

$$\left[ E\alpha A \right]=\frac{k_{1}k_{2}}{k_{1}^{'}\left( k_{2}^{'}+k_{3} \right)}\left[ A \right]\left[ E \right]\left[ \alpha\right]$$

$$v=k_{3}\left[ E\alpha A \right]=\frac{k_{1}k_{2}k_{3}}{k_{1}^{'}\left( k_{2}^{'}+k_{3} \right)}\left[ A \right]\left[ E \right]\left[ \alpha\right]$$

The total enzyme concentration is given as:

$$\left[ E_{t} \right]=\left[ E \right]+\left[ E\alpha\right]+[E\alpha A]$$

Making substitutions for the enzyme complexes in the above equation gives:

$$\left[ E_{t} \right]=\left[ E \right]+\frac{k_{1}}{k_{1}^{'}}\left[ E \right]\left[ \alpha\right]+\frac{k_{1}k_{2}}{k_{1}^{'}\left( k_{2}^{'}+k_{3} \right)}\left[ A \right]\left[ E \right]\left[ \alpha\right]$$

Solving for $[E]$ gives:

$$\left[ E_{t} \right]=\left[ E \right]\left( 1+\frac{k_{1}}{k_{1}^{'}}\left[ \alpha\right]+\frac{k_{1}k_{2}}{k_{1}^{'}\left( k_{2}^{'}+k_{3} \right)}\left[ A \right]\left[ \alpha\right] \right)$$

$$\left[ E_{t} \right]=\left[ E \right]\left( \frac{\left( k_{2}^{'}+k_{3} \right)}{\left( k_{2}^{'}+k_{3} \right)}+\frac{k_{1}\left( k_{2}^{'}+k_{3} \right)}{k_{1}^{'}\left( k_{2}^{'}+k_{3} \right)}\left[ \alpha\right]+\frac{k_{1}k_{2}}{k_{1}^{'}\left( k_{2}^{'}+k_{3} \right)}\left[ A \right]\left[ \alpha\right] \right)$$

$$\left[ E_{t} \right]=\left[ E \right]\left( \frac{\left( k_{2}^{'}+k_{3} \right)+k_{1}\left( k_{2}^{'}+k_{3} \right)\left[ \alpha\right]+k_{1}k_{2}\left[ A \right]\left[ \alpha\right]}{k_{1}^{'}\left( k_{2}^{'}+k_{3} \right)} \right)$$

$$\left[ E \right]=\frac{\left[ E_{t} \right]k_{1}^{'}\left( k_{2}^{'}+k_{3} \right)}{\left( k_{2}^{'}+k_{3} \right)+k_{1}\left( k_{2}^{'}+k_{3} \right)\left[ \alpha\right]+k_{1}k_{2}\left[ A \right]\left[ \alpha\right]}$$

Substituting into the reaction velocity equation gives:

$$v=\frac{k_{1}k_{2}k_{3}}{k_{1}^{'}\left( k_{2}^{'}+k_{3} \right)}\left[ A \right]\left[ E \right]\left[ \alpha\right]=\frac{k_{1}k_{2}k_{3}}{k_{1}^{'}\left( k_{2}^{'}+k_{3} \right)}\frac{\left[ E_{t} \right]k_{1}^{'}\left( k_{2}^{'}+k_{3} \right)}{\left( k_{2}^{'}+k_{3} \right)+k_{1}\left( k_{2}^{'}+k_{3} \right)\left[ \alpha\right]+k_{1}k_{2}\left[ A \right]\left[ \alpha\right]}\left[ A \right]\left[ \alpha\right]$$

$$v=\frac{k_{1}k_{2}k_{3}\left[ E_{t} \right]\left[ A \right]\left[ \alpha\right]}{\left( k_{2}^{'}+k_{3} \right)+k_{1}\left( k_{2}^{'}+k_{3} \right)\left[ \alpha\right]+k_{1}k_{2}\left[ A \right]\left[ \alpha\right]}$$

$$v=\frac{k_{1}k_{2}k_{3}\left[ E_{t} \right]\left[ A \right]\left[ \alpha\right]}{\left( k_{2}^{'}+k_{3} \right)\left( 1+k_{1}\left[ \alpha\right]+\frac{k_{1}k_{2}}{\left( k_{2}^{'}+k_{3} \right)}\left[ A \right]\left[ \alpha\right] \right)}$$

$$v=\frac{\frac{k_{1}k_{2}k_{3}}{\left( k_{2}^{'}+k_{3} \right)}\left[ E_{t} \right]\left[ A \right]\left[ \alpha\right]}{1+k_{1}\left[ \alpha\right]+\frac{k_{1}k_{2}}{\left( k_{2}^{'}+k_{3} \right)}\left[ A \right]\left[ \alpha\right]}$$

$$v=\frac{k_{cat}\left[ E_{t} \right]\left[ A \right]\left[ \alpha\right]}{1+K_{\alpha}\left[ \alpha\right]+K_{\alpha A}\left[ A \right]\left[ \alpha\right]}$$

Therefore the kinetic equation is:

$$\boldsymbol{v=}\frac{\boldsymbol{k}_{\boldsymbol{cat}}\left[ \boldsymbol{E}_{\boldsymbol{t}} \right]\left[ \boldsymbol{A} \right]\left[ \boldsymbol{\alpha} \right]}{\boldsymbol{1+}\boldsymbol{K}_{\boldsymbol{\alpha}}\left[ \boldsymbol{\alpha} \right]\boldsymbol{+}\boldsymbol{K}_{\boldsymbol{\alpha A}}\left[ \boldsymbol{A} \right]\left[ \boldsymbol{\alpha} \right]}\boldsymbol{=}\frac{\boldsymbol{V}_{\boldsymbol{max}}\left[ \boldsymbol{A} \right]\left[ \boldsymbol{\alpha} \right]}{\boldsymbol{1+}\boldsymbol{K}_{\boldsymbol{\alpha}}\left[ \boldsymbol{\alpha} \right]\boldsymbol{+}\boldsymbol{K}_{\boldsymbol{\alpha A}}\left[ \boldsymbol{A} \right]\left[ \boldsymbol{\alpha} \right]}$$

$$\boldsymbol{parameters:}\left[ \boldsymbol{\alpha} \right]\boldsymbol{,}\left[ \boldsymbol{A} \right]\boldsymbol{,}\boldsymbol{V}_{\boldsymbol{max}}$$

$$\boldsymbol{variables:}\boldsymbol{K}_{\boldsymbol{\alpha}}\boldsymbol{,}\boldsymbol{K}_{\boldsymbol{\alpha A}}$$

**2.1.3 Irreversible Single-Substrate Allosteric Inhibition Kinetics**

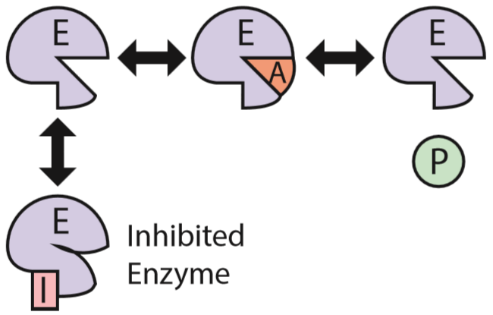
Mixed inhibition kinetics elementary steps are:

$$E+A\rightleftharpoons EA$$

$$E+I\rightleftharpoons EI$$

$$EA\to E+P$$

Giving the kinetics equation of (from *Elements of Chemical Reaction Engineering* by Fogler)

$$\boldsymbol{v=}\frac{\boldsymbol{k}_{\boldsymbol{cat}}\boldsymbol{[}\boldsymbol{E}_{\boldsymbol{t}}\boldsymbol{][A]}}{\left[ \boldsymbol{A} \right]\boldsymbol{+}\boldsymbol{K}_{\boldsymbol{M}}\left( \boldsymbol{1+}\frac{\left[ \boldsymbol{I} \right]}{\boldsymbol{K}_{\boldsymbol{I}}} \right)}\boldsymbol{=}\frac{\boldsymbol{V}_{\boldsymbol{max}}\boldsymbol{[A]}}{\left[ \boldsymbol{A} \right]\boldsymbol{+}\boldsymbol{K}_{\boldsymbol{M}}\left( \boldsymbol{1+}\frac{\left[ \boldsymbol{I} \right]}{\boldsymbol{K}_{\boldsymbol{I}}} \right)}$$

$$\boldsymbol{parameters:}\left[ \boldsymbol{A} \right]\boldsymbol{,}\boldsymbol{V}_{\boldsymbol{max}}\boldsymbol{,}\left[ \boldsymbol{I} \right]$$

$$\boldsymbol{variables:}\boldsymbol{K}_{\boldsymbol{M}}\boldsymbol{,}\boldsymbol{K}_{\boldsymbol{I}}$$

#### **2.2 Reversible Single Substrate Kinetics**

##### **2.2.1 Reversible Single-Substrate Michealis-Menten Kinetics**

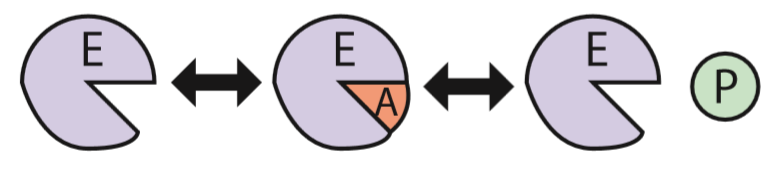
This kinetic scheme is largely the same as normal Michealis-Menten kinetics except the final product-forming step is now reversible (allowing for reversibility of enzyme function). The elementary steps for reversible Michealis-Menten kinetics is:

$$A+E\rightleftharpoons EA, k_{1}\to,k_{1}^{'}\leftarrow$$

$$EA\rightleftharpoons P+E, k_{2}\to,k_{2}^{'}\leftarrow$$

The reaction rate equations for enzyme complexes and reaction velocity:

$$r_{EA}=k_{1}\left[ A \right]\left[ E \right]-k_{1}^{'}\left[ EA \right]-k_{2}\left[ EA \right]+k_{2}^{'}\left[ E \right]\left[ P \right]$$

$$v=k_{2}\left[ EA \right]-k_{2}^{'}\left[ E \right]\left[ P \right]$$

Solving for $[EA]$ using the PSSH gives:

$$r_{EA}=0=k_{1}\left[ A \right]\left[ E \right]-\left( k_{1}^{'}+k_{2} \right)\left[ EA \right]+k_{2}^{'}\left[ E \right]\left[ P_{1} \right]\ldots\left[ P_{N} \right]$$

$$\left( k_{1}^{'}+k_{2} \right)\left[ EA \right]=k_{1}\left[ A \right]\left[ E \right]+k_{2}^{'}\left[ E \right]\left[ P_{1} \right]\ldots\left[ P_{N} \right]$$

$$\left[ EA \right]=\left( \frac{k_{1}\left[ A \right]}{k_{1}^{'}+k_{2}}+\frac{k_{2}^{'}\left[ P_{1} \right]\ldots\left[ P_{N} \right]}{k_{1}^{'}+k_{2}} \right)\left[ E \right]$$

$$\left[ EA \right]=\left( \frac{k_{1}\left[ A \right]+k_{2}^{'}\left[ P_{1} \right]\ldots\left[ P_{N} \right]}{k_{1}^{'}+k_{2}} \right)\left[ E \right]$$

Since determining the concentration of the non-complexed enzyme is difficult, use the total enzyme concentration and solving for non-complexed enzyme concentration:

$$\left[ E_{t} \right]=\left[ E \right]+[EA]$$

$$\left[ E_{t} \right]=\left[ E \right]+\left( \frac{k_{1}\left[ A \right]+k_{2}^{'}\left[ P_{1} \right]\ldots\left[ P_{N} \right]}{k_{1}^{'}+k_{2}} \right)\left[ E \right]$$

$$\left[ E_{t} \right]=\left( 1+\frac{k_{1}\left[ A \right]+k_{2}^{'}\left[ P_{1} \right]\ldots\left[ P_{N} \right]}{k_{1}^{'}+k_{2}} \right)\left[ E \right]$$

$$\left[ E_{t} \right]=\left( \frac{\left( k_{1}^{'}+k_{2} \right)+k_{1}\left[ A \right]+k_{2}^{'}\left[ P_{1} \right]\ldots\left[ P_{N} \right]}{k_{1}^{'}+k_{2}} \right)\left[ E \right]$$

$$\left[ E_{t} \right]=\left( k_{1}^{'}+k_{2}+\frac{k_{1}}{k_{1}^{'}+k_{2}}\left[ A \right]+\frac{k_{2}^{'}}{k_{1}^{'}+k_{2}}\left[ P_{1} \right]\ldots\left[ P_{N} \right] \right)\left[ E \right]$$

$$\frac{\left[ E_{t} \right]}{1+\frac{k_{1}}{k_{1}^{'}+k_{2}}\left[ A \right]+\frac{k_{2}^{'}}{k_{1}^{'}+k_{2}}\left[ P_{1} \right]\ldots\left[ P_{N} \right]}=[E]$$

Making the substitution above into $\left[ EA \right]$:

$$\left[ EA \right]=\left( \frac{k_{1}\left[ A \right]}{k_{1}^{'}+k_{2}}+\frac{k_{2}^{'}\left[ P_{1} \right]\ldots\left[ P_{N} \right]}{k_{1}^{'}+k_{2}} \right)\left[ E \right]$$

$$\left[ EA \right]=\left( \frac{k_{1}\left[ A \right]+k_{2}^{'}\left[ P_{1} \right]\ldots\left[ P_{N} \right]}{k_{1}^{'}+k_{2}} \right)\left[ E \right]$$

$$\left[ EA \right]=\left( \frac{k_{1}\left[ A \right]+k_{2}^{'}\left[ P_{1} \right]\ldots\left[ P_{N} \right]}{k_{1}^{'}+k_{2}} \right)\left( \frac{\left[ E_{t} \right]}{1+\frac{k_{1}}{k_{1}^{'}+k_{2}}\left[ A \right]+\frac{k_{2}^{'}}{k_{1}^{'}+k_{2}}\left[ P_{1} \right]\ldots\left[ P_{N} \right]} \right)$$

$$\left[ EA \right]=\left( \frac{k_{1}\left[ A \right]\left[ E_{t} \right]+k_{2}^{'}\left[ P_{1} \right]\ldots\left[ P_{N} \right]\left[ E_{t} \right]}{\left( k_{1}^{'}+k_{2} \right)\left( 1+\frac{k_{1}}{k_{1}^{'}+k_{2}}\left[ A \right]+\frac{k_{2}^{'}}{k_{1}^{'}+k_{2}}\left[ P_{1} \right]\ldots\left[ P_{N} \right] \right)} \right)$$

$$\left[ EA \right]=\left( \frac{k_{1}\left[ A \right]\left[ E_{t} \right]+k_{2}^{'}\left[ P_{1} \right]\ldots\left[ P_{N} \right]\left[ E_{t} \right]}{\left( k_{1}^{'}+k_{2} \right)+\frac{k_{1}\left( k_{1}^{'}+k_{2} \right)}{k_{1}^{'}+k_{2}}\left[ A \right]+\frac{k_{2}^{'}\left( k_{1}^{'}+k_{2} \right)}{k_{1}^{'}+k_{2}}\left[ P_{1} \right]\ldots\left[ P_{N} \right]} \right)$$

$$\left[ EA \right]=\left( \frac{k_{1}\left[ A \right]\left[ E_{t} \right]+k_{2}^{'}\left[ P_{1} \right]\ldots\left[ P_{N} \right]\left[ E_{t} \right]}{\left( k_{1}^{'}+k_{2} \right)+k_{1}\left[ A \right]+k_{2}^{'}\left[ P_{1} \right]\ldots\left[ P_{N} \right]} \right)$$

Finally the reaction velocity is:

$$v=k_{2}\left[ EA \right]-k_{2}^{'}\left[ E \right]\left[ P_{1} \right]\ldots\left[ P_{N} \right]$$

$$v=\left( \frac{k_{1}\left[ A \right]\left[ E_{t} \right]+k_{2}^{'}\left[ P_{1} \right]\ldots\left[ P_{N} \right]\left[ E_{t} \right]}{\left( \left( k_{1}^{'}+k_{2} \right)+k_{1}\left[ A \right]+k_{2}^{'}\left[ P_{1} \right]\ldots\left[ P_{N} \right] \right)} \right)-\frac{k_{2}^{'}\left[ E_{t} \right]\left[ P_{1} \right]\ldots\left[ P_{N} \right]}{1+\frac{k_{1}}{k_{1}^{'}+k_{2}}\left[ A \right]+\frac{k_{2}^{'}}{k_{1}^{'}+k_{2}}\left[ P_{1} \right]\ldots\left[ P_{N} \right]}$$

$$v=\left( \frac{k_{1}\left[ A \right]\left[ E_{t} \right]+k_{2}^{'}\left[ P_{1} \right]\ldots\left[ P_{N} \right]\left[ E_{t} \right]}{\left( k_{1}^{'}+k_{2} \right)\left( 1+\frac{k_{1}}{k_{1}^{'}+k_{2}}\left[ A \right]+\frac{k_{2}^{'}}{k_{1}^{'}+k_{2}}\left[ P_{1} \right]\ldots\left[ P_{N} \right] \right)} \right)-\frac{k_{2}^{'}\left[ E_{t} \right]\left[ P_{1} \right]\ldots\left[ P_{N} \right]}{1+\frac{k_{1}}{k_{1}^{'}+k_{2}}\left[ A \right]+\frac{k_{2}^{'}}{k_{1}^{'}+k_{2}}\left[ P_{1} \right]\ldots\left[ P_{N} \right]}$$

$$v=\left( \frac{\frac{k_{1}}{k_{1}^{'}+k_{2}}\left[ A \right]\left[ E_{t} \right]+\frac{k_{2}^{'}}{k_{1}^{'}+k_{2}}\left[ P_{1} \right]\ldots\left[ P_{N} \right]\left[ E_{t} \right]}{1+\frac{k_{1}}{k_{1}^{'}+k_{2}}\left[ A \right]+\frac{k_{2}^{'}}{k_{1}^{'}+k_{2}}\left[ P_{1} \right]\ldots\left[ P_{N} \right]} \right)-\frac{k_{2}^{'}\left[ E_{t} \right]\left[ P_{1} \right]\ldots\left[ P_{N} \right]}{1+\frac{k_{1}}{k_{1}^{'}+k_{2}}\left[ A \right]+\frac{k_{2}^{'}}{k_{1}^{'}+k_{2}}\left[ P_{1} \right]\ldots\left[ P_{N} \right]}$$

$$v=\left( \frac{\frac{k_{1}}{k_{1}^{'}+k_{2}}\left[ A \right]\left[ E_{t} \right]+\left( \frac{k_{2}^{'}}{k_{1}^{'}+k_{2}}-k_{2}^{'} \right)\left[ P_{1} \right]\ldots\left[ P_{N} \right]\left[ E_{t} \right]}{1+\frac{k_{1}}{k_{1}^{'}+k_{2}}\left[ A \right]+\frac{k_{2}^{'}}{k_{1}^{'}+k_{2}}\left[ P_{1} \right]\ldots\left[ P_{N} \right]} \right)$$

$$v=\frac{\frac{k_{1}}{k_{1}^{'}+k_{2}}\left[ A \right]\left[ E_{t} \right]+\left( \frac{k_{2}^{'}}{k_{1}^{'}+k_{2}}-k_{2}^{'} \right)\left[ P_{1} \right]\ldots\left[ P_{N} \right]\left[ E_{t} \right]}{1+\frac{k_{1}}{k_{1}^{'}+k_{2}}\left[ A \right]+\frac{k_{2}^{'}}{k_{1}^{'}+k_{2}}\left[ P_{1} \right]\ldots\left[ P_{N} \right]}$$

$$Let:$$

$$K_{1}=\frac{k_{1}}{k_{1}^{'}+k_{2}}\left[ E_{t} \right]\geq0$$

$$K_{2}=\left( k_{2}^{'}-\frac{k_{2}^{'}}{k_{1}^{'}+k_{2}} \right)\left[ E_{t} \right]\mathbb{\in R}$$

$$K_{3}=\frac{k_{1}}{k_{1}^{'}+k_{2}}\geq0$$

$$K_{4}=\frac{k_{2}^{'}}{k_{1}^{'}+k_{2}}\geq0$$

$$v=\frac{K_{1}\left[ A \right]-K_{2}\left[ P_{1} \right]\ldots\left[ P_{N} \right]}{1+K_{3}\left[ A \right]+K_{4}\left[ P_{1} \right]\ldots\left[ P_{N} \right]}$$

Note that the Briggs-Haldane Equation for this type of enzyme kinetics is as follows:

$$-r_{A}=\frac{V_{max}\left( \left[ A \right]-\frac{\left[ P \right]}{K_{c}} \right)}{\left[ A \right]+K_{m}+K_{p}\left[ P \right]}$$

Note that our equation can be re-arranged to these values:

$$v=-r_{A}=\frac{K_{1}\left( \left[ A \right]-\frac{K_{2}}{K_{1}}\left[ P_{1} \right]\ldots\left[ P_{N} \right] \right)}{K_{3}\left( \frac{1}{K_{3}}+\left[ A \right]+\frac{K_{4}}{K_{3}}\left[ P_{1} \right]\ldots\left[ P_{N} \right] \right)}$$

$$-r_{A}=\frac{\frac{K_{1}}{K_{3}}\left( \left[ A \right]-\frac{\left[ P_{1} \right]\ldots\left[ P_{N} \right]}{\frac{K_{1}}{K_{2}}} \right)}{\left( \frac{1}{K_{3}}+\left[ A \right]+\frac{K_{4}}{K_{3}}\left[ P_{1} \right]\ldots\left[ P_{N} \right] \right)}$$

$$-r_{A}=\frac{\frac{K_{1}}{K_{3}}\left( \left[ A \right]-\frac{\left[ P_{1} \right]\ldots\left[ P_{N} \right]}{\frac{K_{1}}{K_{2}}} \right)}{\left( \frac{1}{K_{3}}+\left[ A \right]+\frac{K_{4}}{K_{3}}\left[ P_{1} \right]\ldots\left[ P_{N} \right] \right)}$$

Let:

$$V_{max}=\frac{K_{1}}{K_{3}}$$

$$K_{c}=-\frac{K_{1}}{K_{2}}$$

$$K_{m}=\frac{1}{K_{3}}$$

$$K_{p}=\frac{K_{4}}{K_{3}}$$

Therefore:

$$-r_{A}=\frac{V_{max}\left( \left[ A \right]-\frac{\left[ P_{1} \right]\ldots\left[ P_{N} \right]}{K_{c}} \right)}{\left( K_{m}+\left[ A \right]+K_{p}\left[ P_{1} \right]\ldots\left[ P_{N} \right] \right)}$$

$$-r_{A}=\frac{V_{max}\left( \left[ A \right]-\frac{\left[ P_{1} \right]\ldots\left[ P_{N} \right]}{K_{c}} \right)}{\left( \left[ A \right]+K_{m}+K_{p}\left[ P_{1} \right]\ldots\left[ P_{N} \right] \right)}$$

Our reaction form is used as it avoid issues of division by zero errors that can occur if the Briggs-Haldane Equation is used with variable kinetic parameters.

##### **2.2.2 Reversible Single-Substrate Allosteric Activation Kinetics**

As with the irreversible allosteric activation kinetics, the enzyme must be activated before substrate binding. The major difference is that the final product-forming step is now reversible.

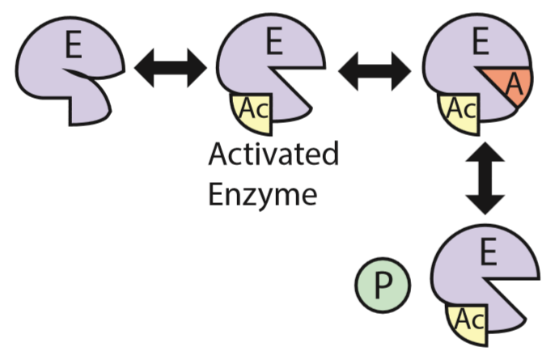

$$E+\alpha\rightleftharpoons E\alpha,k_{1}\to,k_{1}^{'}\leftarrow$$

$$E\alpha+A\rightleftharpoons E\alpha A, k_{2}\to,k_{2}^{'}\leftarrow$$

$$E\alpha A\rightleftharpoons E\alpha+P,k_{3}\to,k_{3}^{'}\leftarrow$$

From these elementary steps it can be shown that the reaction velocity and enzyme complex formation rates can be given as:

$$r_{E\alpha}=k_{1}\left[ E \right]\left[ \alpha\right]-k_{1}^{'}\left[ E\alpha\right]-k_{2}\left[ E\alpha\right]\left[ A \right]+k_{2}^{'}\left[ E\alpha A \right]+k_{3}\left[ E\alpha A \right]-k_{3}^{'}[E\alpha]\left[ P_{1} \right]\ldots\left[ P_{N} \right]$$

$$r_{E\alpha A}=k_{2}\left[ E\alpha\right]\left[ A \right]-k_{2}^{'}\left[ E\alpha A \right]-k_{3}\left[ E\alpha A \right]+k_{3}^{'}[E\alpha]\left[ P_{1} \right]\ldots\left[ P_{N} \right]$$

$$v=k_{3}\left[ E\alpha A \right]-k_{3}^{'}[E\alpha]\left[ P_{1} \right]\ldots\left[ P_{N} \right]$$

Solving for enzyme complex concentrations using the PSSH

$$r_{E\alpha}=0=k_{1}\left[ E \right]\left[ \alpha\right]-(k_{1}^{'}+k_{2}\left[ A \right]+k_{3}^{'}\left[ P_{1} \right]\ldots\left[ P_{N} \right])\left[ E\alpha\right]+\left( k_{2}^{'}+k_{3} \right)\left[ E\alpha A \right]$$

$$\left( k_{1}^{'}+k_{2}\left[ A \right]+k_{3}^{'}\left[ P_{1} \right]\ldots\left[ P_{N} \right] \right)\left[ E\alpha\right]=k_{1}\left[ E \right]\left[ \alpha\right]+\left( k_{2}^{'}+k_{3} \right)\left[ E\alpha A \right]$$

$$\left[ E\alpha\right]=\frac{k_{1}\left[ E \right]\left[ \alpha\right]}{k_{1}^{'}+k_{2}\left[ A \right]+k_{3}^{'}\left[ P_{1} \right]\ldots\left[ P_{N} \right]}+\frac{\left( k_{2}^{'}+k_{3} \right)}{k_{1}^{'}+k_{2}\left[ A \right]+k_{3}^{'}\left[ P_{1} \right]\ldots\left[ P_{N} \right]}\left[ E\alpha A \right]$$

$$r_{E\alpha A}=0=\left( k_{2}\left[ A \right]+k_{3}^{'}\left[ P_{1} \right]\ldots\left[ P_{N} \right] \right)\left[ E\alpha\right]-\left( k_{2}^{'}+k_{3} \right)\left[ E\alpha A \right]$$

$$\left( k_{2}^{'}+k_{3} \right)\left[ E\alpha A \right]=\left( k_{2}\left[ A \right]+k_{3}^{'}\left[ P_{1} \right]\ldots\left[ P_{N} \right] \right)\left[ E\alpha\right]$$

$$\left[ E\alpha A \right]=\frac{k_{2}\left[ A \right]+k_{3}^{'}\left[ P_{1} \right]\ldots\left[ P_{N} \right]}{k_{2}^{'}+k_{3}}\left[ E\alpha\right]$$

$$\left[ E\alpha\right]=\frac{k_{1}\left[ E \right]\left[ \alpha\right]}{k_{1}^{'}+k_{2}\left[ A \right]+k_{3}^{'}\left[ P_{1} \right]\ldots\left[ P_{N} \right]}+\frac{\left( k_{2}^{'}+k_{3} \right)}{k_{1}^{'}+k_{2}\left[ A \right]+k_{3}^{'}\left[ P_{1} \right]\ldots\left[ P_{N} \right]}\frac{k_{2}\left[ A \right]+k_{3}^{'}\left[ P_{1} \right]\ldots\left[ P_{N} \right]}{k_{2}^{'}+k_{3}}\left[ E\alpha\right]$$

$$\left[ E\alpha\right]=\frac{k_{1}\left[ E \right]\left[ \alpha\right]}{k_{1}^{'}+k_{2}\left[ A \right]+k_{3}^{'}\left[ P_{1} \right]\ldots\left[ P_{N} \right]}+\frac{k_{2}\left[ A \right]+k_{3}^{'}\left[ P_{1} \right]\ldots\left[ P_{N} \right]}{k_{1}^{'}+k_{2}\left[ A \right]+k_{3}^{'}\left[ P_{1} \right]\ldots\left[ P_{N} \right]}\left[ E\alpha\right]$$

$$\left( 1-\frac{k_{2}\left[ A \right]+k_{3}^{'}\left[ P_{1} \right]\ldots\left[ P_{N} \right]}{k_{1}^{'}+k_{2}\left[ A \right]+k_{3}^{'}\left[ P_{1} \right]\ldots\left[ P_{N} \right]} \right)\left[ E\alpha\right]=\frac{k_{1}\left[ E \right]\left[ \alpha\right]}{k_{1}^{'}+k_{2}\left[ A \right]+k_{3}^{'}\left[ P_{1} \right]\ldots\left[ P_{N} \right]}$$

$$\left( \frac{k_{1}^{'}+k_{2}\left[ A \right]+k_{3}^{'}\left[ P_{1} \right]\ldots\left[ P_{N} \right]-k_{2}\left[ A \right]-k_{3}^{'}\left[ P_{1} \right]\ldots\left[ P_{N} \right]}{k_{1}^{'}+k_{2}\left[ A \right]+k_{3}^{'}\left[ P_{1} \right]\ldots\left[ P_{N} \right]} \right)\left[ E\alpha\right]=\frac{k_{1}\left[ E \right]\left[ \alpha\right]}{k_{1}^{'}+k_{2}\left[ A \right]+k_{3}^{'}\left[ P_{1} \right]\ldots\left[ P_{N} \right]}$$

$$\left( \frac{k_{1}^{'}}{k_{1}^{'}+k_{2}\left[ A \right]+k_{3}^{'}\left[ P_{1} \right]\ldots\left[ P_{N} \right]} \right)\left[ E\alpha\right]=\frac{k_{1}\left[ E \right]\left[ \alpha\right]}{k_{1}^{'}+k_{2}\left[ A \right]+k_{3}^{'}\left[ P_{1} \right]\ldots\left[ P_{N} \right]}$$

$$\left[ E\alpha\right]=\frac{k_{1}\left[ E \right]\left[ \alpha\right]}{k_{1}^{'}+k_{2}\left[ A \right]+k_{3}^{'}\left[ P_{1} \right]\ldots\left[ P_{N} \right]}\frac{k_{1}^{'}+k_{2}\left[ A \right]+k_{3}^{'}\left[ P_{1} \right]\ldots\left[ P_{N} \right]}{k_{1}^{'}}$$

$$\left[ E\alpha\right]=\frac{k_{1}}{k_{1}^{'}}\left[ E \right]\left[ \alpha\right]$$

$$\left[ E\alpha A \right]=\frac{k_{2}\left[ A \right]+k_{3}^{'}\left[ P_{1} \right]\ldots\left[ P_{N} \right]}{k_{2}^{'}+k_{3}}\left[ E\alpha\right]=\frac{k_{2}\left[ A \right]+k_{3}^{'}\left[ P_{1} \right]\ldots\left[ P_{N} \right]}{k_{2}^{'}+k_{3}}\frac{k_{1}}{k_{1}^{'}}\left[ E \right]\left[ \alpha\right]$$

$$\left[ E\alpha A \right]=\frac{k_{1}\left( k_{2}\left[ A \right]+k_{3}^{'}\left[ P_{1} \right]\ldots\left[ P_{N} \right] \right)}{k_{1}^{'}\left( k_{2}^{'}+k_{3} \right)}\left[ E \right]\left[ \alpha\right]$$

Since un-complexed enzymes are difficult to measure precisely, use the total enzyme concentration and solve for the uncomplexed enzyme concentration:

$$\left[ E_{t} \right]=\left[ E \right]+\left[ E\alpha\right]+\left[ E\alpha A \right]$$

$$\left[ E_{t} \right]=\left[ E \right]+\frac{k_{1}}{k_{1}^{'}}\left[ E \right]\left[ \alpha\right]+\frac{k_{1}\left( k_{2}\left[ A \right]+k_{3}^{'}\left[ P_{1} \right]\ldots\left[ P_{N} \right] \right)}{k_{1}^{'}\left( k_{2}^{'}+k_{3} \right)}\left[ E \right]\left[ \alpha\right]$$

$$\left[ E_{t} \right]=\left[ E \right]\left( 1+\frac{k_{1}}{k_{1}^{'}}\left[ \alpha\right]+\frac{k_{1}\left( k_{2}\left[ A \right]+k_{3}^{'}[\left[ P_{1} \right]\ldots\left[ P_{N} \right] \right)}{k_{1}^{'}\left( k_{2}^{'}+k_{3} \right)}\left[ \alpha\right] \right)$$

$$\left[ E_{t} \right]=\left[ E \right]\left( \frac{k_{1}^{'}\left( k_{2}^{'}+k_{3} \right)}{k_{1}^{'}\left( k_{2}^{'}+k_{3} \right)}+\frac{k_{1}\left( k_{2}^{'}+k_{3} \right)\left[ \alpha\right]}{k_{1}^{'}\left( k_{2}^{'}+k_{3} \right)}+\frac{k_{1}\left( k_{2}\left[ A \right]+k_{3}^{'}\left[ P_{1} \right]\ldots\left[ P_{N} \right] \right)\left[ \alpha\right]}{k_{1}^{'}\left( k_{2}^{'}+k_{3} \right)} \right)$$

$$\left[ E_{t} \right]=\left[ E \right]\left( \frac{k_{1}^{'}\left( k_{2}^{'}+k_{3} \right)+k_{1}\left( k_{2}^{'}+k_{3} \right)\left[ \alpha\right]+k_{1}\left( k_{2}\left[ A \right]+k_{3}^{'}\left[ P_{1} \right]\ldots\left[ P_{N} \right] \right)\left[ \alpha\right]}{k_{1}^{'}\left( k_{2}^{'}+k_{3} \right)} \right)$$

$$\left[ E \right]=\left[ E_{t} \right]\left( \frac{k_{1}^{'}\left( k_{2}^{'}+k_{3} \right)}{k_{1}^{'}\left( k_{2}^{'}+k_{3} \right)+k_{1}\left( k_{2}^{'}+k_{3} \right)\left[ \alpha\right]+k_{1}\left( k_{2}\left[ A \right]+k_{3}^{'}\left[ P_{1} \right]\ldots\left[ P_{N} \right] \right)\left[ \alpha\right]} \right)$$

Substituting above results into the reaction velocity yields:

$$v=k_{3}\left[ E\alpha A \right]-k_{3}^{'}[E\alpha]\left[ P_{1} \right]\ldots\left[ P_{N} \right]$$

$$v=\frac{k_{1}k_{3}\left( k_{2}\left[ A \right]+k_{3}^{'}\left[ P_{1} \right]\ldots\left[ P_{N} \right] \right)}{k_{1}^{'}\left( k_{2}^{'}+k_{3} \right)}\left[ E \right]\left[ \alpha\right]-\frac{k_{1}k_{3}^{'}}{k_{1}^{'}}\left[ E \right]\left[ \alpha\right]\left[ P_{1} \right]\ldots\left[ P_{N} \right]$$

$$v=\left[ E \right]\left[ \alpha\right]\left( \frac{k_{1}k_{3}\left( k_{2}\left[ A \right]+k_{3}^{'}\left[ P_{1} \right]\ldots\left[ P_{N} \right] \right)}{k_{1}^{'}\left( k_{2}^{'}+k_{3} \right)}-\frac{k_{1}k_{3}^{'}\left[ P_{1} \right]\ldots\left[ P_{N} \right]}{k_{1}^{'}} \right)$$

$$v=\left[ E \right]\left[ \alpha\right]\left( \frac{k_{1}k_{3}\left( k_{2}\left[ A \right]+k_{3}^{'}\left[ P_{1} \right]\ldots\left[ P_{N} \right] \right)}{k_{1}^{'}\left( k_{2}^{'}+k_{3} \right)}-\frac{k_{1}k_{3}^{'}\left( k_{2}^{'}+k_{3} \right)\left[ P_{1} \right]\ldots\left[ P_{N} \right]}{k_{1}^{'}\left( k_{2}^{'}+k_{3} \right)} \right)$$

$$v=\left[ E \right]\left[ \alpha\right]\left( \frac{k_{1}k_{2}k_{3}\left[ A \right]+k_{1}k_{3}^{'}k_{3}\left[ P \right]-k_{1}k_{2}^{'}k_{3}^{'}\left[ P_{1} \right]\ldots\left[ P_{N} \right]-k_{1}k_{3}^{'}k_{3}\left[ P_{1} \right]\ldots\left[ P_{N} \right]}{k_{1}^{'}\left( k_{2}^{'}+k_{3} \right)} \right)$$

$$v=\left[ E \right]\left[ \alpha\right]\left( \frac{k_{1}k_{2}k_{3}\left[ A \right]-k_{1}k_{2}^{'}k_{3}^{'}\left[ P_{1} \right]\ldots\left[ P_{N} \right]}{k_{1}^{'}\left( k_{2}^{'}+k_{3} \right)} \right)$$

$$v=\left[ \alpha\right]\left[ E_{t} \right]\left( \frac{k_{1}^{'}\left( k_{2}^{'}+k_{3} \right)}{k_{1}^{'}\left( k_{2}^{'}+k_{3} \right)+k_{1}\left( k_{2}^{'}+k_{3} \right)\left[ \alpha\right]+k_{1}\left( k_{2}\left[ A \right]+k_{3}^{'}\left[ P_{1} \right]\ldots\left[ P_{N} \right] \right)\left[ \alpha\right]} \right)\left( \frac{k_{1}k_{2}k_{3}\left[ A \right]-k_{1}k_{2}^{'}k_{3}^{'}\left[ P_{1} \right]\ldots\left[ P_{N} \right]}{k_{1}^{'}\left( k_{2}^{'}+k_{3} \right)} \right)$$

$$v=\left[ \alpha\right]\left[ E_{t} \right]\left( \frac{k_{1}k_{2}k_{3}\left[ A \right]-k_{1}k_{2}^{'}k_{3}^{'}\left[ P_{1} \right]\ldots\left[ P_{N} \right]}{k_{1}^{'}\left( k_{2}^{'}+k_{3} \right)+k_{1}\left( k_{2}^{'}+k_{3} \right)\left[ \alpha\right]+k_{1}\left( k_{2}\left[ A \right]+k_{3}^{'}\left[ P_{1} \right]\ldots\left[ P_{N} \right] \right)\left[ \alpha\right]} \right)$$

$$v=\left[ \alpha\right]\left( \frac{k_{1}k_{2}k_{3}\left[ E_{t} \right]\left[ A \right]-k_{1}k_{2}^{'}k_{3}^{'}\left[ E_{t} \right]\left[ P_{1} \right]\ldots\left[ P_{N} \right]}{k_{1}^{'}\left( k_{2}^{'}+k_{3} \right)\left( 1+\frac{k_{1}}{k_{1}^{'}}\left[ \alpha\right]+\frac{k_{1}k_{2}}{k_{1}^{'}\left( k_{2}^{'}+k_{3} \right)}\left[ A \right]\left[ \alpha\right]+\frac{k_{1}k_{3}^{'}}{k_{1}^{'}\left( k_{2}^{'}+k_{3} \right)}\left[ P_{1} \right]\ldots\left[ P_{N} \right]\left[ \alpha\right] \right)} \right)$$

$$v=\left[ \alpha\right]\left( \frac{\frac{k_{1}k_{2}k_{3}}{k_{1}^{'}\left( k_{2}^{'}+k_{3} \right)}\left[ E_{t} \right]\left[ A \right]-\frac{k_{1}k_{2}^{'}k_{3}^{'}}{k_{1}^{'}\left( k_{2}^{'}+k_{3} \right)}\left[ E_{t} \right]\left[ P_{1} \right]\ldots\left[ P_{N} \right]}{1+\frac{k_{1}}{k_{1}^{'}}\left[ \alpha\right]+\frac{k_{1}k_{2}}{k_{1}^{'}\left( k_{2}^{'}+k_{3} \right)}\left[ A \right]\left[ \alpha\right]+\frac{k_{1}k_{3}^{'}}{k_{1}^{'}\left( k_{2}^{'}+k_{3} \right)}\left[ P_{1} \right]\ldots\left[ P_{N} \right]\left[ \alpha\right]} \right)$$

The final equation for reaction velocity is then:

$$\boldsymbol{v=}\frac{\boldsymbol{k}_{\boldsymbol{cat,f}}\left[ \boldsymbol{E}_{\boldsymbol{t}} \right]\left[ \boldsymbol{A} \right]\left[ \boldsymbol{\alpha} \right]\boldsymbol{-}\boldsymbol{k}_{\boldsymbol{cat,b}}\left[ \boldsymbol{E}_{\boldsymbol{t}} \right]\left[ \boldsymbol{P}_{\boldsymbol{1}} \right]\boldsymbol{\ldots}\left[ \boldsymbol{P}_{\boldsymbol{N}} \right]\left[ \boldsymbol{\alpha} \right]}{\boldsymbol{1+}\boldsymbol{K}_{\boldsymbol{\alpha}}\left[ \boldsymbol{\alpha} \right]\boldsymbol{+}\boldsymbol{K}_{\boldsymbol{\alpha A}}\left[ \boldsymbol{A} \right]\left[ \boldsymbol{\alpha} \right]\boldsymbol{+}\boldsymbol{K}_{\boldsymbol{\alpha P}}\left[ \boldsymbol{P}_{\boldsymbol{1}} \right]\boldsymbol{\ldots}\left[ \boldsymbol{P}_{\boldsymbol{N}} \right]\left[ \boldsymbol{\alpha} \right]}\boldsymbol{=}\frac{\boldsymbol{V}_{\boldsymbol{max,f}}\left[ \boldsymbol{A} \right]\left[ \boldsymbol{\alpha} \right]\boldsymbol{-}\boldsymbol{V}_{\boldsymbol{max,b}}\left[ \boldsymbol{P}_{\boldsymbol{1}} \right]\boldsymbol{\ldots}\left[ \boldsymbol{P}_{\boldsymbol{N}} \right]\left[ \boldsymbol{\alpha} \right]}{\boldsymbol{1+}\boldsymbol{K}_{\boldsymbol{\alpha}}\left[ \boldsymbol{\alpha} \right]\boldsymbol{+}\boldsymbol{K}_{\boldsymbol{\alpha A}}\left[ \boldsymbol{A} \right]\left[ \boldsymbol{\alpha} \right]\boldsymbol{+}\boldsymbol{K}_{\boldsymbol{\alpha P}}\left[ \boldsymbol{P}_{\boldsymbol{1}} \right]\boldsymbol{\ldots}\left[ \boldsymbol{P}_{\boldsymbol{N}} \right]\left[ \boldsymbol{\alpha} \right]}$$

$$\boldsymbol{parameters:}\boldsymbol{V}_{\boldsymbol{max,f}}\boldsymbol{,}\boldsymbol{V}_{\boldsymbol{max,b}}\boldsymbol{,}\left[ \boldsymbol{A} \right]\boldsymbol{,}\left[ \boldsymbol{\alpha} \right]\left[ \boldsymbol{P}_{\boldsymbol{1}} \right]\boldsymbol{,\ldots,}\left[ \boldsymbol{P}_{\boldsymbol{N}} \right]$$

$$\boldsymbol{variables:}\boldsymbol{K}_{\boldsymbol{\alpha}}\boldsymbol{,}\boldsymbol{K}_{\boldsymbol{\alpha A}}\boldsymbol{,}\boldsymbol{K}_{\boldsymbol{\alpha P}}$$

##### **2.2.3 Reversible Single-Substrate Allosteric Inhibition Kinetics**

Like the other reversible kinetics here derived, the main difference is that the product-forming step is reversible and so the entire enzyme-catalyzed reaction is now reversible.

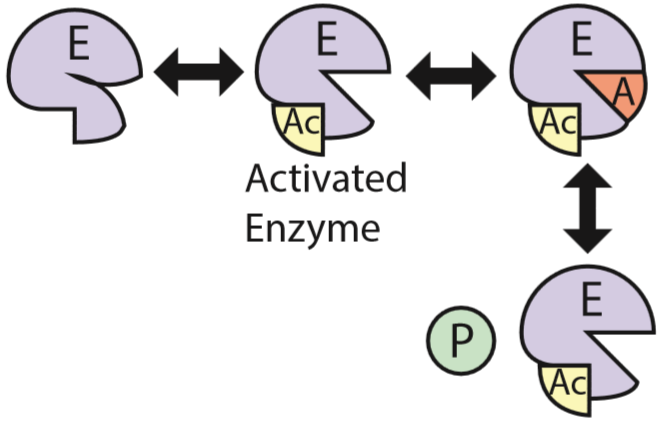

$$E+I\rightleftharpoons EI,k_{1}\to,k_{1}^{'}\leftarrow$$

$$E+A\rightleftharpoons EA,k_{2}\to,k_{2}^{'}\leftarrow$$

$$EA\rightleftharpoons E+P,k_{3}\to,k_{3}^{'}\leftarrow$$

This mechanism and elementary equations result in reaction rates for the enzyme complexes of:

$$r_{EA}=k_{2}\left[ E \right]\left[ A \right]-k_{2}^{'}\left[ EA \right]-k_{3}\left[ EA \right]+k_{3}^{'}[E]\left[ P_{1} \right]\ldots\left[ P_{N} \right]$$

$$r_{EI}=k_{1}\left[ E \right]\left[ I \right]-k_{1}^{'}[EI]$$

$$v=k_{3}\left[ EA \right]-k_{3}^{'}[E]\left[ P_{1} \right]\ldots\left[ P_{N} \right]$$

Solving for enzyme complex concentration by assuming the PSSH gives:

$$r_{EA}=0=k_{2}\left[ E \right]\left[ A \right]-k_{2}^{'}\left[ EA \right]-k_{3}\left[ EA \right]+k_{3}^{'}[E]\left[ P_{1} \right]\ldots\left[ P_{N} \right]$$

$$0=k_{2}\left[ E \right]\left[ A \right]-\left( k_{2}^{'}+k_{3} \right)\left[ EA \right]+k_{3}^{'}[E]\left[ P_{1} \right]\ldots\left[ P_{N} \right]$$

$$\left( k_{2}^{'}+k_{3} \right)\left[ EA \right]=k_{2}\left[ E \right]\left[ A \right]+k_{3}^{'}[E]\left[ P_{1} \right]\ldots\left[ P_{N} \right]$$

$$\left[ EA \right]=\frac{k_{2}\left[ E \right]\left[ A \right]+k_{3}^{'}[E]\left[ P_{1} \right]\ldots\left[ P_{N} \right]}{k_{2}^{'}+k_{3}}$$

$$r_{EI}=0=k_{1}\left[ E \right]\left[ I \right]-k_{1}^{'}[EI]$$

$$k_{1}^{'}[EI]=k_{1}\left[ E \right]\left[ I \right]$$

$$[EI]=\frac{k_{1}}{k_{1}^{'}}\left[ E \right]\left[ I \right]$$

$$v=k_{3}\left[ EA \right]-k_{3}^{'}[E]\left[ P_{1} \right]\ldots\left[ P_{N} \right]$$

$$v=k_{3}\frac{k_{2}\left[ E \right]\left[ A \right]+k_{3}^{'}[E]\left[ P_{1} \right]\ldots\left[ P_{N} \right]}{k_{2}^{'}+k_{3}}-k_{3}^{'}[E]\left[ P_{1} \right]\ldots\left[ P_{N} \right]$$

Since the fraction of uncomplexed enzyme is difficult to empirically measure, use the total enzyme concentration:

$$\left[ E_{t} \right]=\left[ E \right]+\left[ EI \right]+\left[ EA \right]$$

$$\left[ E_{t} \right]=\left[ E \right]+\frac{k_{1}}{k_{1}^{'}}\left[ E \right]\left[ I \right]+\frac{k_{2}\left[ E \right]\left[ A \right]+k_{3}^{'}\left[ E \right]\left[ P_{1} \right]\ldots\left[ P_{N} \right]}{k_{2}^{'}+k_{3}}$$

$$\left[ E_{t} \right]=\left( 1+\frac{k_{1}}{k_{1}^{'}}\left[ I \right]+\frac{k_{2}\left[ A \right]+k_{3}^{'}\left[ P_{1} \right]\ldots\left[ P_{N} \right]}{k_{2}^{'}+k_{3}} \right)\left[ E \right]$$

$$\left[ E_{t} \right]=\left( \frac{k_{1}^{'}\left( k_{2}^{'}+k_{3} \right)}{k_{1}^{'}\left( k_{2}^{'}+k_{3} \right)}+\frac{k_{1}\left( k_{2}^{'}+k_{3} \right)\left[ I \right]}{k_{1}^{'}\left( k_{2}^{'}+k_{3} \right)}+\frac{k_{1}^{'}k_{2}\left[ A \right]+k_{1}^{'}k_{3}^{'}\left[ P_{1} \right]\ldots\left[ P_{N} \right]}{k_{1}^{'}\left( k_{2}^{'}+k_{3} \right)} \right)\left[ E \right]$$

$$\left[ E_{t} \right]=\left( \frac{k_{1}^{'}\left( k_{2}^{'}+k_{3} \right)+k_{1}\left( k_{2}^{'}+k_{3} \right)\left[ I \right]+k_{1}^{'}k_{2}\left[ A \right]+k_{1}^{'}k_{3}^{'}\left[ P_{1} \right]\ldots\left[ P_{N} \right]}{k_{1}^{'}\left( k_{2}^{'}+k_{3} \right)} \right)\left[ E \right]$$

$$\left[ E \right]=\frac{k_{1}^{'}\left( k_{2}^{'}+k_{3} \right)\left[ E_{t} \right]}{k_{1}^{'}\left( k_{2}^{'}+k_{3} \right)+k_{1}\left( k_{2}^{'}+k_{3} \right)\left[ I \right]+k_{1}^{'}k_{2}\left[ A \right]+k_{1}^{'}k_{3}^{'}\left[ P_{1} \right]\ldots\left[ P_{N} \right]}$$

$$v=\left( k_{3}\frac{k_{2}\left[ A \right]+k_{3}^{'}\left[ P \right]}{k_{2}^{'}+k_{3}}-k_{3}^{'}\left[ P_{1} \right]\ldots\left[ P_{N} \right] \right)\left[ E \right]$$

$$v=\left( k_{3}\frac{k_{2}\left[ A \right]+k_{3}^{'}\left[ P_{1} \right]\ldots\left[ P_{N} \right]}{k_{2}^{'}+k_{3}}-k_{3}^{'}\left[ P_{1} \right]\ldots\left[ P_{N} \right] \right)\frac{k_{1}^{'}\left( k_{2}^{'}+k_{3} \right)\left[ E_{t} \right]}{k_{1}^{'}\left( k_{2}^{'}+k_{3} \right)+k_{1}\left( k_{2}^{'}+k_{3} \right)\left[ I \right]+k_{1}^{'}k_{2}\left[ A \right]+k_{1}^{'}k_{3}^{'}\left[ P_{1} \right]\ldots\left[ P_{N} \right]}$$

$$v=\left( k_{3}\frac{k_{2}\left[ A \right]+k_{3}^{'}\left[ P_{1} \right]\ldots\left[ P_{N} \right]}{k_{2}^{'}+k_{3}}\frac{k_{1}^{'}\left( k_{2}^{'}+k_{3} \right)\left[ E_{t} \right]}{k_{1}^{'}\left( k_{2}^{'}+k_{3} \right)+k_{1}\left( k_{2}^{'}+k_{3} \right)\left[ I \right]+k_{1}^{'}k_{2}\left[ A \right]+k_{1}^{'}k_{3}^{'}\left[ P_{1} \right]\ldots\left[ P_{N} \right]}-k_{3}^{'}\frac{k_{1}^{'}\left( k_{2}^{'}+k_{3} \right)\left[ E_{t} \right]}{k_{1}^{'}\left( k_{2}^{'}+k_{3} \right)+k_{1}\left( k_{2}^{'}+k_{3} \right)\left[ I \right]+k_{1}^{'}k_{2}\left[ A \right]+k_{1}^{'}k_{3}^{'}\left[ P_{1} \right]\ldots\left[ P_{N} \right]}\left[ P_{1} \right]\ldots\left[ P_{N} \right] \right)$$

$$v=\left( \frac{k_{1}^{'}k_{2}k_{3}\left[ A \right]\left[ E_{t} \right]+k_{1}^{'}k_{3}^{'}k_{3}\left[ P_{1} \right]\ldots\left[ P_{N} \right]\left[ E_{t} \right]}{k_{1}^{'}\left( k_{2}^{'}+k_{3} \right)+k_{1}\left( k_{2}^{'}+k_{3} \right)\left[ I \right]+k_{1}^{'}k_{2}\left[ A \right]+k_{1}^{'}k_{3}^{'}\left[ P_{1} \right]\ldots\left[ P_{N} \right]}-\frac{k_{1}^{'}k_{3}^{'}\left( k_{2}^{'}+k_{3} \right)\left[ E_{t} \right]\left[ P_{1} \right]\ldots\left[ P_{N} \right]}{k_{1}^{'}\left( k_{2}^{'}+k_{3} \right)+k_{1}\left( k_{2}^{'}+k_{3} \right)\left[ I \right]+k_{1}^{'}k_{2}\left[ A \right]+k_{1}^{'}k_{3}^{'}\left[ P_{1} \right]\ldots\left[ P_{N} \right]} \right)$$

$$v=\left( \frac{k_{1}^{'}k_{2}k_{3}\left[ A \right]\left[ E_{t} \right]+k_{1}^{'}k_{3}^{'}k_{3}\left[ P_{1} \right]\ldots\left[ P_{N} \right]\left[ E_{t} \right]-k_{1}^{'}k_{3}^{'}\left( k_{2}^{'}+k_{3} \right)\left[ E_{t} \right]\left[ P_{1} \right]\ldots\left[ P_{N} \right]}{k_{1}^{'}\left( k_{2}^{'}+k_{3} \right)+k_{1}\left( k_{2}^{'}+k_{3} \right)\left[ I \right]+k_{1}^{'}k_{2}\left[ A \right]+k_{1}^{'}k_{3}^{'}\left[ P_{1} \right]\ldots\left[ P_{N} \right]} \right)$$

$$v=\left( \frac{k_{1}^{'}k_{2}k_{3}\left[ A \right]\left[ E_{t} \right]+k_{1}^{'}k_{3}^{'}k_{3}\left[ P_{1} \right]\ldots\left[ P_{N} \right]\left[ E_{t} \right]-k_{2}^{'}k_{1}^{'}k_{3}^{'}\left[ E_{t} \right]\left[ P_{1} \right]\ldots\left[ P_{N} \right]-k_{3}k_{1}^{'}k_{3}^{'}\left[ E_{t} \right]\left[ P_{1} \right]\ldots\left[ P_{N} \right]}{k_{1}^{'}\left( k_{2}^{'}+k_{3} \right)\left( 1+\frac{k_{1}\left( k_{2}^{'}+k_{3} \right)}{k_{1}^{'}\left( k_{2}^{'}+k_{3} \right)}\left[ I \right]+\frac{k_{1}^{'}k_{2}}{k_{1}^{'}\left( k_{2}^{'}+k_{3} \right)}\left[ A \right]+\frac{k_{1}^{'}k_{3}^{'}}{k_{1}^{'}\left( k_{2}^{'}+k_{3} \right)}\left[ P_{1} \right]\ldots\left[ P_{N} \right] \right)} \right)$$

$$v=\left( \frac{\frac{k_{1}^{'}k_{2}k_{3}}{k_{1}^{'}\left( k_{2}^{'}+k_{3} \right)}\left[ A \right]\left[ E_{t} \right]-\frac{k_{2}^{'}k_{1}^{'}k_{3}^{'}}{k_{1}^{'}\left( k_{2}^{'}+k_{3} \right)}\left[ E_{t} \right]\left[ P_{1} \right]\ldots\left[ P_{N} \right]}{\left( 1+\frac{k_{1}\left( k_{2}^{'}+k_{3} \right)}{k_{1}^{'}\left( k_{2}^{'}+k_{3} \right)}\left[ I \right]+\frac{k_{1}^{'}k_{2}}{k_{1}^{'}\left( k_{2}^{'}+k_{3} \right)}\left[ A \right]+\frac{k_{1}^{'}k_{3}^{'}}{k_{1}^{'}\left( k_{2}^{'}+k_{3} \right)}\left[ P_{1} \right]\ldots\left[ P_{N} \right] \right)} \right)$$

$$\boldsymbol{v}\boldsymbol{=}\frac{\boldsymbol{k}_{\boldsymbol{cat,f}}\left[ \boldsymbol{A} \right]\left[ \boldsymbol{E}_{\boldsymbol{t}} \right]\boldsymbol{-}\boldsymbol{k}_{\boldsymbol{cat,b}}\left[ \boldsymbol{E}_{\boldsymbol{t}} \right]\left[ \boldsymbol{P}_{\boldsymbol{1}} \right]\boldsymbol{\ldots}\left[ \boldsymbol{P}_{\boldsymbol{N}} \right]}{\boldsymbol{1+}\boldsymbol{K}_{\boldsymbol{I}}\left[ \boldsymbol{I} \right]\boldsymbol{+}\boldsymbol{K}_{\boldsymbol{A}}\left[ \boldsymbol{A} \right]\boldsymbol{+}\boldsymbol{K}_{\boldsymbol{P}}\left[ \boldsymbol{P}_{\boldsymbol{1}} \right]\boldsymbol{\ldots}\left[ \boldsymbol{P}_{\boldsymbol{N}} \right]}\boldsymbol{=}\frac{\boldsymbol{V}_{\boldsymbol{max,f}}\left[ \boldsymbol{A} \right]\boldsymbol{-}\boldsymbol{V}_{\boldsymbol{max,b}}\left[ \boldsymbol{P}_{\boldsymbol{1}} \right]\boldsymbol{\ldots}\left[ \boldsymbol{P}_{\boldsymbol{N}} \right]}{\boldsymbol{1+}\boldsymbol{K}_{\boldsymbol{I}}\left[ \boldsymbol{I} \right]\boldsymbol{+}\boldsymbol{K}_{\boldsymbol{A}}\left[ \boldsymbol{A} \right]\boldsymbol{+}\boldsymbol{K}_{\boldsymbol{P}}\left[ \boldsymbol{P}_{\boldsymbol{1}} \right]\boldsymbol{\ldots}\left[ \boldsymbol{P}_{\boldsymbol{N}} \right]}$$

$$\boldsymbol{parameters:}\boldsymbol{V}_{\boldsymbol{max.f}}\boldsymbol{,}\boldsymbol{V}_{\boldsymbol{max.b}}\boldsymbol{,}\left[ \boldsymbol{P}_{\boldsymbol{1}} \right]\boldsymbol{,\ldots,}\left[ \boldsymbol{P}_{\boldsymbol{N}} \right]\boldsymbol{,}\left[ \boldsymbol{I} \right]\boldsymbol{,[A]}$$

$$\boldsymbol{variables:}\boldsymbol{K}_{\boldsymbol{I}}\boldsymbol{,}\boldsymbol{K}_{\boldsymbol{A}}\boldsymbol{,}\boldsymbol{K}_{\boldsymbol{P}}$$

#### **2.3 Irreversible Dual Substrate Kinetics**

Derivation or statement of equations to use for irreversible dual substrate kinetics.

**2.3.1 Irreversible Dual-Substrate Kinetics**

Two-substrate enzyme kinetics, irreversible final step. Assuming non-specific binding sequence (otherwise two equations would be required for this type of kinetics).

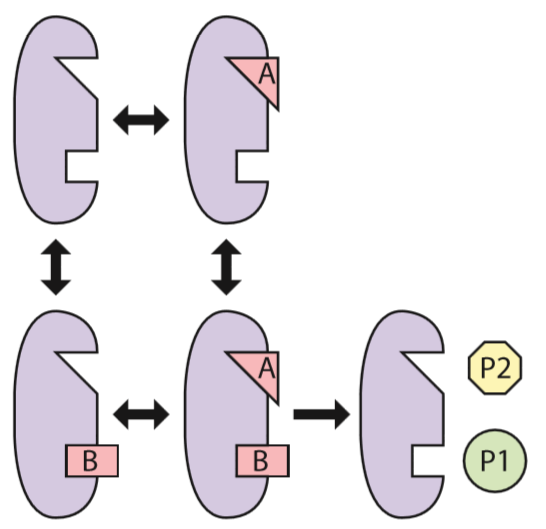

The above mechanism has the following elementary steps:

$$E+A\rightleftharpoons EA,K_{1}E+A\rightleftharpoons EA,K_{1}$$

$$E+B\rightleftharpoons EB,K_{2}E+A\rightleftharpoons EA,K_{2}$$

$$EA+B\rightleftharpoons EAB,K_{3}E+A\rightleftharpoons EA,K_{3}$$

$$EB+A\rightleftharpoons EAB,K_{4}E+A\rightleftharpoons EA,K_{4}$$

$$EAB\to E+P,k_{5}\to$$

The PSSH assumes that all reactions involving enzyme complexes have formation rates of zero. Therefore $r_{EA}=r_{EB}=r_{EAB}=0$. This indicates that the system is a pseudoequillibrium (rate of formation is equal to rate of consumption), and therefore $r_{forward}=r_{backward}$ for each reaction to which the PSSH applies.

$$K_{1}=\frac{\left[ EA \right]}{\left[ E \right]\left[ A \right]}$$

$$K_{2}=\frac{\left[ EB \right]}{\left[ E \right]\left[ B \right]}$$

$$K_{3}=\frac{\left[ EAB \right]}{\left[ EA \right]\left[ B \right]}$$

$$K_{4}=\frac{\left[ EAB \right]}{\left[ EB \right]\left[ A \right]}$$

$$v=k_{5}\left[ EAB \right]$$

Therefore:

$$\left[ EA \right]=K_{1}\left[ E \right]\left[ A \right]$$

$$\left[ EB \right]=K_{2}\left[ E \right]\left[ B \right]$$

$$K_{3}\left[ EA \right]\left[ B \right]=\left[ EAB \right]$$

$$K_{3}K_{1}\left[ E \right]\left[ A \right]\left[ B \right]=\left[ EAB \right]$$

Also,

$$K_{4}\left[ EB \right]\left[ A \right]=\left[ EAB \right]$$

$$K_{4}K_{2}\left[ E \right]\left[ B \right]\left[ A \right]=\left[ EAB \right]$$

$$\therefore K_{3}K_{1}=K_{4}K_{2}$$

Further it can be easily shown that:

$$K_{3}K_{1}=\frac{\left[ EAB \right]}{\left[ EA \right]\left[ B \right]}\frac{\left[ EA \right]}{\left[ E \right]\left[ A \right]}=\frac{\left[ EAB \right]}{\left[ E \right]\left[ A \right]\left[ B \right]}$$

And that:

$$K_{4}K_{2}=\frac{\left[ EB \right]}{\left[ E \right]\left[ B \right]}\frac{\left[ EAB \right]}{\left[ EB \right]\left[ A \right]}=\frac{\left[ EAB \right]}{\left[ E \right]\left[ A \right]\left[ B \right]}$$

Since uncomplexed enzyme concentration cannot be readily measured, substitute $[E]$ for total enzyme concentration:

$$\left[ E_{t} \right]=\left[ E \right]+\left[ EA \right]+\left[ EB \right]+\left[ EAB \right]$$

$$\left[ E_{t} \right]=\left[ E \right]+K_{1}\left[ E \right]\left[ A \right]+K_{2}\left[ E \right]\left[ B \right]+K_{3}K_{1}\left[ E \right]\left[ A \right]\left[ B \right]$$

$$\left[ E_{t} \right]=\left( 1+K_{1}\left[ A \right]+K_{2}\left[ B \right]+K_{3}K_{1}\left[ E \right]\left[ A \right]\left[ B \right] \right)\left[ E \right]$$

$$\left[ E \right]=\frac{\left[ E_{t} \right]}{1+K_{1}\left[ A \right]+K_{2}\left[ B \right]+K_{3}K_{1}\left[ A \right]\left[ B \right]}$$

Solving for the reaction velocity using the above substitutions:

$$v=k_{5}\left[ EAB \right]$$

$$v=k_{5}K_{3}K_{1}\left[ E \right]\left[ A \right]\left[ B \right]$$

$$v=\frac{k_{5}K_{3}K_{1}\left[ E_{t} \right]\left[ A \right]\left[ B \right]}{1+K_{1}\left[ A \right]+K_{2}\left[ B \right]+K_{3}K_{1}\left[ A \right]\left[ B \right]}$$

$$v=\frac{k_{5}\left[ E_{t} \right]}{\frac{1}{K_{3}K_{1}\left[ A \right]\left[ B \right]}\left( 1+K_{1}\left[ A \right]+K_{2}\left[ B \right]+K_{3}K_{1}\left[ A \right]\left[ B \right] \right)}$$

$$v=\frac{k_{5}\left[ E_{t} \right]}{\left( \frac{1}{K_{3}K_{1}\left[ A \right]\left[ B \right]}+\frac{1}{K_{3}\left[ B \right]}+\frac{K_{2}}{K_{3}K_{1}\left[ A \right]}+\frac{K_{3}K_{1}}{K_{3}K_{1}} \right)}$$

$$v=\frac{k_{5}\left[ E_{t} \right]}{\left( \frac{1}{K_{3}K_{1}\left[ A \right]\left[ B \right]}+\frac{1}{K_{3}\left[ B \right]}+\frac{K_{2}}{K_{3}K_{1}\left[ A \right]}+1 \right)}$$

$$\boldsymbol{v=}\frac{\boldsymbol{k}_{\boldsymbol{cat}}\left[ \boldsymbol{E}_{\boldsymbol{t}} \right]}{\left( \frac{\boldsymbol{K}_{\boldsymbol{AB}}}{\left[ \boldsymbol{A} \right]\left[ \boldsymbol{B} \right]}\boldsymbol{+}\frac{\boldsymbol{K}_{\boldsymbol{B}}}{\left[ \boldsymbol{B} \right]}\boldsymbol{+}\frac{\boldsymbol{K}_{\boldsymbol{A}}}{\left[ \boldsymbol{A} \right]}\boldsymbol{+1} \right)}\boldsymbol{=}\frac{\boldsymbol{V}_{\boldsymbol{max}}}{\left( \frac{\boldsymbol{K}_{\boldsymbol{AB}}}{\left[ \boldsymbol{A} \right]\left[ \boldsymbol{B} \right]}\boldsymbol{+}\frac{\boldsymbol{K}_{\boldsymbol{B}}}{\left[ \boldsymbol{B} \right]}\boldsymbol{+}\frac{\boldsymbol{K}_{\boldsymbol{A}}}{\left[ \boldsymbol{A} \right]}\boldsymbol{+1} \right)}$$

$$\boldsymbol{parameters:}\boldsymbol{V}_{\boldsymbol{max}}\boldsymbol{,}\left[ \boldsymbol{A} \right]\boldsymbol{,[B]}$$

$$\boldsymbol{variables:}\boldsymbol{K}_{\boldsymbol{A}}\boldsymbol{,}\boldsymbol{K}_{\boldsymbol{B}}\boldsymbol{,}\boldsymbol{K}_{\boldsymbol{AB}}$$

##### **2.3.2 Irreversible Dual-Substrate Allosteric Inhibition Kinetics**

Derevation of a kinetics equation for allosteric inhibition. Inhibitor binding and substrate binding are mutually exclusive in the derived model (see the kinetics drawing below).

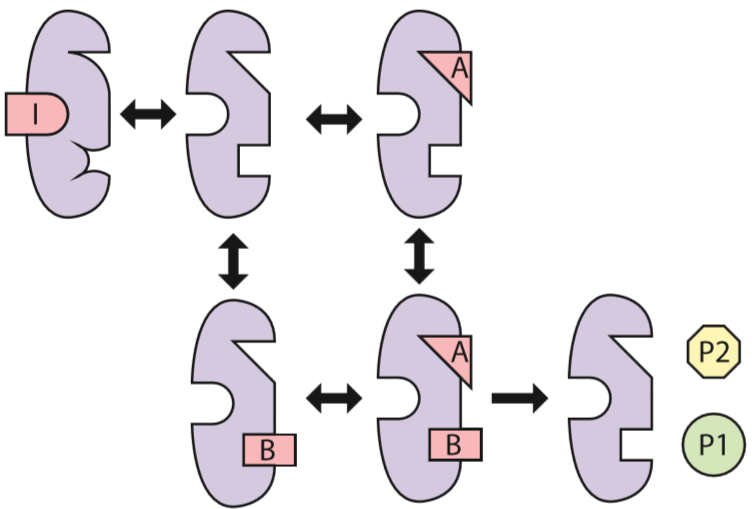

Equilibrium equations are used in the derivation, which assumes that the PSSH hypothesis applies to all reactions except the product-forming step. The equilibrium equations and the reaction velocity equation are given below:

$$K_{1}=\frac{[EI]}{\left[ E \right][I]}\to\left[ EI \right]=K_{1}\left[ E \right][I]$$

$$K_{2}=\frac{[EA]}{\left[ E \right]\left[ A \right]}\to\left[ EA \right]=K_{2}\left[ E \right]\left[ A \right]$$

$$K_{3}=\frac{[EB]}{\left[ E \right][B]}\to\left[ EB \right]=K_{3}\left[ E \right]\left[ B \right]$$

$$K_{4}=\frac{[EAB]}{\left[ EA \right][B]}\to\left[ EAB \right]=K_{4}\left[ EA \right]\left[ B \right]\to\left[ EAB \right]=K_{2}K_{4}\left[ E \right]\left[ A \right]\left[ B \right]$$

$$K_{5}=\frac{[EAB]}{\left[ EB \right][A]}\to\left[ EAB \right]=K_{5}\left[ EB \right]\left[ A \right]\to K_{3}K_{5}\left[ E \right]\left[ B \right]\left[ A \right]$$

$$\therefore K_{3}K_{5}=K_{2}K_{4}$$

$$v=k_{6}\left[ EAB \right]\to v=k_{6}K_{2}K_{4}\left[ E \right]\left[ A \right]\left[ B \right]$$

Since the concentration of uncomplexed enzymes is difficult to estimate or measure, substitute the total enzyme concentration.

$$\left[ E_{t} \right]=\left[ E \right]+\left[ EA \right]+\left[ EB \right]+\left[ EI \right]+\left[ EAB \right]$$

$$\left[ E_{t} \right]=\left[ E \right]+K_{2}\left[ E \right]\left[ A \right]+K_{3}\left[ E \right]\left[ B \right]+K_{1}\left[ E \right][I]+K_{2}K_{4}\left[ E \right]\left[ A \right]\left[ B \right]$$

$$\left[ E_{t} \right]=\left[ E \right]\left( 1+K_{2}\left[ A \right]+K_{3}\left[ B \right]+K_{1}[I]+K_{2}K_{4}\left[ A \right]\left[ B \right] \right)$$

$$\left[ E \right]=\frac{\left[ E_{t} \right]}{1+K_{2}\left[ A \right]+K_{3}\left[ B \right]+K_{1}[I]+K_{2}K_{4}\left[ A \right]\left[ B \right]}$$

$$v=k_{6}K_{2}K_{4}\left[ E \right]\left[ A \right]\left[ B \right]$$

$$v=k_{6}K_{2}K_{4}\left[ A \right]\left[ B \right]\frac{\left[ E_{t} \right]}{1+K_{2}\left[ A \right]+K_{3}\left[ B \right]+K_{1}[I]+K_{2}K_{4}\left[ A \right]\left[ B \right]}$$

$$v=\frac{k_{6}K_{2}K_{4}\left[ A \right]\left[ B \right]\left[ E_{t} \right]}{1+K_{2}\left[ A \right]+K_{3}\left[ B \right]+K_{1}[I]+K_{2}K_{4}\left[ A \right]\left[ B \right]}$$

$$\boldsymbol{v=}\frac{\boldsymbol{k}_{\boldsymbol{cat}}\left[ \boldsymbol{A} \right]\left[ \boldsymbol{B} \right]\left[ \boldsymbol{E}_{\boldsymbol{t}} \right]}{\boldsymbol{1+}\boldsymbol{K}_{\boldsymbol{A}}\left[ \boldsymbol{A} \right]\boldsymbol{+}\boldsymbol{K}_{\boldsymbol{B}}\left[ \boldsymbol{B} \right]\boldsymbol{+}\boldsymbol{K}_{\boldsymbol{I}}\boldsymbol{[I]+}\boldsymbol{K}_{\boldsymbol{AB}}\left[ \boldsymbol{A} \right]\left[ \boldsymbol{B} \right]}\boldsymbol{=}\frac{\boldsymbol{V}_{\boldsymbol{max}}\left[ \boldsymbol{A} \right]\left[ \boldsymbol{B} \right]}{\boldsymbol{1+}\boldsymbol{K}_{\boldsymbol{A}}\left[ \boldsymbol{A} \right]\boldsymbol{+}\boldsymbol{K}_{\boldsymbol{B}}\left[ \boldsymbol{B} \right]\boldsymbol{+}\boldsymbol{K}_{\boldsymbol{I}}\boldsymbol{[I]+}\boldsymbol{K}_{\boldsymbol{AB}}\left[ \boldsymbol{A} \right]\left[ \boldsymbol{B} \right]}$$

$$\boldsymbol{parameters:}\boldsymbol{V}_{\boldsymbol{max}}\boldsymbol{,}\left[ \boldsymbol{A} \right]\boldsymbol{,}\left[ \boldsymbol{B} \right]\boldsymbol{,[I]}$$

$$\boldsymbol{variables:}\boldsymbol{K}_{\boldsymbol{A}}\boldsymbol{,}\boldsymbol{K}_{\boldsymbol{B}}\boldsymbol{,}\boldsymbol{K}_{\boldsymbol{I}}\boldsymbol{,}\boldsymbol{K}_{\boldsymbol{AB}}$$

##### **2.3.3 Irreversible Dual-Substrate Allosteric Activation Kinetics**

Below is the derivation of irreversible dual substrate allosteric activation kinetics, made so primarily because the final step is irreversible (product formation step).

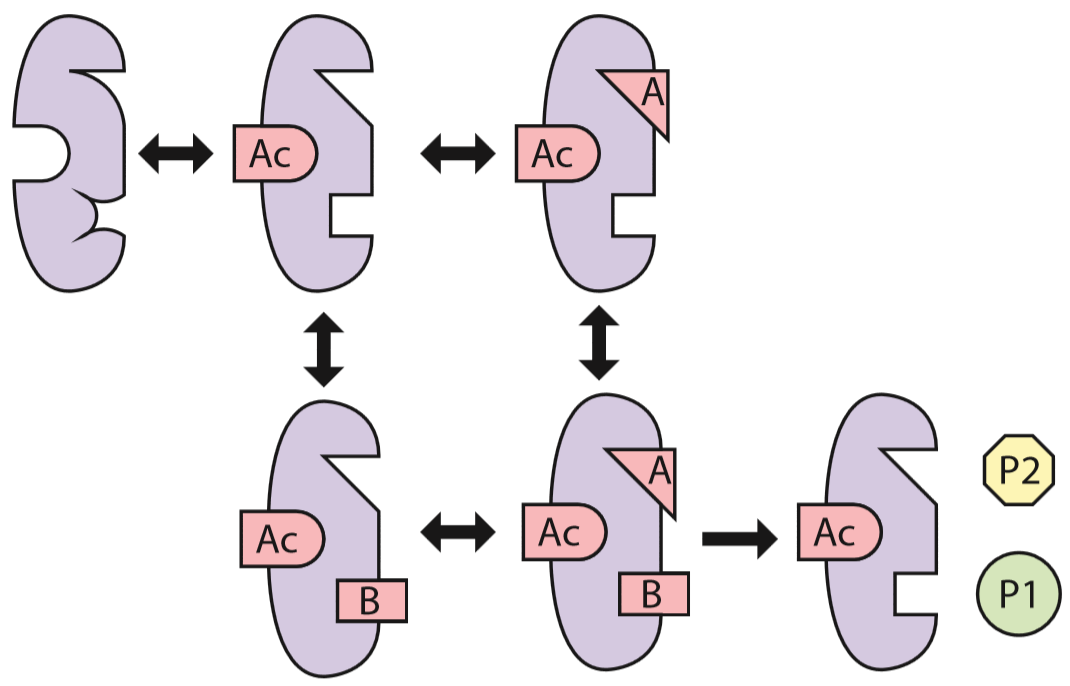

As an extension of the PSSH as applied to reactions 1, 2, 3, 4, and 5, these reactions are also at equilibrium (easily proven, but not done here). The equilibrium equations and the reaction velocity equation is as given below:

$$K_{1}=\frac{[E\alpha]}{\left[ E \right][\alpha]}\to\left[ E\alpha\right]=K_{1}\left[ E \right]\left[ \alpha\right]$$

$$K_{2}=\frac{\left[ E\alpha A \right]}{\left[ E\alpha\right]\left[ A \right]}\to\left[ E\alpha A \right]=K_{2}\left[ E\alpha\right]\left[ A \right]\to\left[ E\alpha A \right]=K_{1}K_{2}\left[ E \right]\left[ \alpha\right]\left[ A \right]$$

$$K_{3}=\frac{\left[ E\alpha B \right]}{\left[ E\alpha\right]\left[ B \right]}\to\left[ E\alpha B \right]=K_{3}\left[ E\alpha\right]\left[ B \right]\to\left[ E\alpha B \right]=K_{1}K_{3}\left[ E \right]\left[ \alpha\right]\left[ B \right]$$

$$K_{4}=\frac{\left[ E\alpha BA \right]}{\left[ E\alpha B \right]\left[ A \right]}\to\left[ E\alpha BA \right]=K_{4}\left[ E\alpha B \right][A]\to\left[ E\alpha BA \right]=K_{1}K_{3}K_{4}\left[ E \right]\left[ \alpha\right]\left[ B \right]\left[ A \right]$$

$$K_{5}=\frac{\left[ E\alpha BA \right]}{\left[ E\alpha A \right]\left[ B \right]}\to\left[ E\alpha BA \right]=K_{5}\left[ E\alpha A \right]\left[ B \right]\to\left[ E\alpha BA \right]=K_{1}K_{2}K_{5}\left[ E \right]\left[ \alpha\right]\left[ A \right]\left[ B \right]$$

$$consiquently: \therefore K_{2}K_{5}=K_{3}K_{4}$$

$$v=k_{6}\left[ E\alpha BA \right]$$

$$v=k_{6}K_{1}K_{3}K_{4}\left[ E \right]\left[ \alpha\right]\left[ B \right]\left[ A \right]$$

$$v=\left( k_{6}K_{1}K_{3}K_{4}\left[ \alpha\right]\left[ B \right]\left[ A \right] \right)\left[ E \right]$$

The fraction of uncomplexed enzyme is difficult to experimentally determine, and therefore the total enzyme concentration will be substituted in:

$$\left[ E_{t} \right]=\left[ E \right]+\left[ E\alpha\right]+\left[ E\alpha A \right]+\left[ E\alpha B \right]+\left[ E\alpha BA \right]$$

$$\left[ E_{t} \right]=\left[ E \right]+K_{1}\left[ E \right]\left[ \alpha\right]+K_{1}K_{2}\left[ E \right]\left[ \alpha\right]\left[ A \right]+K_{1}K_{3}\left[ E \right]\left[ \alpha\right]\left[ B \right]+K_{1}K_{3}K_{4}\left[ E \right]\left[ \alpha\right]\left[ B \right]\left[ A \right]$$

$$\left[ E_{t} \right]=\left[ E \right]\left( 1+K_{1}\left[ \alpha\right]+K_{1}K_{2}\left[ \alpha\right]\left[ A \right]+K_{1}K_{3}\left[ \alpha\right]\left[ B \right]+K_{1}K_{3}K_{4}\left[ B \right]\left[ A \right]\left[ \alpha\right] \right)$$

$$\left[ E_{t} \right]=\left[ E \right]\left( 1+K_{1}\left[ \alpha\right]\left( 1+K_{2}\left[ A \right]+K_{3}\left[ B \right]+K_{3}K_{4}\left[ B \right]\left[ A \right] \right) \right)$$

$$\left[ E \right]=\frac{\left[ E_{t} \right]}{1+K_{1}\left[ \alpha\right]\left( 1+K_{2}\left[ A \right]+K_{3}\left[ B \right]+K_{3}K_{4}\left[ B \right]\left[ A \right] \right)}$$

Finally, solving for the reaction velocity:

$$v=\left( k_{6}K_{1}K_{3}K_{4}\left[ \alpha\right]\left[ B \right]\left[ A \right] \right)\left[ E \right]$$

$$v=\left( k_{6}K_{1}K_{3}K_{4}\left[ \alpha\right]\left[ B \right]\left[ A \right] \right)\frac{\left[ E_{t} \right]}{1+K_{1}\left[ \alpha\right]\left( 1+K_{2}\left[ A \right]+K_{3}\left[ B \right]+K_{3}K_{4}\left[ B \right]\left[ A \right] \right)}$$

$$v=\frac{k_{6}K_{1}K_{3}K_{4}\left[ E_{t} \right]\left[ \alpha\right]\left[ B \right]\left[ A \right]}{1+K_{1}\left[ \alpha\right]\left( 1+K_{2}\left[ A \right]+K_{3}\left[ B \right]+K_{3}K_{4}\left[ B \right]\left[ A \right] \right)}$$

$$v\boldsymbol{=}\frac{\boldsymbol{k}_{\boldsymbol{cat}}\left[ \boldsymbol{E}_{\boldsymbol{t}} \right]\left[ \boldsymbol{\alpha} \right]\left[ \boldsymbol{B} \right]\left[ \boldsymbol{A} \right]}{\boldsymbol{1+}\boldsymbol{K}_{\boldsymbol{\alpha}}\left[ \boldsymbol{\alpha} \right]\left( \boldsymbol{1+}\boldsymbol{K}_{\boldsymbol{A\alpha}}\left[ \boldsymbol{A} \right]\boldsymbol{+}\boldsymbol{K}_{\boldsymbol{B\alpha}}\left[ \boldsymbol{B} \right]\boldsymbol{+}\boldsymbol{K}_{\boldsymbol{AB\alpha}}\left[ \boldsymbol{B} \right]\left[ \boldsymbol{A} \right] \right)}$$

$$\boldsymbol{v=}\frac{\boldsymbol{V}_{\boldsymbol{max}}\left[ \boldsymbol{\alpha} \right]\left[ \boldsymbol{B} \right]\left[ \boldsymbol{A} \right]}{\boldsymbol{1+}\boldsymbol{K}_{\boldsymbol{\alpha}}\left[ \boldsymbol{\alpha} \right]\left( \boldsymbol{1+}\boldsymbol{K}_{\boldsymbol{A\alpha}}\left[ \boldsymbol{A} \right]\boldsymbol{+}\boldsymbol{K}_{\boldsymbol{B\alpha}}\left[ \boldsymbol{B} \right]\boldsymbol{+}\boldsymbol{K}_{\boldsymbol{AB\alpha}}\left[ \boldsymbol{B} \right]\left[ \boldsymbol{A} \right] \right)}$$

$$\boldsymbol{parameters:} \boldsymbol{V}_{\boldsymbol{max}}\boldsymbol{,}\left[ \boldsymbol{\alpha} \right]\boldsymbol{,}\left[ \boldsymbol{B} \right]\boldsymbol{,}\left[ \boldsymbol{A} \right]$$

$$\boldsymbol{variables:} \boldsymbol{K}_{\boldsymbol{\alpha}}\boldsymbol{,}\boldsymbol{K}_{\boldsymbol{A\alpha}}\boldsymbol{,}\boldsymbol{K}_{\boldsymbol{B\alpha}}\boldsymbol{,}\boldsymbol{K}_{\boldsymbol{AB\alpha}}$$

#### **2.4 Reversible Dual Substrate**

Below are derivations or statements of kinetic equations for reversible dual-substrate reactions

##### **2.4.1 Reversible Dual-Substrate Kinetics**

Reversible dual-substrate Michealis-Menten like kinetics

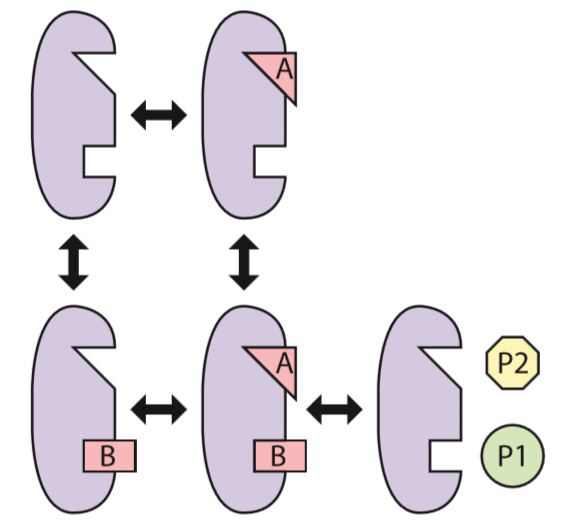

As an extension of the PSSH as applied to reaction 1, 2, 3, and 4 can be used to prove that all four of these reactions are at equilibrium, and therefore the equilibrium equations will be used for the derivation of kinetics. The equilibrium equations and the reaction velocity equation are given below:

$$K_{1}=\frac{\left[ EA \right]}{\left[ E \right]\left[ A \right]}\to\left[ EA \right]=K_{1}\left[ E \right]\left[ A \right]$$

$$K_{2}=\frac{\left[ EB \right]}{\left[ E \right]\left[ B \right]}\to\left[ EB \right]=K_{2}\left[ E \right]\left[ B \right]$$

$$K_{3}=\frac{\left[ EBA \right]}{\left[ EA \right]\left[ B \right]}\to\left[ EAB \right]=K_{3}\left[ EA \right]\left[ B \right]\to\left[ EAB \right]=K_{1}K_{3}\left[ E \right]\left[ A \right]\left[ B \right]$$

$$K_{4}=\frac{\left[ EBA \right]}{\left[ EB \right]\left[ A \right]}\to\left[ EAB \right]=K_{4}\left[ EB \right]\left[ A \right]\to\left[ EAB \right]=K_{2}K_{4}\left[ E \right]\left[ A \right]\left[ B \right]$$

$$\therefore K_{2}K_{4}=K_{1}K_{3}$$

$$v=k_{5}\left[ EAB \right]-k_{5}^{'}\left[ E \right]\left[ P_{1} \right]\ldots\left[ P_{N} \right]$$

$$v=k_{5}K_{1}K_{3}\left[ E \right]\left[ A \right]\left[ B \right]-k_{5}^{'}\left[ E \right]\left[ P_{1} \right]\ldots\left[ P_{N} \right]$$

$$v=\left( k_{5}K_{1}K_{3}\left[ A \right]\left[ B \right]-k_{5}^{'}\left[ P_{1} \right]\ldots\left[ P_{N} \right] \right)\left[ E \right]$$

Since uncomplexed enzyme concentration is difficult to measure, the total enzyme concentration is substituted:

$$\left[ E_{t} \right]=\left[ E \right]+\left[ EA \right]+\left[ EB \right]+[EAB]$$

$$\left[ E_{t} \right]=\left[ E \right]+K_{1}\left[ E \right]\left[ A \right]+K_{2}\left[ E \right]\left[ B \right]+K_{1}K_{3}\left[ E \right]\left[ A \right]\left[ B \right]$$

$$\left[ E_{t} \right]=\left[ E \right]\left( 1+K_{1}\left[ A \right]+K_{2}\left[ B \right]+K_{1}K_{3}\left[ A \right]\left[ B \right] \right)$$

$$\left[ E \right]=\frac{\left[ E_{t} \right]}{1+K_{1}\left[ A \right]+K_{2}\left[ B \right]+K_{1}K_{3}\left[ A \right]\left[ B \right]}$$

Making the above substitution into the reaction velocity equation

$$v=\left( k_{5}K_{1}K_{3}\left[ A \right]\left[ B \right]-k_{5}^{'}\left[ P_{1} \right]\ldots\left[ P_{N} \right] \right)\left[ E \right]$$

$$v=\left( k_{5}K_{1}K_{3}\left[ A \right]\left[ B \right]-k_{5}^{'}\left[ P_{1} \right]\ldots\left[ P_{N} \right] \right)\frac{\left[ E_{t} \right]}{1+K_{1}\left[ A \right]+K_{2}\left[ B \right]+K_{1}K_{3}\left[ A \right]\left[ B \right]}$$

$$v=\frac{\left( k_{5}K_{1}K_{3}\left[ A \right]\left[ B \right]-k_{5}^{'}\left[ P_{1} \right]\ldots\left[ P_{N} \right] \right)\left[ E_{t} \right]}{1+K_{1}\left[ A \right]+K_{2}\left[ B \right]+K_{1}K_{3}\left[ A \right]\left[ B \right]}$$

$$v=\frac{k_{5}K_{1}K_{3}\left[ E_{t} \right]\left[ A \right]\left[ B \right]-k_{5}^{'}\left[ E_{t} \right]\left[ P_{1} \right]\ldots\left[ P_{N} \right]}{1+K_{1}\left[ A \right]+K_{2}\left[ B \right]+K_{1}K_{3}\left[ A \right]\left[ B \right]}$$

$$\boldsymbol{v=}\frac{\boldsymbol{k}_{\boldsymbol{cat,f}}\left[ \boldsymbol{E}_{\boldsymbol{t}} \right]\left[ \boldsymbol{A} \right]\left[ \boldsymbol{B} \right]\boldsymbol{-}\boldsymbol{k}_{\boldsymbol{cat,b}}\left[ \boldsymbol{E}_{\boldsymbol{t}} \right]\left[ \boldsymbol{P}_{\boldsymbol{1}} \right]\boldsymbol{\ldots}\left[ \boldsymbol{P}_{\boldsymbol{N}} \right]}{\boldsymbol{1+}\boldsymbol{K}_{\boldsymbol{1}}\left[ \boldsymbol{A} \right]\boldsymbol{+}\boldsymbol{K}_{\boldsymbol{2}}\left[ \boldsymbol{B} \right]\boldsymbol{+}\boldsymbol{K}_{\boldsymbol{1}}\boldsymbol{K}_{\boldsymbol{3}}\left[ \boldsymbol{A} \right]\left[ \boldsymbol{B} \right]}$$

$$\boldsymbol{v=}\frac{\boldsymbol{V}_{\boldsymbol{max,f}}\left[ \boldsymbol{A} \right]\left[ \boldsymbol{B} \right]\boldsymbol{-}\boldsymbol{V}_{\boldsymbol{max,b}}\left[ \boldsymbol{P}_{\boldsymbol{1}} \right]\boldsymbol{\ldots}\left[ \boldsymbol{P}_{\boldsymbol{N}} \right]}{\boldsymbol{1+}\boldsymbol{K}_{\boldsymbol{A}}\left[ \boldsymbol{A} \right]\boldsymbol{+}\boldsymbol{K}_{\boldsymbol{B}}\left[ \boldsymbol{B} \right]\boldsymbol{+}\boldsymbol{K}_{\boldsymbol{AB}}\left[ \boldsymbol{A} \right]\left[ \boldsymbol{B} \right]}$$

$$\boldsymbol{parameters:}\boldsymbol{V}_{\boldsymbol{max,f}}\boldsymbol{,}\left[ \boldsymbol{A} \right]\boldsymbol{,}\left[ \boldsymbol{B} \right]\boldsymbol{,}\boldsymbol{V}_{\boldsymbol{max,b}}\boldsymbol{,}\left[ \boldsymbol{P}_{\boldsymbol{1}} \right]\boldsymbol{,\ldots,}\left[ \boldsymbol{P}_{\boldsymbol{N}} \right]$$

$$\boldsymbol{variables:}\boldsymbol{K}_{\boldsymbol{A}}\boldsymbol{,}\boldsymbol{K}_{\boldsymbol{B}}\boldsymbol{,}\boldsymbol{K}_{\boldsymbol{AB}}$$

**2.4.2 Reversible Dual-Substrate Allosteric Inhibition Kinetics**

Assuming reversible and unspecific ordering of binding of both substrates. For simplicity of derivation, it will be assumed that the allosteric inhibitor deforms the active sites and prevents binding (substrates and inhibitor are mutually exclusive). Further, inhibition binding is not permitted when a substrate is bound.

The elementary reactions are:

$$E+I\leftrightharpoons EI,K_{1}$$

$$E+A\leftrightharpoons EA, K_{2}$$

$$E+B\leftrightharpoons EB,K_{3}$$

$$EA+B\leftrightharpoons EAB,K_{4}$$

$$EB+A\leftrightharpoons EAB, K_{5}$$

$$EAB\leftrightharpoons E+P_{1}+\ldots+P_{N},K_{6}$$

Using the equilibrium constants gives the equations:

$$K_{1}=\frac{[EI]}{\left[ E \right][I]}$$

$$K_{2}=\frac{[EA]}{\left[ E \right][A]}$$

$$K_{3}=\frac{[EB]}{\left[ E \right][B]}$$

$$K_{4}=\frac{[EAB]}{\left[ EA \right][B]}$$

$$K_{5}=\frac{[EAB]}{\left[ EB \right][A]}$$

$$v=k_{6}\left[ EAB \right]-k_{6}^{'}\left[ E \right]\left[ P_{1} \right]\ldots\left[ P_{N} \right]$$

Solving these equations for unknown enzyme concentrations and for the reaction velocity gives:

$$K_{1}=\frac{[EI]}{\left[ E \right][I]}\to\left[ EI \right]=K_{1}\left[ E \right]\left[ I \right]$$

$$K_{2}=\frac{[EA]}{\left[ E \right][A]}\to\left[ EA \right]=K_{2}\left[ E \right][A]$$

$$K_{3}=\frac{[EB]}{\left[ E \right][B]}\to\left[ EB \right]=K_{3}\left[ E \right][B]$$

$$K_{4}=\frac{[EAB]}{\left[ EA \right][B]}\to\left[ EAB \right]=K_{4}\left[ EA \right]\left[ B \right]\to\left[ EAB \right]=K_{4}K_{2}\left[ E \right][A]\left[ B \right]$$

$$K_{5}=\frac{[EAB]}{\left[ EB \right][A]}\to\left[ EAB \right]=K_{5}\left[ EB \right]\left[ A \right]\to\left[ EAB \right]=K_{5}K_{3}\left[ E \right]\left[ A \right][B]$$

$$\therefore K_{4}K_{2}=K_{5}K_{3}$$

It must be assumed that the final reaction is not at equilibrium (otherwise $v$ will become zero, independent of any variable or parameter). Since it is difficult to measure uncomplexed enzyme concentration, substitute the total enzyme concentration:

$$\left[ E_{t} \right]=\left[ E \right]+\left[ EA \right]+\left[ EI \right]+\left[ EB \right]+\left[ EAB \right]$$

$$\left[ E_{t} \right]=\left[ E \right]+K_{2}\left[ E \right][A]+K_{1}\left[ E \right]\left[ I \right]+K_{3}\left[ E \right][B]+K_{4}K_{2}\left[ E \right][A]\left[ B \right]$$

$$\left[ E_{t} \right]=\left( 1+K_{2}[A]+K_{1}\left[ I \right]+K_{3}[B]+K_{4}K_{2}[A]\left[ B \right] \right)\left[ E \right]$$

$$\left[ E \right]=\frac{\left[ E_{t} \right]}{1+K_{2}[A]+K_{1}\left[ I \right]+K_{3}[B]+K_{4}K_{2}[A]\left[ B \right]}$$

$$v=k_{6}\left[ EAB \right]-k_{6}^{'}\left[ E \right]\left[ P_{1} \right]\ldots\left[ P_{N} \right]$$

$$v=\left( k_{6}K_{4}K_{2}[A]\left[ B \right]-k_{6}^{'}\left[ P_{1} \right]\ldots\left[ P_{N} \right] \right)\left[ E \right]$$

$$v=\left( k_{6}K_{4}K_{2}[A]\left[ B \right]-k_{6}^{'}\left[ P_{1} \right]\ldots\left[ P_{N} \right] \right)\frac{\left[ E_{t} \right]}{1+K_{2}[A]+K_{1}\left[ I \right]+K_{3}[B]+K_{4}K_{2}[A]\left[ B \right]}$$

$$v=\frac{\left[ E_{t} \right]\left( k_{6}K_{4}K_{2}[A]\left[ B \right]-k_{6}^{'}\left[ P_{1} \right]\ldots\left[ P_{N} \right] \right)}{1+K_{2}[A]+K_{1}\left[ I \right]+K_{3}[B]+K_{4}K_{2}[A]\left[ B \right]}$$

$$v=\frac{k_{6}K_{4}K_{2}\left[ E_{t} \right][A]\left[ B \right]-k_{6}^{'}\left[ E_{t} \right]\left[ P_{1} \right]\ldots\left[ P_{N} \right]}{1+K_{2}[A]+K_{1}\left[ I \right]+K_{3}[B]+K_{4}K_{2}[A]\left[ B \right]}$$

$$\boldsymbol{v=}\frac{\boldsymbol{k}_{\boldsymbol{6}}\boldsymbol{K}_{\boldsymbol{4}}\boldsymbol{K}_{\boldsymbol{2}}\left[ \boldsymbol{E}_{\boldsymbol{t}} \right]\boldsymbol{[A]}\left[ \boldsymbol{B} \right]\boldsymbol{-}\boldsymbol{k}_{\boldsymbol{6}}^{\boldsymbol{'}}\left[ \boldsymbol{E}_{\boldsymbol{t}} \right]\left[ \boldsymbol{P}_{\boldsymbol{1}} \right]\boldsymbol{\ldots}\left[ \boldsymbol{P}_{\boldsymbol{N}} \right]}{\boldsymbol{1+}\boldsymbol{K}_{\boldsymbol{A}}\boldsymbol{[A]+}\boldsymbol{K}_{\boldsymbol{I}}\left[ \boldsymbol{I} \right]\boldsymbol{+}\boldsymbol{K}_{\boldsymbol{B}}\boldsymbol{[B]+}\boldsymbol{K}_{\boldsymbol{AB}}\boldsymbol{[A]}\left[ \boldsymbol{B} \right]}\boldsymbol{=}\frac{\boldsymbol{V}_{\boldsymbol{max,f}}\boldsymbol{[A]}\left[ \boldsymbol{B} \right]\boldsymbol{-}\boldsymbol{V}_{\boldsymbol{max,b}}\left[ \boldsymbol{P}_{\boldsymbol{1}} \right]\boldsymbol{\ldots}\left[ \boldsymbol{P}_{\boldsymbol{N}} \right]}{\boldsymbol{1+}\boldsymbol{K}_{\boldsymbol{A}}\boldsymbol{[A]+}\boldsymbol{K}_{\boldsymbol{I}}\left[ \boldsymbol{I} \right]\boldsymbol{+}\boldsymbol{K}_{\boldsymbol{B}}\boldsymbol{[B]+}\boldsymbol{K}_{\boldsymbol{AB}}\boldsymbol{[A]}\left[ \boldsymbol{B} \right]}$$

$$\boldsymbol{parameters:}\boldsymbol{V}_{\boldsymbol{max,f}}\boldsymbol{,}\boldsymbol{V}_{\boldsymbol{max,b}}\boldsymbol{,}\left[ \boldsymbol{A} \right]\boldsymbol{,}\left[ \boldsymbol{B} \right]\boldsymbol{,}\left[ \boldsymbol{I} \right]\boldsymbol{,}\left[ \boldsymbol{P}_{\boldsymbol{1}} \right]\boldsymbol{,\ldots,}\left[ \boldsymbol{P}_{\boldsymbol{N}} \right]$$

$$\boldsymbol{variables:}\boldsymbol{K}_{\boldsymbol{A}}\boldsymbol{,}\boldsymbol{K}_{\boldsymbol{B}}\boldsymbol{,}\boldsymbol{K}_{\boldsymbol{I}}\boldsymbol{,}\boldsymbol{K}_{\boldsymbol{AB}}\boldsymbol{,}\boldsymbol{K}_{\boldsymbol{ABI}}$$

##### **2.4.3 Reversible Dual-Substrate Allosteric Activation Kinetics**

Assumes that activation is a prerequisite for substrate binding, then allows non-specific substrate binding order (to avoid having to derive an equation for each case). Once the activator is bound, it is assumed to remain bound until the enzyme is bound to nothing else.

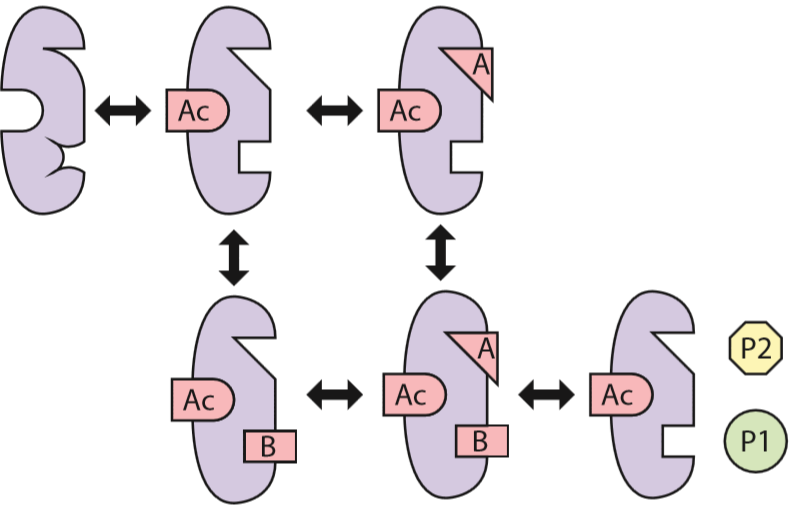

Here we will again use equilibrium constants for all reactions except the product-forming one. This is essentially an extension of the PSSH as applied to equilibrium constants as opposed to reaction rates (that this is an extension of PSSH can be easily proven). Therefore the equilibrium equations and reaction velocity may be given as:

$$K_{1}=\frac{[E\alpha]}{\left[ E \right][\alpha]}\to\left[ E\alpha\right]=K_{1}\left[ E \right]\left[ \alpha\right]$$

$$K_{2}=\frac{\left[ E\alpha A \right]}{\left[ E\alpha\right]\left[ A \right]}\to\left[ E\alpha A \right]=K_{2}\left[ E\alpha\right]\left[ A \right]\to\left[ E\alpha A \right]=K_{1}K_{2}\left[ E \right]\left[ \alpha\right]\left[ A \right]$$

$$K_{3}=\frac{\left[ E\alpha B \right]}{\left[ E\alpha\right]\left[ B \right]}\to\left[ E\alpha B \right]=K_{3}\left[ E\alpha\right]\left[ B \right]\to\left[ E\alpha B \right]=K_{1}K_{3}\left[ E \right]\left[ \alpha\right]\left[ B \right]$$

$$K_{4}=\frac{\left[ E\alpha BA \right]}{\left[ E\alpha B \right]\left[ A \right]}\to\left[ E\alpha BA \right]=K_{4}\left[ E\alpha B \right][A]\to\left[ E\alpha BA \right]=K_{1}K_{3}K_{4}\left[ E \right]\left[ \alpha\right]\left[ B \right]\left[ A \right]$$

$$K_{5}=\frac{\left[ E\alpha BA \right]}{\left[ E\alpha A \right]\left[ B \right]}\to\left[ E\alpha BA \right]=K_{5}\left[ E\alpha A \right]\left[ B \right]\to\left[ E\alpha BA \right]=K_{1}K_{2}K_{5}\left[ E \right]\left[ \alpha\right]\left[ A \right]\left[ B \right]$$

$$consiquently: \therefore K_{2}K_{5}=K_{3}K_{4}$$

$$v=k_{6}\left[ E\alpha BA \right]-k_{6}^{'}\left[ E\alpha\right]\left[ P_{1} \right]\ldots\left[ P_{N} \right]$$

$$v=k_{6}K_{1}K_{3}K_{4}\left[ E \right]\left[ \alpha\right]\left[ B \right]\left[ A \right]-k_{6}^{'}K_{1}\left[ E \right]\left[ \alpha\right]\left[ P_{1} \right]\ldots\left[ P_{N} \right]$$

$$v=\left( k_{6}K_{1}K_{3}K_{4}\left[ \alpha\right]\left[ B \right]\left[ A \right]-k_{6}^{'}K_{1}\left[ \alpha\right]\left[ P_{1} \right]\ldots\left[ P_{N} \right] \right)\left[ E \right]$$

Since the concentration of uncomplexed enzymes is difficult to measure, the total enzyme concentration will be used:

$$\left[ E_{t} \right]=\left[ E \right]+\left[ E\alpha\right]+\left[ E\alpha A \right]+\left[ E\alpha B \right]+\left[ E\alpha BA \right]$$

$$\left[ E_{t} \right]=\left[ E \right]+K_{1}\left[ E \right]\left[ \alpha\right]+K_{1}K_{2}\left[ E \right]\left[ \alpha\right]\left[ A \right]+K_{1}K_{3}\left[ E \right]\left[ \alpha\right]\left[ B \right]+K_{1}K_{3}K_{4}\left[ E \right]\left[ \alpha\right]\left[ B \right]\left[ A \right]$$

$$\left[ E_{t} \right]=\left[ E \right]\left( 1+K_{1}\left[ \alpha\right]+K_{1}K_{2}\left[ \alpha\right]\left[ A \right]+K_{1}K_{3}\left[ \alpha\right]\left[ B \right]+K_{1}K_{3}K_{4}\left[ B \right]\left[ A \right]\left[ \alpha\right] \right)$$

$$\left[ E_{t} \right]=\left[ E \right]\left( 1+K_{1}\left[ \alpha\right]\left( 1+K_{2}\left[ A \right]+K_{3}\left[ B \right]+K_{3}K_{4}\left[ B \right]\left[ A \right] \right) \right)$$

$$\left[ E \right]=\frac{\left[ E_{t} \right]}{1+K_{1}\left[ \alpha\right]\left( 1+K_{2}\left[ A \right]+K_{3}\left[ B \right]+K_{3}K_{4}\left[ B \right]\left[ A \right] \right)}$$

$$v=\left( k_{6}K_{1}K_{3}K_{4}\left[ \alpha\right]\left[ B \right]\left[ A \right]-k_{6}^{'}K_{1}\left[ \alpha\right]\left[ P_{1} \right]\ldots\left[ P_{N} \right] \right)\left[ E \right]$$

$$v=\left( k_{6}K_{1}K_{3}K_{4}\left[ \alpha\right]\left[ B \right]\left[ A \right]-k_{6}^{'}K_{1}\left[ \alpha\right]\left[ P_{1} \right]\ldots\left[ P_{N} \right] \right)\frac{\left[ E_{t} \right]}{1+K_{1}\left[ \alpha\right]\left( 1+K_{2}\left[ A \right]+K_{3}\left[ B \right]+K_{3}K_{4}\left[ B \right]\left[ A \right] \right)}$$

$$v=\frac{\left[ E_{t} \right]\left( k_{6}K_{1}K_{3}K_{4}\left[ \alpha\right]\left[ B \right]\left[ A \right]-k_{6}^{'}K_{1}\left[ \alpha\right]\left[ P_{1} \right]\ldots\left[ P_{N} \right] \right)}{1+K_{1}\left[ \alpha\right]\left( 1+K_{2}\left[ A \right]+K_{3}\left[ B \right]+K_{3}K_{4}\left[ B \right]\left[ A \right] \right)}$$

$$v=\frac{k_{6}K_{1}K_{3}K_{4}\left[ \alpha\right]\left[ B \right]\left[ A \right]\left[ E_{t} \right]-k_{6}^{'}K_{1}\left[ E_{t} \right]\left[ \alpha\right]\left[ P_{1} \right]\ldots\left[ P_{N} \right]}{1+K_{1}\left[ \alpha\right]\left( 1+K_{2}\left[ A \right]+K_{3}\left[ B \right]+K_{3}K_{4}\left[ B \right]\left[ A \right] \right)}$$

$$\boldsymbol{v=}\frac{\boldsymbol{k}_{\boldsymbol{cat,f}}\left[ \boldsymbol{E}_{\boldsymbol{t}} \right]\left[ \boldsymbol{\alpha} \right]\left[ \boldsymbol{B} \right]\left[ \boldsymbol{A} \right]\boldsymbol{-}\boldsymbol{k}_{\boldsymbol{cat,b}}\left[ \boldsymbol{E}_{\boldsymbol{t}} \right]\left[ \boldsymbol{\alpha} \right]\left[ \boldsymbol{P}_{\boldsymbol{1}} \right]\boldsymbol{\ldots}\left[ \boldsymbol{P}_{\boldsymbol{N}} \right]}{\boldsymbol{1+}\boldsymbol{K}_{\boldsymbol{\alpha}}\left[ \boldsymbol{\alpha} \right]\left( \boldsymbol{1+}\boldsymbol{K}_{\boldsymbol{A\alpha}}\left[ \boldsymbol{A} \right]\boldsymbol{+}\boldsymbol{K}_{\boldsymbol{B\alpha}}\left[ \boldsymbol{B} \right]\boldsymbol{+}\boldsymbol{K}_{\boldsymbol{AB\alpha}}\left[ \boldsymbol{B} \right]\left[ \boldsymbol{A} \right] \right)}$$

$$\boldsymbol{v=}\frac{\boldsymbol{V}_{\boldsymbol{max,f}}\left[ \boldsymbol{E}_{\boldsymbol{t}} \right]\left[ \boldsymbol{\alpha} \right]\left[ \boldsymbol{B} \right]\left[ \boldsymbol{A} \right]\boldsymbol{-}\boldsymbol{V}_{\boldsymbol{max,b}}\left[ \boldsymbol{E}_{\boldsymbol{t}} \right]\left[ \boldsymbol{\alpha} \right]\left[ \boldsymbol{P}_{\boldsymbol{1}} \right]\boldsymbol{\ldots}\left[ \boldsymbol{P}_{\boldsymbol{N}} \right]}{\boldsymbol{1+}\boldsymbol{K}_{\boldsymbol{\alpha}}\left[ \boldsymbol{\alpha} \right]\left( \boldsymbol{1+}\boldsymbol{K}_{\boldsymbol{A\alpha}}\left[ \boldsymbol{A} \right]\boldsymbol{+}\boldsymbol{K}_{\boldsymbol{B\alpha}}\left[ \boldsymbol{B} \right]\boldsymbol{+}\boldsymbol{K}_{\boldsymbol{AB\alpha}}\left[ \boldsymbol{B} \right]\left[ \boldsymbol{A} \right] \right)}$$

$$\boldsymbol{parameters:}\left[ \boldsymbol{E}_{\boldsymbol{t}} \right]\boldsymbol{,}\left[ \boldsymbol{\alpha} \right]\boldsymbol{,}\left[ \boldsymbol{B} \right]\boldsymbol{,}\left[ \boldsymbol{A} \right]\boldsymbol{,}\boldsymbol{V}_{\boldsymbol{max,b}}\boldsymbol{,}\boldsymbol{V}_{\boldsymbol{max,f}}\boldsymbol{,}\left[ \boldsymbol{P}_{\boldsymbol{1}} \right]\boldsymbol{,\ldots,}\left[ \boldsymbol{P}_{\boldsymbol{N}} \right]$$

$$\boldsymbol{variables:} \boldsymbol{K}_{\boldsymbol{AB\alpha}}\boldsymbol{,}\boldsymbol{K}_{\boldsymbol{B\alpha}}\boldsymbol{,}\boldsymbol{K}_{\boldsymbol{A\alpha}}\boldsymbol{,}\boldsymbol{K}_{\boldsymbol{\alpha}}$$

#### **2.5 Irreversible Triple Substrate Kinetics**

Reaction kinetics for all three triple substrate kinetic forms is shown below.

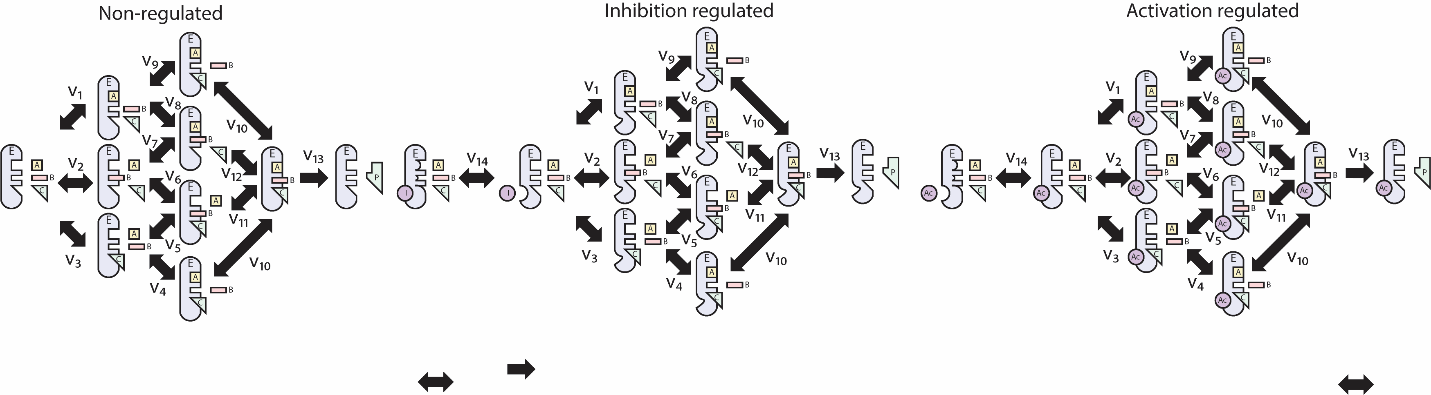

##### **2.5.1 Irreversible Triple Substrate Kinetics**

Assuming the final product-forming step is irreversible and the intermediate steps are reversible, and that the binding order of metabolites is not important. We will assume the pseudo-steady state hypothesis in order to derive the kinetic equation. Equilibrium constants will be used for each reaction:

$$K_{1}=\frac{\left[ EA \right]}{\left[ E \right]\left[ A \right]}\to\left[ EA \right]=K_{1}\left[ E \right]\left[ A \right]$$

$$K_{2}=\frac{\left[ EB \right]}{\left[ E \right]\left[ B \right]}\to\left[ EB \right]=K_{2}\left[ E \right]\left[ B \right]$$

$$K_{3}=\frac{\left[ EC \right]}{\left[ E \right]\left[ C \right]}\to\left[ EC \right]=K_{3}\left[ E \right]\left[ C \right]$$

$$K_{4}=\frac{\left[ EAC \right]}{\left[ EC \right]\left[ A \right]}\to\left[ EAC \right]=K_{4}\left[ EC \right]\left[ A \right]\to\left[ EAC \right]=K_{3}K_{4}\left[ E \right]\left[ A \right]\left[ C \right]$$

$$K_{5}=\frac{\left[ EBC \right]}{\left[ EC \right]\left[ B \right]}\to\left[ EBC \right]=K_{5}\left[ EC \right]\left[ B \right]\to\left[ EBC \right]=K_{3}K_{5}\left[ E \right]\left[ B \right]\left[ C \right]$$

$$K_{6}=\frac{\left[ EBC \right]}{\left[ EB \right]\left[ C \right]}\to\left[ EBC \right]=K_{6}\left[ EB \right]\left[ C \right]\to\left[ EBC \right]=K_{2}K_{6}\left[ E \right]\left[ B \right]\left[ C \right]$$

$$\therefore K_{2}K_{6}=K_{3}K_{5}$$

$$K_{7}=\frac{\left[ EAB \right]}{\left[ EB \right]\left[ A \right]}\to\left[ EAB \right]=K_{7}\left[ EB \right]\left[ A \right]\to K_{2}K_{7}\left[ E \right]\left[ A \right]\left[ B \right]$$

$$K_{8}=\frac{\left[ EAB \right]}{\left[ EA \right]\left[ B \right]}\to\left[ EAB \right]=K_{8}\left[ EA \right]\left[ B \right]\to K_{1}K_{8}\left[ E \right]\left[ A \right]\left[ B \right]$$

$$\therefore K_{1}K_{8}=K_{2}K_{7}$$

$$K_{9}=\frac{\left[ EAC \right]}{\left[ EA \right]\left[ C \right]}\to\left[ EAC \right]=K_{9}\left[ EA \right]\left[ C \right]\to\left[ EAC \right]=K_{1}K_{9}\left[ E \right]\left[ A \right]\left[ C \right]$$

$$\therefore K_{1}K_{9}=K_{3}K_{4}$$

$$K_{10}=\frac{\left[ EABC \right]}{\left[ EAC \right]\left[ B \right]}\to\left[ EABC \right]=K_{10}\left[ EAC \right]\left[ B \right]\to\left[ EABC \right]=K_{1}K_{9}K_{10}\left[ E \right]\left[ A \right]\left[ B \right]\left[ C \right]$$

$$K_{11}=\frac{\left[ EABC \right]}{\left[ EBC \right]\left[ A \right]}\to\left[ EABC \right]=K_{11}\left[ EBC \right]\left[ A \right]\to\left[ EABC \right]=K_{2}K_{6}K_{11}\left[ E \right]\left[ A \right]\left[ B \right]\left[ C \right]$$

$$K_{12}=\frac{\left[ EABC \right]}{\left[ EAB \right]\left[ C \right]}\to\left[ EABC \right]=K_{12}\left[ EAB \right]\left[ C \right]\to\left[ EABC \right]=K_{1}K_{8}K_{12}\left[ E \right]\left[ A \right]\left[ B \right]\left[ C \right]$$

$$\therefore K_{1}K_{9}K_{10}=K_{2}K_{6}K_{11}=K_{1}K_{8}K_{12}$$

Then:

$$v_{total}=k_{13}\left[ EABC \right]=k_{13}K_{1}K_{8}K_{12}\left[ E \right]\left[ A \right]\left[ B \right]\left[ C \right]$$

Since the concentration of the enzyme complexed or uncomplexed is difficult to measure. Therefore, let us consider using the total enzyme concentration to solve for the concentration of uncomplexed enzyme concentration.

$$\left[ E \right]_{total}=\left[ E \right]+\left[ EA \right]+\left[ EB \right]+\left[ EC \right]+\left[ EAB \right]+\left[ EAC \right]+\left[ EBC \right]+\left[ EABC \right]$$

$$\left[ E \right]_{total}=\left[ E \right]+K_{1}\left[ E \right]\left[ A \right]+K_{2}\left[ E \right]\left[ B \right]+K_{3}\left[ E \right]\left[ C \right]+K_{1}K_{8}\left[ E \right]\left[ A \right]\left[ B \right]+K_{3}K_{4}\left[ E \right]\left[ A \right]\left[ C \right]+K_{3}K_{5}\left[ E \right]\left[ B \right]\left[ C \right]+K_{1}K_{8}K_{12}\left[ E \right]\left[ A \right]\left[ B \right]\left[ C \right]$$

$$\left[ E \right]_{total}=\left[ E \right]\left( 1+K_{1}\left[ A \right]+K_{2}\left[ B \right]+K_{3}\left[ C \right]+K_{1}K_{8}\left[ A \right]\left[ B \right]+K_{3}K_{4}\left[ A \right]\left[ C \right]+K_{3}K_{5}\left[ B \right]\left[ C \right]+K_{1}K_{8}K_{12}\left[ A \right]\left[ B \right]\left[ C \right] \right)$$

$$\left[ E \right]=\frac{\left[ E \right]_{total}}{1+K_{1}\left[ A \right]+K_{2}\left[ B \right]+K_{3}\left[ C \right]+K_{1}K_{8}\left[ A \right]\left[ B \right]+K_{3}K_{4}\left[ A \right]\left[ C \right]+K_{3}K_{5}\left[ B \right]\left[ C \right]+K_{1}K_{8}K_{12}\left[ A \right]\left[ B \right]\left[ C \right]}$$

Substituting into the reaction rate equation for the total reaction rate gives:

$$v_{total}=\frac{k_{13}K_{1}K_{8}K_{12}\left[ A \right]\left[ B \right]\left[ C \right]\left[ E \right]_{total}}{1+K_{1}\left[ A \right]+K_{2}\left[ B \right]+K_{3}\left[ C \right]+K_{1}K_{8}\left[ A \right]\left[ B \right]+K_{3}K_{4}\left[ A \right]\left[ C \right]+K_{3}K_{5}\left[ B \right]\left[ C \right]+K_{1}K_{8}K_{12}\left[ A \right]\left[ B \right]\left[ C \right]}$$

Lumping some kinetic parameters together and using a $V_{max}$ term gives:

$$v_{total}=\frac{V_{max}\left[ A \right]\left[ B \right]\left[ C \right]}{1+K_{1}\left[ A \right]+K_{2}\left[ B \right]+K_{3}\left[ C \right]+K_{13}\left[ A \right]\left[ B \right]+K_{14}\left[ A \right]\left[ C \right]+K_{15}\left[ B \right]\left[ C \right]+K_{16}\left[ A \right]\left[ B \right]\left[ C \right]}$$

$$V_{max}=k_{13}K_{1}K_{8}K_{12}\left[ E \right]_{total}$$

$$K_{1}K_{8}=K_{13}$$

$$K_{3}K_{4}=K_{14}$$

$$K_{3}K_{5}=K_{15}$$

$$K_{1}K_{8}K_{12}=K_{16}$$

Simplifying to use the smallest possible coefficient subscripts (ignoring previous definitions of $K_{x}$ terms) gives:

$$\boldsymbol{v}_{\boldsymbol{total}}\boldsymbol{=}\frac{\boldsymbol{K}_{\boldsymbol{1}}\left[ \boldsymbol{A} \right]\left[ \boldsymbol{B} \right]\left[ \boldsymbol{C} \right]}{\boldsymbol{1+}\boldsymbol{K}_{\boldsymbol{2}}\left[ \boldsymbol{A} \right]\boldsymbol{+}\boldsymbol{K}_{\boldsymbol{3}}\left[ \boldsymbol{B} \right]\boldsymbol{+}\boldsymbol{K}_{\boldsymbol{4}}\left[ \boldsymbol{C} \right]\boldsymbol{+}\boldsymbol{K}_{\boldsymbol{5}}\left[ \boldsymbol{A} \right]\left[ \boldsymbol{B} \right]\boldsymbol{+}\boldsymbol{K}_{\boldsymbol{6}}\left[ \boldsymbol{A} \right]\left[ \boldsymbol{C} \right]\boldsymbol{+}\boldsymbol{K}_{\boldsymbol{7}}\left[ \boldsymbol{B} \right]\left[ \boldsymbol{C} \right]\boldsymbol{+}\boldsymbol{K}_{\boldsymbol{8}}\left[ \boldsymbol{A} \right]\left[ \boldsymbol{B} \right]\left[ \boldsymbol{C} \right]}$$

##### **2.5.2 Irreversible Triple Substrate Kinetics with Inhibition**

Assuming the final product-forming step is irreversible and the intermediate steps are reversible, and that the binding order of metabolites is not important. The binding of the inhibitor is incomparable with the binding of any metabolite.

$$K_{1}=\frac{\left[ EA \right]}{\left[ E \right]\left[ A \right]}\to\left[ EA \right]=K_{1}\left[ E \right]\left[ A \right]$$

$$K_{2}=\frac{\left[ EB \right]}{\left[ E \right]\left[ B \right]}\to\left[ EB \right]=K_{2}\left[ E \right]\left[ B \right]$$

$$K_{3}=\frac{\left[ EC \right]}{\left[ E \right]\left[ C \right]}\to\left[ EC \right]=K_{3}\left[ E \right]\left[ C \right]$$

$$K_{4}=\frac{\left[ EAC \right]}{\left[ EC \right]\left[ A \right]}\to\left[ EAC \right]=K_{4}\left[ EC \right]\left[ A \right]\to\left[ EAC \right]=K_{3}K_{4}\left[ E \right]\left[ A \right]\left[ C \right]$$

$$K_{5}=\frac{\left[ EBC \right]}{\left[ EC \right]\left[ B \right]}\to\left[ EBC \right]=K_{5}\left[ EC \right]\left[ B \right]\to\left[ EBC \right]=K_{3}K_{5}\left[ E \right]\left[ B \right]\left[ C \right]$$

$$K_{6}=\frac{\left[ EBC \right]}{\left[ EB \right]\left[ C \right]}\to\left[ EBC \right]=K_{6}\left[ EB \right]\left[ C \right]\to\left[ EBC \right]=K_{2}K_{6}\left[ E \right]\left[ B \right]\left[ C \right]$$

$$\therefore K_{2}K_{6}=K_{3}K_{5}$$

$$K_{7}=\frac{\left[ EAB \right]}{\left[ EB \right]\left[ A \right]}\to\left[ EAB \right]=K_{7}\left[ EB \right]\left[ A \right]\to\left[ EAB \right]=K_{2}K_{7}\left[ E \right]\left[ A \right]\left[ B \right]$$

$$K_{8}=\frac{\left[ EAB \right]}{\left[ EA \right]\left[ B \right]}\to\left[ EAB \right]=K_{8}\left[ EA \right]\left[ B \right]\to\left[ EAB \right]=K_{1}K_{8}\left[ E \right]\left[ A \right]\left[ B \right]$$

$$\therefore K_{1}K_{8}=K_{2}K_{7}$$

$$K_{9}=\frac{\left[ EAC \right]}{\left[ EA \right]\left[ C \right]}\to\left[ EAC \right]=K_{9}\left[ EA \right]\left[ C \right]\to\left[ EAC \right]=K_{1}K_{9}\left[ E \right]\left[ A \right]\left[ C \right]$$

$$\therefore K_{1}K_{9}=K_{3}K_{4}$$

$$K_{10}=\frac{\left[ EABC \right]}{\left[ EAC \right]\left[ B \right]}\to\left[ EABC \right]=K_{10}\left[ EAC \right]\left[ B \right]\to\left[ EABC \right]=K_{1}K_{9}K_{10}\left[ E \right]\left[ A \right]\left[ B \right]\left[ C \right]$$

$$K_{11}=\frac{\left[ EABC \right]}{\left[ EBC \right]\left[ A \right]}\to\left[ EABC \right]=K_{11}\left[ EBC \right]\left[ A \right]\to\left[ EABC \right]=K_{2}K_{6}K_{11}\left[ E \right]\left[ A \right]\left[ B \right]\left[ C \right]$$

$$K_{12}=\frac{\left[ EABC \right]}{\left[ EAB \right]\left[ C \right]}\to\left[ EABC \right]=K_{12}\left[ EAB \right]\left[ C \right]\to\left[ EABC \right]=K_{1}K_{8}K_{12}\left[ E \right]\left[ A \right]\left[ B \right]\left[ C \right]$$

$$\therefore K_{1}K_{9}K_{10}=K_{2}K_{6}K_{11}=K_{1}K_{8}K_{12}$$

$$K_{14}=\frac{\left[ E \right]\left[ I \right]}{\left[ EI \right]}\to\left[ EI \right]=\frac{1}{K_{14}}\left[ E \right]\left[ I \right]$$

Then:

$$v_{total}=k_{13}\left[ EABC \right]$$

$$v_{total}=k_{13}K_{1}K_{8}K_{12}\left[ E \right]\left[ A \right]\left[ B \right]\left[ C \right]$$

Since the concentration of uncomplexed or complexed enzymes is difficult to measure, the total enzyme concentration must be used.

$$\left[ E \right]_{total}=\left[ E \right]+\left[ EA \right]+\left[ EB \right]+\left[ EC \right]+\left[ EI \right]+\left[ EAB \right]+\left[ EAC \right]+\left[ EBC \right]+\left[ EABC \right]$$

$$\left[ E \right]_{total}=\left[ E \right]+K_{1}\left[ E \right]\left[ A \right]+K_{2}\left[ E \right]\left[ B \right]+K_{3}\left[ E \right]\left[ C \right]+\frac{1}{K_{14}}\left[ E \right]\left[ I \right]+K_{1}K_{8}\left[ E \right]\left[ A \right]\left[ B \right]+K_{3}K_{4}\left[ E \right]\left[ A \right]\left[ C \right]+K_{2}K_{6}\left[ E \right]\left[ B \right]\left[ C \right]+K_{1}K_{8}K_{12}\left[ E \right]\left[ A \right]\left[ B \right]\left[ C \right]$$

$$\left[ E \right]_{total}=\left[ E \right]\left( 1+K_{1}\left[ A \right]+K_{2}\left[ B \right]+K_{3}\left[ C \right]+\frac{1}{K_{14}}\left[ I \right]+K_{1}K_{8}\left[ A \right]\left[ B \right]+K_{3}K_{4}\left[ A \right]\left[ C \right]+K_{2}K_{6}\left[ B \right]\left[ C \right]+K_{1}K_{8}K_{12}\left[ A \right]\left[ B \right]\left[ C \right] \right)$$

$$\left[ E \right]=\frac{\left[ E \right]_{total}}{1+K_{1}\left[ A \right]+K_{2}\left[ B \right]+K_{3}\left[ C \right]+\frac{1}{K_{14}}\left[ I \right]+K_{1}K_{8}\left[ A \right]\left[ B \right]+K_{3}K_{4}\left[ A \right]\left[ C \right]+K_{2}K_{6}\left[ B \right]\left[ C \right]+K_{1}K_{8}K_{12}\left[ A \right]\left[ B \right]\left[ C \right]}$$

Therefore, the reaction rate is given as:

$$v_{total}=\frac{k_{13}K_{1}K_{8}K_{12}\left[ E \right]_{total}\left[ A \right]\left[ B \right]\left[ C \right]}{1+K_{1}\left[ A \right]+K_{2}\left[ B \right]+K_{3}\left[ C \right]+\frac{1}{K_{14}}\left[ I \right]+K_{1}K_{8}\left[ A \right]\left[ B \right]+K_{3}K_{4}\left[ A \right]\left[ C \right]+K_{2}K_{6}\left[ B \right]\left[ C \right]+K_{1}K_{8}K_{12}\left[ A \right]\left[ B \right]\left[ C \right]}$$

Ignoring the subscripts previously defined and seeking to minimize the number of kinetic parameters, this gives:

$$\boldsymbol{v}_{\boldsymbol{total}}\boldsymbol{=}\frac{\boldsymbol{K}_{\boldsymbol{1}}\left[ \boldsymbol{A} \right]\left[ \boldsymbol{B} \right]\left[ \boldsymbol{C} \right]}{\boldsymbol{1+}\boldsymbol{K}_{\boldsymbol{2}}\left[ \boldsymbol{A} \right]\boldsymbol{+}\boldsymbol{K}_{\boldsymbol{3}}\left[ \boldsymbol{B} \right]\boldsymbol{+}\boldsymbol{K}_{\boldsymbol{4}}\left[ \boldsymbol{C} \right]\boldsymbol{+}\boldsymbol{K}_{\boldsymbol{5}}\left[ \boldsymbol{I} \right]\boldsymbol{+}\boldsymbol{K}_{\boldsymbol{6}}\left[ \boldsymbol{A} \right]\left[ \boldsymbol{B} \right]\boldsymbol{+}\boldsymbol{K}_{\boldsymbol{7}}\left[ \boldsymbol{A} \right]\left[ \boldsymbol{C} \right]\boldsymbol{+}\boldsymbol{K}_{\boldsymbol{8}}\left[ \boldsymbol{B} \right]\left[ \boldsymbol{C} \right]\boldsymbol{+}\boldsymbol{K}_{\boldsymbol{9}}\left[ \boldsymbol{A} \right]\left[ \boldsymbol{B} \right]\left[ \boldsymbol{C} \right]}$$

$$K_{1}=k_{13}K_{1}K_{8}K_{12}\left[ E \right]_{total}$$

$$K_{5}=\frac{1}{K_{14}}$$

$$K_{6}=K_{1}K_{8}$$

$$K_{7}=K_{3}K_{4}$$

$$K_{8}=K_{2}K_{6}$$

$$K_{9}=K_{1}K_{8}K_{12}$$

##### **2.5.3 Irreversible Triple Substrate Kinetics with Activation**

Assuming the final product-forming step is irreversible and the intermediate steps are reversible, and that the binding order of metabolites is not important. The binding of the activator is required for the binding of any metabolite.

$$K_{1}=\frac{\left[ E\alpha A \right]}{\left[ E\alpha\right]\left[ A \right]}\to\left[ E\alpha A \right]=K_{1}\left[ E\alpha\right]\left[ A \right]\to\left[ E\alpha A \right]=K_{1}K_{14}\left[ E \right]\left[ \alpha\right]\left[ A \right]$$

$$K_{2}=\frac{\left[ E\alpha B \right]}{\left[ E\alpha\right]\left[ B \right]}\to\left[ E\alpha B \right]=K_{2}\left[ E\alpha\right]\left[ B \right]\to\left[ E\alpha B \right]=K_{2}K_{14}\left[ E \right]\left[ \alpha\right]\left[ B \right]$$

$$K_{3}=\frac{\left[ E\alpha C \right]}{\left[ E\alpha\right]\left[ C \right]}\to\left[ E\alpha C \right]=K_{3}\left[ E\alpha\right]\left[ C \right]\to\left[ E\alpha C \right]=K_{3}K_{14}\left[ E \right]\left[ \alpha\right]\left[ C \right]$$

$$K_{4}=\frac{\left[ E\alpha AC \right]}{\left[ E\alpha C \right]\left[ A \right]}\to\left[ E\alpha AC \right]=K_{4}\left[ E\alpha C \right]\left[ A \right]\to\left[ E\alpha AC \right]=K_{3}K_{4}\left[ E\alpha\right]\left[ A \right]\left[ C \right]\to\left[ E\alpha AC \right]=K_{3}K_{4}K_{14}\left[ E \right]\left[ \alpha\right]\left[ A \right]\left[ C \right]$$

$$K_{5}=\frac{\left[ E\alpha BC \right]}{\left[ E\alpha C \right]\left[ B \right]}\to\left[ E\alpha BC \right]=K_{5}\left[ E\alpha C \right]\left[ B \right]\to\left[ E\alpha BC \right]=K_{3}K_{5}\left[ E\alpha\right]\left[ B \right]\left[ C \right]\to\left[ E\alpha BC \right]=K_{3}K_{5}K_{14}\left[ E \right]\left[ \alpha\right]\left[ B \right]\left[ C \right]$$

$$K_{6}=\frac{\left[ E\alpha BC \right]}{\left[ E\alpha B \right]\left[ C \right]}\to\left[ E\alpha BC \right]=K_{6}\left[ E\alpha B \right]\left[ C \right]\to\left[ E\alpha BC \right]=K_{2}K_{6}\left[ E\alpha\right]\left[ B \right]\left[ C \right]\to\left[ E\alpha BC \right]=K_{2}K_{6}K_{14}\left[ E \right]\left[ \alpha\right]\left[ B \right]\left[ C \right]$$

$$\therefore K_{2}K_{6}=K_{3}K_{5}$$

$$K_{7}=\frac{\left[ E\alpha AB \right]}{\left[ E\alpha B \right]\left[ A \right]}\to\left[ E\alpha AB \right]=K_{7}\left[ E\alpha B \right]\left[ A \right]\to\left[ E\alpha AB \right]=K_{2}K_{7}\left[ E\alpha\right]\left[ A \right]\left[ B \right]\to\left[ E\alpha AB \right]=K_{2}K_{7}K_{14}\left[ E \right]\left[ \alpha\right]\left[ A \right]\left[ B \right]$$

$$K_{8}=\frac{\left[ E\alpha AB \right]}{\left[ E\alpha A \right]\left[ B \right]}\to\left[ E\alpha AB \right]=K_{8}\left[ E\alpha A \right]\left[ B \right]\to\left[ E\alpha AB \right]=K_{1}K_{8}\left[ E\alpha\right]\left[ A \right]\left[ B \right]\to\left[ E\alpha AB \right]=K_{1}K_{8}K_{14}\left[ E \right]\left[ \alpha\right]\left[ A \right]\left[ B \right]$$

$$\therefore K_{1}K_{8}=K_{2}K_{7}$$

$$K_{9}=\frac{\left[ E\alpha AC \right]}{\left[ E\alpha A \right]\left[ C \right]}\to\left[ E\alpha AC \right]=K_{9}\left[ E\alpha A \right]\left[ C \right]\to\left[ E\alpha AC \right]=K_{1}K_{9}\left[ E\alpha\right]\left[ A \right]\left[ C \right]\to\left[ E\alpha AC \right]=K_{1}K_{9}K_{14}\left[ E \right]\left[ \alpha\right]\left[ A \right]\left[ C \right]$$

$$\therefore K_{1}K_{9}=K_{3}K_{4}$$

$$K_{10}=\frac{\left[ E\alpha ABC \right]}{\left[ E\alpha AC \right]\left[ B \right]}\to\left[ E\alpha ABC \right]=K_{10}\left[ E\alpha AC \right]\left[ B \right]\to\left[ E\alpha ABC \right]=K_{1}K_{9}K_{10}\left[ E\alpha\right]\left[ A \right]\left[ B \right]\left[ C \right]\to\left[ E\alpha ABC \right]=K_{1}K_{9}K_{10}K_{14}\left[ E \right]\left[ \alpha\right]\left[ A \right]\left[ B \right]\left[ C \right]$$

$$K_{11}=\frac{\left[ E\alpha ABC \right]}{\left[ E\alpha BC \right]\left[ A \right]}\to\left[ E\alpha ABC \right]=K_{11}\left[ E\alpha BC \right]\left[ A \right]\to\left[ E\alpha ABC \right]=K_{2}K_{6}K_{11}\left[ E\alpha\right]\left[ A \right]\left[ B \right]\left[ C \right]\to\left[ E\alpha ABC \right]=K_{2}K_{6}K_{11}K_{14}\left[ E \right]\left[ \alpha\right]\left[ A \right]\left[ B \right]\left[ C \right]$$

$$K_{12}=\frac{\left[ E\alpha ABC \right]}{\left[ E\alpha AB \right]\left[ C \right]}\to\left[ E\alpha ABC \right]=K_{12}\left[ E\alpha AB \right]\left[ C \right]\to\left[ E\alpha ABC \right]=K_{1}K_{8}K_{12}\left[ E\alpha\right]\left[ A \right]\left[ B \right]\left[ C \right]\to\left[ E\alpha ABC \right]=K_{1}K_{8}K_{12}K_{14}\left[ E \right]\left[ \alpha\right]\left[ A \right]\left[ B \right]\left[ C \right]$$

$$\therefore K_{1}K_{9}K_{10}=K_{2}K_{6}K_{11}=K_{1}K_{8}K_{12}$$

$$K_{14}=\frac{\left[ E\alpha\right]}{\left[ E \right][\alpha]}\to\left[ E\alpha\right]=K_{14}\left[ E \right]\left[ \alpha\right]$$

The reaction rate is then:

$$v_{total}=k_{13}\left[ E\alpha ABC \right]$$

$$v_{total}=k_{13}K_{1}K_{8}K_{12}K_{14}\left[ E \right]\left[ \alpha\right]\left[ A \right]\left[ B \right]\left[ C \right]$$

Since the concentration of complexed and uncomplexed enzymes is difficult to measure, the total enzyme concetration will be used in the formulation:

$$\left[ E \right]_{total}=\left[ E \right]+\left[ E\alpha\right]+\left[ E\alpha A \right]+\left[ E\alpha B \right]+\left[ E\alpha C \right]+\left[ E\alpha AB \right]+\left[ E\alpha AC \right]+\left[ E\alpha BC \right]+\left[ E\alpha ABC \right]$$

$$\left[ E \right]_{total}=\left[ E \right]+K_{14}\left[ E \right]\left[ \alpha\right]+K_{1}K_{14}\left[ E \right]\left[ \alpha\right]\left[ A \right]+K_{2}K_{14}\left[ E \right]\left[ \alpha\right]\left[ B \right]+K_{3}K_{14}\left[ E \right]\left[ \alpha\right]\left[ C \right]+K_{2}K_{7}K_{14}\left[ E \right]\left[ \alpha\right]\left[ A \right]\left[ B \right]+K_{1}K_{9}K_{14}\left[ E \right]\left[ \alpha\right]\left[ A \right]\left[ C \right]+K_{2}K_{6}K_{14}\left[ E \right]\left[ \alpha\right]\left[ B \right]\left[ C \right]+K_{1}K_{8}K_{12}K_{14}\left[ E \right]\left[ \alpha\right]\left[ A \right]\left[ B \right]\left[ C \right]$$

$$\left[ E \right]_{total}=\left[ E \right]\left( 1+K_{14}\left[ \alpha\right]+K_{1}K_{14}\left[ \alpha\right]\left[ A \right]+K_{2}K_{14}\left[ \alpha\right]\left[ B \right]+K_{3}K_{14}\left[ \alpha\right]\left[ C \right]+K_{2}K_{7}K_{14}\left[ \alpha\right]\left[ A \right]\left[ B \right]+K_{1}K_{9}K_{14}\left[ \alpha\right]\left[ A \right]\left[ C \right]+K_{2}K_{6}K_{14}\left[ \alpha\right]\left[ B \right]\left[ C \right]+K_{1}K_{8}K_{12}K_{14}\left[ \alpha\right]\left[ A \right]\left[ B \right]\left[ C \right] \right)$$

$$\frac{\left[ E \right]_{total}}{1+K_{14}\left[ \alpha\right]+K_{1}K_{14}\left[ \alpha\right]\left[ A \right]+K_{2}K_{14}\left[ \alpha\right]\left[ B \right]+K_{3}K_{14}\left[ \alpha\right]\left[ C \right]+K_{2}K_{7}K_{14}\left[ \alpha\right]\left[ A \right]\left[ B \right]+K_{1}K_{9}K_{14}\left[ \alpha\right]\left[ A \right]\left[ C \right]+K_{2}K_{6}K_{14}\left[ \alpha\right]\left[ B \right]\left[ C \right]+K_{1}K_{8}K_{12}K_{14}\left[ \alpha\right]\left[ A \right]\left[ B \right]\left[ C \right]}=\left[ E \right]$$

Therefore, the reaction rate is given as below after combining kinetic parameters into as few as possible:

$$\boldsymbol{v}_{\boldsymbol{total}}\boldsymbol{=}\frac{\boldsymbol{K}_{\boldsymbol{1}}\left[ \boldsymbol{\alpha} \right]\left[ \boldsymbol{A} \right]\left[ \boldsymbol{B} \right]\left[ \boldsymbol{C} \right]}{\boldsymbol{1+}\boldsymbol{K}_{\boldsymbol{2}}\left[ \boldsymbol{\alpha} \right]\boldsymbol{+}\boldsymbol{K}_{\boldsymbol{3}}\left[ \boldsymbol{\alpha} \right]\left[ \boldsymbol{A} \right]\boldsymbol{+}\boldsymbol{K}_{\boldsymbol{4}}\left[ \boldsymbol{\alpha} \right]\left[ \boldsymbol{B} \right]\boldsymbol{+}\boldsymbol{K}_{\boldsymbol{5}}\left[ \boldsymbol{\alpha} \right]\left[ \boldsymbol{C} \right]\boldsymbol{+}\boldsymbol{K}_{\boldsymbol{6}}\left[ \boldsymbol{\alpha} \right]\left[ \boldsymbol{A} \right]\left[ \boldsymbol{B} \right]\boldsymbol{+}\boldsymbol{K}_{\boldsymbol{7}}\left[ \boldsymbol{\alpha} \right]\left[ \boldsymbol{A} \right]\left[ \boldsymbol{C} \right]\boldsymbol{+}\boldsymbol{K}_{\boldsymbol{8}}\left[ \boldsymbol{\alpha} \right]\left[ \boldsymbol{B} \right]\left[ \boldsymbol{C} \right]\boldsymbol{+}\boldsymbol{K}_{\boldsymbol{9}}\left[ \boldsymbol{\alpha} \right]\left[ \boldsymbol{A} \right]\left[ \boldsymbol{B} \right]\left[ \boldsymbol{C} \right]}$$

$$K_{1}=\left[ \boldsymbol{E} \right]_{\boldsymbol{total}}k_{13}K_{1}K_{8}K_{12}K_{14}$$

$$K_{2}=K_{14}$$

$$K_{3}=K_{1}K_{14}$$

$$K_{4}=K_{2}K_{14}$$

$$K_{5}=K_{3}K_{14}$$

$$K_{6}=K_{2}K_{7}K_{14}$$

$$K_{7}=K_{1}K_{9}K_{14}$$

$$K_{8}=K_{2}K_{6}K_{14}$$

$$K_{9}=K_{1}K_{8}K_{12}K_{14}$$

#### **2.6 Reversible Triple Substrate Kinetics**

Reaction kinetics of reversible substrate kinetics are shown below.

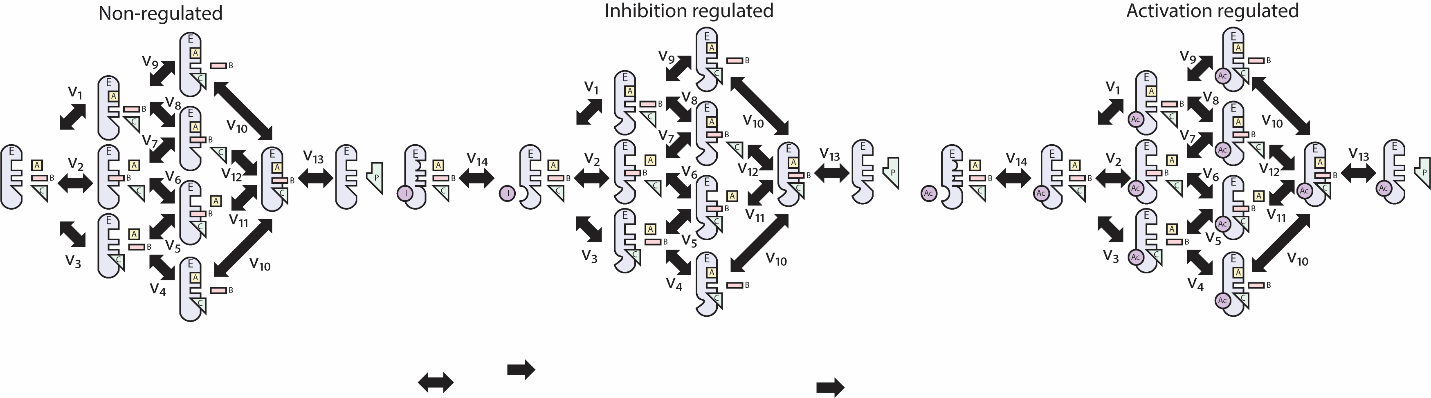

##### **2.6.1 Reversible Triple Substrate Kinetics**

Assumes all reaction steps are reversible and have some equilibrium constant $K_{j}$.

$$K_{1}=\frac{\left[ EA \right]}{\left[ E \right]\left[ A \right]}\to\left[ EA \right]=K_{1}\left[ E \right]\left[ A \right]$$

$$K_{2}=\frac{\left[ EB \right]}{\left[ E \right]\left[ B \right]}\to\left[ EB \right]=K_{2}\left[ E \right]\left[ B \right]$$

$$K_{3}=\frac{\left[ EC \right]}{\left[ E \right]\left[ C \right]}\to\left[ EC \right]=K_{3}\left[ E \right]\left[ C \right]$$

$$K_{4}=\frac{\left[ EAC \right]}{\left[ EC \right]\left[ A \right]}\to\left[ EAC \right]=K_{4}\left[ EC \right]\left[ A \right]\to\left[ EAC \right]=K_{3}K_{4}\left[ E \right]\left[ A \right]\left[ C \right]$$

$$K_{5}=\frac{\left[ EBC \right]}{\left[ EC \right]\left[ B \right]}\to\left[ EBC \right]=K_{5}\left[ EC \right]\left[ B \right]\to\left[ EBC \right]=K_{3}K_{5}\left[ E \right]\left[ B \right]\left[ C \right]$$

$$K_{6}=\frac{\left[ EBC \right]}{\left[ EB \right]\left[ C \right]}\to\left[ EBC \right]=K_{6}\left[ EB \right]\left[ C \right]\to\left[ EBC \right]=K_{2}K_{6}\left[ E \right]\left[ B \right]\left[ C \right]$$

$$\therefore K_{2}K_{6}=K_{3}K_{5}$$

$$K_{7}=\frac{\left[ EAB \right]}{\left[ EB \right]\left[ A \right]}\to\left[ EAB \right]=K_{7}\left[ EB \right]\left[ A \right]\to K_{2}K_{7}\left[ E \right]\left[ A \right]\left[ B \right]$$

$$K_{8}=\frac{\left[ EAB \right]}{\left[ EA \right]\left[ B \right]}\to\left[ EAB \right]=K_{8}\left[ EA \right]\left[ B \right]\to K_{1}K_{8}\left[ E \right]\left[ A \right]\left[ B \right]$$

$$\therefore K_{1}K_{8}=K_{2}K_{7}$$

$$K_{9}=\frac{\left[ EAC \right]}{\left[ EA \right]\left[ C \right]}\to\left[ EAC \right]=K_{9}\left[ EA \right]\left[ C \right]\to\left[ EAC \right]=K_{1}K_{9}\left[ E \right]\left[ A \right]\left[ C \right]$$

$$\therefore K_{1}K_{9}=K_{3}K_{4}$$

$$K_{10}=\frac{\left[ EABC \right]}{\left[ EAC \right]\left[ B \right]}\to\left[ EABC \right]=K_{10}\left[ EAC \right]\left[ B \right]\to\left[ EABC \right]=K_{1}K_{9}K_{10}\left[ E \right]\left[ A \right]\left[ B \right]\left[ C \right]$$

$$K_{11}=\frac{\left[ EABC \right]}{\left[ EBC \right]\left[ A \right]}\to\left[ EABC \right]=K_{11}\left[ EBC \right]\left[ A \right]\to\left[ EABC \right]=K_{2}K_{6}K_{11}\left[ E \right]\left[ A \right]\left[ B \right]\left[ C \right]$$

$$K_{12}=\frac{\left[ EABC \right]}{\left[ EAB \right]\left[ C \right]}\to\left[ EABC \right]=K_{12}\left[ EAB \right]\left[ C \right]\to\left[ EABC \right]=K_{1}K_{8}K_{12}\left[ E \right]\left[ A \right]\left[ B \right]\left[ C \right]$$

$$\therefore K_{1}K_{9}K_{10}=K_{2}K_{6}K_{11}=K_{1}K_{8}K_{12}$$

Therefore the reaction rate is:

$$v_{total}=k_{13}\left[ EABC \right]-k_{13}^{'}\left[ E \right]\left[ P \right]$$

$$v_{total}=K_{13}K_{1}K_{8}K_{12}\left[ E \right]\left[ A \right]\left[ B \right]\left[ C \right]-k_{13}^{'}\left[ E \right]\left[ P \right]$$

Since the concentration of uncomplexed or complexed enzymes cannot be measured, we will use the total enzyme concetration:

$$\left[ E \right]_{total}=\left[ E \right]+\left[ EA \right]+\left[ EB \right]+\left[ EC \right]+\left[ EAB \right]+\left[ EAC \right]+\left[ EBC \right]+\left[ EABC \right]$$

$$\left[ E \right]_{total}=\left[ E \right]+K_{1}\left[ E \right]\left[ A \right]+K_{2}\left[ E \right]\left[ B \right]+K_{3}\left[ E \right]\left[ C \right]+K_{1}K_{8}\left[ E \right]\left[ A \right]\left[ B \right]+K_{1}K_{9}\left[ E \right]\left[ A \right]\left[ C \right]+K_{2}K_{6}\left[ E \right]\left[ B \right]\left[ C \right]+K_{1}K_{8}K_{12}\left[ E \right]\left[ A \right]\left[ B \right]\left[ C \right]$$

$$\left[ E \right]_{total}=\left[ E \right]\left( 1+K_{1}\left[ A \right]+K_{2}\left[ B \right]+K_{3}\left[ C \right]+K_{1}K_{8}\left[ A \right]\left[ B \right]+K_{1}K_{9}\left[ A \right]\left[ C \right]+K_{2}K_{6}\left[ B \right]\left[ C \right]+K_{1}K_{8}K_{12}\left[ A \right]\left[ B \right]\left[ C \right] \right)$$

$$\left[ E \right]=\frac{\left[ E \right]_{total}}{1+K_{1}\left[ A \right]+K_{2}\left[ B \right]+K_{3}\left[ C \right]+K_{1}K_{8}\left[ A \right]\left[ B \right]+K_{1}K_{9}\left[ A \right]\left[ C \right]+K_{2}K_{6}\left[ B \right]\left[ C \right]+K_{1}K_{8}K_{12}\left[ A \right]\left[ B \right]\left[ C \right]}$$

Therefore, the total reaction rate is the:

$$\frac{k_{13}K_{1}K_{8}K_{12}\left[ E \right]_{total}\left[ A \right]\left[ B \right]\left[ C \right]-k_{13}^{'}\left[ E \right]_{total}\left[ P \right]}{1+K_{1}\left[ A \right]+K_{2}\left[ B \right]+K_{3}\left[ C \right]+K_{1}K_{8}\left[ A \right]\left[ B \right]+K_{1}K_{9}\left[ A \right]\left[ C \right]+K_{2}K_{6}\left[ B \right]\left[ C \right]+K_{1}K_{8}K_{12}\left[ A \right]\left[ B \right]\left[ C \right]}$$

##### **Reducing the number of kinetic parameters gives:**

$$\boldsymbol{v}_{\boldsymbol{total}}\boldsymbol{=}\frac{\boldsymbol{K}_{\boldsymbol{1}}\left[ \boldsymbol{A} \right]\left[ \boldsymbol{B} \right]\left[ \boldsymbol{C} \right]\boldsymbol{-}\boldsymbol{K}_{\boldsymbol{2}}\left[ \boldsymbol{P} \right]}{\boldsymbol{1+}\boldsymbol{K}_{\boldsymbol{3}}\left[ \boldsymbol{A} \right]\boldsymbol{+}\boldsymbol{K}_{\boldsymbol{4}}\left[ \boldsymbol{B} \right]\boldsymbol{+}\boldsymbol{K}_{\boldsymbol{5}}\left[ \boldsymbol{C} \right]\boldsymbol{+}\boldsymbol{K}_{\boldsymbol{6}}\left[ \boldsymbol{A} \right]\left[ \boldsymbol{B} \right]\boldsymbol{+}\boldsymbol{K}_{\boldsymbol{7}}\left[ \boldsymbol{A} \right]\left[ \boldsymbol{C} \right]\boldsymbol{+}\boldsymbol{K}_{\boldsymbol{8}}\left[ \boldsymbol{B} \right]\left[ \boldsymbol{C} \right]\boldsymbol{+}\boldsymbol{K}_{\boldsymbol{9}}\left[ \boldsymbol{A} \right]\left[ \boldsymbol{B} \right]\left[ \boldsymbol{C} \right]}$$

$$K_{1}=k_{13}K_{1}K_{8}K_{12}\left[ E \right]_{total}$$

$$K_{2}=K_{1}K_{8}K_{12}\left[ E \right]_{total}$$

$$K_{3}=K_{1}$$

$$K_{4}\boldsymbol{=}K_{2}$$

$$K_{5}=K_{3}$$

$$K_{6}=K_{1}K_{8}$$

$$K_{7}=K_{1}K_{9}$$

$$K_{8}=K_{2}K_{6}$$

$$K_{8}=K_{1}K_{8}K_{12}$$

##### **2.6.2 Reversible Triple Substrate Kinetics with Inhibition**

Assuming the final product-forming step is irreversible and the intermediate steps are reversible, and that the binding order of metabolites is not important. The binding of the inhibitor is incomparable with the binding of any metabolite.

$$K_{1}=\frac{\left[ EA \right]}{\left[ E \right]\left[ A \right]}\to\left[ EA \right]=K_{1}\left[ E \right]\left[ A \right]$$

$$K_{2}=\frac{\left[ EB \right]}{\left[ E \right]\left[ B \right]}\to\left[ EB \right]=K_{2}\left[ E \right]\left[ B \right]$$

$$K_{3}=\frac{\left[ EC \right]}{\left[ E \right]\left[ C \right]}\to\left[ EC \right]=K_{3}\left[ E \right]\left[ C \right]$$

$$K_{4}=\frac{\left[ EAC \right]}{\left[ EC \right]\left[ A \right]}\to\left[ EAC \right]=K_{4}\left[ EC \right]\left[ A \right]\to\left[ EAC \right]=K_{3}K_{4}\left[ E \right]\left[ A \right]\left[ C \right]$$

$$K_{5}=\frac{\left[ EBC \right]}{\left[ EC \right]\left[ B \right]}\to\left[ EBC \right]=K_{5}\left[ EC \right]\left[ B \right]\to\left[ EBC \right]=K_{3}K_{5}\left[ E \right]\left[ B \right]\left[ C \right]$$

$$K_{6}=\frac{\left[ EBC \right]}{\left[ EB \right]\left[ C \right]}\to\left[ EBC \right]=K_{6}\left[ EB \right]\left[ C \right]\to\left[ EBC \right]=K_{2}K_{6}\left[ E \right]\left[ B \right]\left[ C \right]$$

$$\therefore K_{2}K_{6}=K_{3}K_{5}$$

$$K_{7}=\frac{\left[ EAB \right]}{\left[ EB \right]\left[ A \right]}\to\left[ EAB \right]=K_{7}\left[ EB \right]\left[ A \right]\to\left[ EAB \right]=K_{2}K_{7}\left[ E \right]\left[ A \right]\left[ B \right]$$

$$K_{8}=\frac{\left[ EAB \right]}{\left[ EA \right]\left[ B \right]}\to\left[ EAB \right]=K_{8}\left[ EA \right]\left[ B \right]\to\left[ EAB \right]=K_{1}K_{8}\left[ E \right]\left[ A \right]\left[ B \right]$$

$$\therefore K_{1}K_{8}=K_{2}K_{7}$$

$$K_{9}=\frac{\left[ EAC \right]}{\left[ EA \right]\left[ C \right]}\to\left[ EAC \right]=K_{9}\left[ EA \right]\left[ C \right]\to\left[ EAC \right]=K_{1}K_{9}\left[ E \right]\left[ A \right]\left[ C \right]$$

$$\therefore K_{1}K_{9}=K_{3}K_{4}$$

$$K_{10}=\frac{\left[ EABC \right]}{\left[ EAC \right]\left[ B \right]}\to\left[ EABC \right]=K_{10}\left[ EAC \right]\left[ B \right]\to\left[ EABC \right]=K_{1}K_{9}K_{10}\left[ E \right]\left[ A \right]\left[ B \right]\left[ C \right]$$

$$K_{11}=\frac{\left[ EABC \right]}{\left[ EBC \right]\left[ A \right]}\to\left[ EABC \right]=K_{11}\left[ EBC \right]\left[ A \right]\to\left[ EABC \right]=K_{2}K_{6}K_{11}\left[ E \right]\left[ A \right]\left[ B \right]\left[ C \right]$$

$$K_{12}=\frac{\left[ EABC \right]}{\left[ EAB \right]\left[ C \right]}\to\left[ EABC \right]=K_{12}\left[ EAB \right]\left[ C \right]\to\left[ EABC \right]=K_{1}K_{8}K_{12}\left[ E \right]\left[ A \right]\left[ B \right]\left[ C \right]$$

$$\therefore K_{1}K_{9}K_{10}=K_{2}K_{6}K_{11}=K_{1}K_{8}K_{12}$$

$$K_{14}=\frac{\left[ E \right]\left[ I \right]}{\left[ EI \right]}\to\left[ EI \right]=\frac{1}{K_{14}}\left[ E \right]\left[ I \right]$$

Then:

$$v_{total}=k_{13}\left[ EABC \right]-k_{13}^{'}\left[ E \right]\left[ P \right]$$

$$v_{total}=k_{13}K_{1}K_{8}K_{12}\left[ E \right]\left[ A \right]\left[ B \right]\left[ C \right]-k_{13}^{'}\left[ E \right]\left[ P \right]$$

$$v_{total}=\left[ E \right]\left( k_{13}K_{1}K_{8}K_{12}\left[ A \right]\left[ B \right]\left[ C \right]-k_{13}^{'}\left[ P \right] \right)$$

Since the concentration of uncomplexed or complexed enzymes is difficult to measure, the total enzyme concentration must be used.

$$\left[ E \right]_{total}=\left[ E \right]+\left[ EA \right]+\left[ EB \right]+\left[ EC \right]+\left[ EI \right]+\left[ EAB \right]+\left[ EAC \right]+\left[ EBC \right]+\left[ EABC \right]$$

$$\left[ E \right]_{total}=\left[ E \right]+K_{1}\left[ E \right]\left[ A \right]+K_{2}\left[ E \right]\left[ B \right]+K_{3}\left[ E \right]\left[ C \right]+\frac{1}{K_{14}}\left[ E \right]\left[ I \right]+K_{1}K_{8}\left[ E \right]\left[ A \right]\left[ B \right]+K_{3}K_{4}\left[ E \right]\left[ A \right]\left[ C \right]+K_{2}K_{6}\left[ E \right]\left[ B \right]\left[ C \right]+K_{1}K_{8}K_{12}\left[ E \right]\left[ A \right]\left[ B \right]\left[ C \right]$$

$$\left[ E \right]_{total}=\left[ E \right]\left( 1+K_{1}\left[ A \right]+K_{2}\left[ B \right]+K_{3}\left[ C \right]+\frac{1}{K_{14}}\left[ I \right]+K_{1}K_{8}\left[ A \right]\left[ B \right]+K_{3}K_{4}\left[ A \right]\left[ C \right]+K_{2}K_{6}\left[ B \right]\left[ C \right]+K_{1}K_{8}K_{12}\left[ A \right]\left[ B \right]\left[ C \right] \right)$$

$$\left[ E \right]=\frac{\left[ E \right]_{total}}{1+K_{1}\left[ A \right]+K_{2}\left[ B \right]+K_{3}\left[ C \right]+\frac{1}{K_{14}}\left[ I \right]+K_{1}K_{8}\left[ A \right]\left[ B \right]+K_{3}K_{4}\left[ A \right]\left[ C \right]+K_{2}K_{6}\left[ B \right]\left[ C \right]+K_{1}K_{8}K_{12}\left[ A \right]\left[ B \right]\left[ C \right]}$$

Therefore, the reaction rate is given as:

$$v_{total}=\frac{k_{13}K_{1}K_{8}K_{12}\left[ E \right]_{total}\left[ A \right]\left[ B \right]\left[ C \right]-k_{13}^{'}\left[ E \right]_{total}\left[ P \right]}{1+K_{1}\left[ A \right]+K_{2}\left[ B \right]+K_{3}\left[ C \right]+\frac{1}{K_{14}}\left[ I \right]+K_{1}K_{8}\left[ A \right]\left[ B \right]+K_{3}K_{4}\left[ A \right]\left[ C \right]+K_{2}K_{6}\left[ B \right]\left[ C \right]+K_{1}K_{8}K_{12}\left[ A \right]\left[ B \right]\left[ C \right]}$$

Ignoring the subscripts previously defined and seeking to minimize the number of kinetic parameters, this gives:

$$\boldsymbol{v}_{\boldsymbol{total}}\boldsymbol{=}\frac{\boldsymbol{K}_{\boldsymbol{1}}\left[ \boldsymbol{A} \right]\left[ \boldsymbol{B} \right]\left[ \boldsymbol{C} \right]\boldsymbol{-}\boldsymbol{K}_{\boldsymbol{2}}\left[ \boldsymbol{P} \right]}{\boldsymbol{1+}\boldsymbol{K}_{\boldsymbol{3}}\left[ \boldsymbol{A} \right]\boldsymbol{+}\boldsymbol{K}_{\boldsymbol{4}}\left[ \boldsymbol{B} \right]\boldsymbol{+}\boldsymbol{K}_{\boldsymbol{5}}\left[ \boldsymbol{C} \right]\boldsymbol{+}\boldsymbol{K}_{\boldsymbol{6}}\left[ \boldsymbol{I} \right]\boldsymbol{+}\boldsymbol{K}_{\boldsymbol{7}}\left[ \boldsymbol{A} \right]\left[ \boldsymbol{B} \right]\boldsymbol{+}\boldsymbol{K}_{\boldsymbol{8}}\left[ \boldsymbol{A} \right]\left[ \boldsymbol{C} \right]\boldsymbol{+}\boldsymbol{K}_{\boldsymbol{9}}\left[ \boldsymbol{B} \right]\left[ \boldsymbol{C} \right]\boldsymbol{+}\boldsymbol{K}_{\boldsymbol{10}}\left[ \boldsymbol{A} \right]\left[ \boldsymbol{B} \right]\left[ \boldsymbol{C} \right]}$$

$$K_{1}=k_{13}K_{1}K_{8}K_{12}\left[ \boldsymbol{E} \right]_{\boldsymbol{total}}$$

$$K_{2}=k_{13}^{'}K_{1}K_{8}K_{12}\left[ \boldsymbol{E} \right]_{\boldsymbol{total}}$$

$$K_{3}=K_{1}$$

$$K_{4}=K_{2}$$

$$K_{5}=K_{3}$$

$$K_{6}=\frac{1}{K_{14}}$$

$$K_{7}=K_{1}K_{8}$$

$$K_{8}=K_{3}K_{4}$$

$$K_{9}=K_{2}K_{6}$$

$$K_{10}=K_{1}K_{8}K_{12}$$

##### **2.6.3 Reversible Triple Substrate Kinetics with Activation**

Assuming the final product-forming step is irreversible and the intermediate steps are reversible, and that the binding order of metabolites is not important. The binding of the activator is required for the binding of any metabolite.

$$K_{1}=\frac{\left[ E\alpha A \right]}{\left[ E\alpha\right]\left[ A \right]}\to\left[ E\alpha A \right]=K_{1}\left[ E\alpha\right]\left[ A \right]\to\left[ E\alpha A \right]=K_{1}K_{14}\left[ E \right]\left[ \alpha\right]\left[ A \right]$$

$$K_{2}=\frac{\left[ E\alpha B \right]}{\left[ E\alpha\right]\left[ B \right]}\to\left[ E\alpha B \right]=K_{2}\left[ E\alpha\right]\left[ B \right]\to\left[ E\alpha B \right]=K_{2}K_{14}\left[ E \right]\left[ \alpha\right]\left[ B \right]$$

$$K_{3}=\frac{\left[ E\alpha C \right]}{\left[ E\alpha\right]\left[ C \right]}\to\left[ E\alpha C \right]=K_{3}\left[ E\alpha\right]\left[ C \right]\to\left[ E\alpha C \right]=K_{3}K_{14}\left[ E \right]\left[ \alpha\right]\left[ C \right]$$

$$K_{4}=\frac{\left[ E\alpha AC \right]}{\left[ E\alpha C \right]\left[ A \right]}\to\left[ E\alpha AC \right]=K_{4}\left[ E\alpha C \right]\left[ A \right]\to\left[ E\alpha AC \right]=K_{3}K_{4}\left[ E\alpha\right]\left[ A \right]\left[ C \right]\to\left[ E\alpha AC \right]=K_{3}K_{4}K_{14}\left[ E \right]\left[ \alpha\right]\left[ A \right]\left[ C \right]$$

$$K_{5}=\frac{\left[ E\alpha BC \right]}{\left[ E\alpha C \right]\left[ B \right]}\to\left[ E\alpha BC \right]=K_{5}\left[ E\alpha C \right]\left[ B \right]\to\left[ E\alpha BC \right]=K_{3}K_{5}\left[ E\alpha\right]\left[ B \right]\left[ C \right]\to\left[ E\alpha BC \right]=K_{3}K_{5}K_{14}\left[ E \right]\left[ \alpha\right]\left[ B \right]\left[ C \right]$$

$$K_{6}=\frac{\left[ E\alpha BC \right]}{\left[ E\alpha B \right]\left[ C \right]}\to\left[ E\alpha BC \right]=K_{6}\left[ E\alpha B \right]\left[ C \right]\to\left[ E\alpha BC \right]=K_{2}K_{6}\left[ E\alpha\right]\left[ B \right]\left[ C \right]\to\left[ E\alpha BC \right]=K_{2}K_{6}K_{14}\left[ E \right]\left[ \alpha\right]\left[ B \right]\left[ C \right]$$

$$\therefore K_{2}K_{6}=K_{3}K_{5}$$

$$K_{7}=\frac{\left[ E\alpha AB \right]}{\left[ E\alpha B \right]\left[ A \right]}\to\left[ E\alpha AB \right]=K_{7}\left[ E\alpha B \right]\left[ A \right]\to\left[ E\alpha AB \right]=K_{2}K_{7}\left[ E\alpha\right]\left[ A \right]\left[ B \right]\to\left[ E\alpha AB \right]=K_{2}K_{7}K_{14}\left[ E \right]\left[ \alpha\right]\left[ A \right]\left[ B \right]$$

$$K_{8}=\frac{\left[ E\alpha AB \right]}{\left[ E\alpha A \right]\left[ B \right]}\to\left[ E\alpha AB \right]=K_{8}\left[ E\alpha A \right]\left[ B \right]\to\left[ E\alpha AB \right]=K_{1}K_{8}\left[ E\alpha\right]\left[ A \right]\left[ B \right]\to\left[ E\alpha AB \right]=K_{1}K_{8}K_{14}\left[ E \right]\left[ \alpha\right]\left[ A \right]\left[ B \right]$$

$$\therefore K_{1}K_{8}=K_{2}K_{7}$$

$$K_{9}=\frac{\left[ E\alpha AC \right]}{\left[ E\alpha A \right]\left[ C \right]}\to\left[ E\alpha AC \right]=K_{9}\left[ E\alpha A \right]\left[ C \right]\to\left[ E\alpha AC \right]=K_{1}K_{9}\left[ E\alpha\right]\left[ A \right]\left[ C \right]\to\left[ E\alpha AC \right]=K_{1}K_{9}K_{14}\left[ E \right]\left[ \alpha\right]\left[ A \right]\left[ C \right]$$

$$\therefore K_{1}K_{9}=K_{3}K_{4}$$

$$K_{10}=\frac{\left[ E\alpha ABC \right]}{\left[ E\alpha AC \right]\left[ B \right]}\to\left[ E\alpha ABC \right]=K_{10}\left[ E\alpha AC \right]\left[ B \right]\to\left[ E\alpha ABC \right]=K_{1}K_{9}K_{10}\left[ E\alpha\right]\left[ A \right]\left[ B \right]\left[ C \right]\to\left[ E\alpha ABC \right]=K_{1}K_{9}K_{10}K_{14}\left[ E \right]\left[ \alpha\right]\left[ A \right]\left[ B \right]\left[ C \right]$$

$$K_{11}=\frac{\left[ E\alpha ABC \right]}{\left[ E\alpha BC \right]\left[ A \right]}\to\left[ E\alpha ABC \right]=K_{11}\left[ E\alpha BC \right]\left[ A \right]\to\left[ E\alpha ABC \right]=K_{2}K_{6}K_{11}\left[ E\alpha\right]\left[ A \right]\left[ B \right]\left[ C \right]\to\left[ E\alpha ABC \right]=K_{2}K_{6}K_{11}K_{14}\left[ E \right]\left[ \alpha\right]\left[ A \right]\left[ B \right]\left[ C \right]$$

$$K_{12}=\frac{\left[ E\alpha ABC \right]}{\left[ E\alpha AB \right]\left[ C \right]}\to\left[ E\alpha ABC \right]=K_{12}\left[ E\alpha AB \right]\left[ C \right]\to\left[ E\alpha ABC \right]=K_{1}K_{8}K_{12}\left[ E\alpha\right]\left[ A \right]\left[ B \right]\left[ C \right]\to\left[ E\alpha ABC \right]=K_{1}K_{8}K_{12}K_{14}\left[ E \right]\left[ \alpha\right]\left[ A \right]\left[ B \right]\left[ C \right]$$

$$\therefore K_{1}K_{9}K_{10}=K_{2}K_{6}K_{11}=K_{1}K_{8}K_{12}$$

$$K_{14}=\frac{\left[ E\alpha\right]}{\left[ E \right][\alpha]}\to\left[ E\alpha\right]=K_{14}\left[ E \right]\left[ \alpha\right]$$

The reaction rate is then:

$$v_{total}=k_{13}\left[ E\alpha ABC \right]-k_{13}^{'}\left[ E\alpha\right]\left[ P \right]$$

$$v_{total}=k_{13}K_{1}K_{8}K_{12}K_{14}\left[ E \right]\left[ \alpha\right]\left[ A \right]\left[ B \right]\left[ C \right]-k_{13}^{'}K_{14}\left[ E \right]\left[ \alpha\right]\left[ P \right]$$

$$v_{total}=\left[ E \right]\left[ \alpha\right]\left( k_{13}K_{1}K_{8}K_{12}K_{14}\left[ A \right]\left[ B \right]\left[ C \right]-k_{13}^{'}\left[ P \right] \right)$$

Since the concentration of complexed and uncomplexed enzymes is difficult to measure, the total enzyme concetration will be used in the formulation:

$$\left[ E \right]_{total}=\left[ E \right]+\left[ E\alpha\right]+\left[ E\alpha A \right]+\left[ E\alpha B \right]+\left[ E\alpha C \right]+\left[ E\alpha AB \right]+\left[ E\alpha AC \right]+\left[ E\alpha BC \right]+\left[ E\alpha ABC \right]$$

$$\left[ E \right]_{total}=\left[ E \right]+K_{14}\left[ E \right]\left[ \alpha\right]+K_{1}K_{14}\left[ E \right]\left[ \alpha\right]\left[ A \right]+K_{2}K_{14}\left[ E \right]\left[ \alpha\right]\left[ B \right]+K_{3}K_{14}\left[ E \right]\left[ \alpha\right]\left[ C \right]+K_{2}K_{7}K_{14}\left[ E \right]\left[ \alpha\right]\left[ A \right]\left[ B \right]+K_{1}K_{9}K_{14}\left[ E \right]\left[ \alpha\right]\left[ A \right]\left[ C \right]+K_{2}K_{6}K_{14}\left[ E \right]\left[ \alpha\right]\left[ B \right]\left[ C \right]+K_{1}K_{8}K_{12}K_{14}\left[ E \right]\left[ \alpha\right]\left[ A \right]\left[ B \right]\left[ C \right]$$

$$\left[ E \right]_{total}=\left[ E \right]\left( 1+K_{14}\left[ \alpha\right]+K_{1}K_{14}\left[ \alpha\right]\left[ A \right]+K_{2}K_{14}\left[ \alpha\right]\left[ B \right]+K_{3}K_{14}\left[ \alpha\right]\left[ C \right]+K_{2}K_{7}K_{14}\left[ \alpha\right]\left[ A \right]\left[ B \right]+K_{1}K_{9}K_{14}\left[ \alpha\right]\left[ A \right]\left[ C \right]+K_{2}K_{6}K_{14}\left[ \alpha\right]\left[ B \right]\left[ C \right]+K_{1}K_{8}K_{12}K_{14}\left[ \alpha\right]\left[ A \right]\left[ B \right]\left[ C \right] \right)$$

$$\frac{\left[ E \right]_{total}}{1+K_{14}\left[ \alpha\right]+K_{1}K_{14}\left[ \alpha\right]\left[ A \right]+K_{2}K_{14}\left[ \alpha\right]\left[ B \right]+K_{3}K_{14}\left[ \alpha\right]\left[ C \right]+K_{2}K_{7}K_{14}\left[ \alpha\right]\left[ A \right]\left[ B \right]+K_{1}K_{9}K_{14}\left[ \alpha\right]\left[ A \right]\left[ C \right]+K_{2}K_{6}K_{14}\left[ \alpha\right]\left[ B \right]\left[ C \right]+K_{1}K_{8}K_{12}K_{14}\left[ \alpha\right]\left[ A \right]\left[ B \right]\left[ C \right]}=\left[ E \right]$$

Therefore, the reaction rate is given as below after combining kinetic parameters into as few as possible:

$$\boldsymbol{v}_{\boldsymbol{total}}\boldsymbol{=}\frac{\boldsymbol{K}_{\boldsymbol{1}}\left[ \boldsymbol{\alpha} \right]\left[ \boldsymbol{A} \right]\left[ \boldsymbol{B} \right]\left[ \boldsymbol{C} \right]\boldsymbol{-}\boldsymbol{K}_{\boldsymbol{2}}\left[ \boldsymbol{\alpha} \right]\left[ \boldsymbol{P} \right]}{\boldsymbol{1+}\boldsymbol{K}_{\boldsymbol{3}}\left[ \boldsymbol{\alpha} \right]\boldsymbol{+}\boldsymbol{K}_{\boldsymbol{4}}\left[ \boldsymbol{\alpha} \right]\left[ \boldsymbol{A} \right]\boldsymbol{+}\boldsymbol{K}_{\boldsymbol{5}}\left[ \boldsymbol{\alpha} \right]\left[ \boldsymbol{B} \right]\boldsymbol{+}\boldsymbol{K}_{\boldsymbol{6}}\left[ \boldsymbol{\alpha} \right]\left[ \boldsymbol{C} \right]\boldsymbol{+}\boldsymbol{K}_{\boldsymbol{7}}\left[ \boldsymbol{\alpha} \right]\left[ \boldsymbol{A} \right]\left[ \boldsymbol{B} \right]\boldsymbol{+}\boldsymbol{K}_{\boldsymbol{8}}\left[ \boldsymbol{\alpha} \right]\left[ \boldsymbol{A} \right]\left[ \boldsymbol{C} \right]\boldsymbol{+}\boldsymbol{K}_{\boldsymbol{9}}\left[ \boldsymbol{\alpha} \right]\left[ \boldsymbol{B} \right]\left[ \boldsymbol{C} \right]\boldsymbol{+}\boldsymbol{K}_{\boldsymbol{10}}\left[ \boldsymbol{\alpha} \right]\left[ \boldsymbol{A} \right]\left[ \boldsymbol{B} \right]\left[ \boldsymbol{C} \right]}$$

$$K_{1}=k_{13}K_{1}K_{8}K_{12}K_{14}\left[ E \right]_{total}$$

$$K_{2}=k_{13}^{'}K_{1}K_{8}K_{12}K_{14}\left[ E \right]_{total}$$

$$K_{3}=K_{14}$$

$$K_{4}=K_{1}K_{14}$$

$$K_{5}=K_{2}K_{14}$$

$$K_{6}=K_{3}K_{14}$$

$$K_{7}=K_{2}K_{7}K_{14}$$

$$K_{8}=K_{1}K_{9}K_{14}$$

$$K_{9}=K_{2}K_{6}K_{14}$$

$$K_{10}=K_{1}K_{8}K_{12}K_{14}$$

### **3 Fixed or Unfixed Kinetic Parameters**

| $b_{1}$ | $b_{2}$ | $b_{3}$ | $\beta_{1}$ | $\beta_{2}$ | $\beta_{3}$ | $\beta_{4}$ | $\beta_{5}$ | $\beta_{6}$ | $K_{1}=$ | $K_{2}=$ | $K_{3}=$ | $K_{4}=$ | $K_{5}=$ | $K_{6}=$ | $K_{7}=$ | $K_{8}=$ | $K_{9}=$ | $K_{10}=$ |
| --- | --- | --- | --- | --- | --- | --- | --- | --- | --- | --- | --- | --- | --- | --- | --- | --- | --- | --- |
| 1 | 0 | 0 | 1 | 0 | 0 | 0 | 0 | 0 | $K_{1}$ | $K_{2}$ | 0 | 0 | 0 | 0 | 0 | 0 | 0 | 0 |
| 0 | 1 | 0 | 1 | 0 | 0 | 0 | 0 | 0 | $K_{1}$ | $K_{2}$ | $K_{3}$ | 0 | 0 | 0 | 0 | 0 | 0 | 0 |
| 0 | 0 | 1 | 1 | 0 | 0 | 0 | 0 | 0 | $K_{1}$ | $K_{2}$ | $K_{3}$ | 0 | 0 | 0 | 0 | 0 | 0 | 0 |
| 1 | 0 | 0 | 0 | 1 | 0 | 0 | 0 | 0 | $K_{1}$ | $K_{2}$ | $K_{3}$ | $K_{4}$ | 0 | 0 | 0 | 0 | 0 | 0 |
| 0 | 1 | 0 | 0 | 1 | 0 | 0 | 0 | 0 | $K_{1}$ | $K_{2}$ | $K_{3}$ | $K_{4}$ | $K_{5}$ | 0 | 0 | 0 | 0 | 0 |
| 0 | 0 | 1 | 0 | 1 | 0 | 0 | 0 | 0 | $K_{1}$ | $K_{2}$ | $K_{3}$ | $K_{4}$ | $K_{5}$ | 0 | 0 | 0 | 0 | 0 |
| 1 | 0 | 0 | 0 | 0 | 1 | 0 | 0 | 0 | $K_{1}$ | $K_{2}$ | $K_{3}$ | $K_{4}$ | 0 | 0 | 0 | 0 | 0 | 0 |
| 0 | 1 | 0 | 0 | 0 | 1 | 0 | 0 | 0 | $K_{1}$ | $K_{2}$ | $K_{3}$ | $K_{4}$ | $K_{5}$ | 0 | 0 | 0 | 0 | 0 |
| 0 | 0 | 1 | 0 | 0 | 1 | 0 | 0 | 0 | $K_{1}$ | $K_{2}$ | $K_{3}$ | $K_{4}$ | $K_{5}$ | 0 | 0 | 0 | 0 | 0 |
| 1 | 0 | 0 | 0 | 0 | 0 | 1 | 0 | 0 | $K_{1}$ | $K_{2}$ | $K_{3}$ | $K_{4}$ | $K_{5}$ | 0 | 0 | 0 | 0 | 0 |
| 0 | 1 | 0 | 0 | 0 | 0 | 1 | 0 | 0 | $K_{1}$ | $K_{2}$ | $K_{3}$ | $K_{4}$ | $K_{5}$ | $K_{6}$ | 0 | 0 | 0 | 0 |
| 0 | 0 | 1 | 0 | 0 | 0 | 1 | 0 | 0 | $K_{1}$ | $K_{2}$ | $K_{3}$ | $K_{4}$ | $K_{5}$ | $K_{6}$ | 0 | 0 | 0 | 0 |
| 1 | 0 | 0 | 0 | 0 | 0 | 0 | 1 | 0 | $K_{1}$ | $K_{2}$ | $K_{3}$ | $K_{4}$ | $K_{5}$ | $K_{6}$ | $K_{7}$ | $K_{8}$ | 0 | 0 |
| 0 | 1 | 0 | 0 | 0 | 0 | 0 | 1 | 0 | $K_{1}$ | $K_{2}$ | $K_{3}$ | $K_{4}$ | $K_{5}$ | $K_{6}$ | $K_{7}$ | $K_{8}$ | $K_{9}$ | 0 |
| 0 | 0 | 1 | 0 | 0 | 0 | 0 | 1 | 0 | $K_{1}$ | $K_{2}$ | $K_{3}$ | $K_{4}$ | $K_{5}$ | $K_{6}$ | $K_{7}$ | $K_{8}$ | $K_{9}$ | 0 |
| 1 | 0 | 0 | 0 | 0 | 0 | 0 | 0 | 1 | $K_{1}$ | $K_{2}$ | $K_{3}$ | $K_{4}$ | $K_{5}$ | $K_{6}$ | $K_{7}$ | $K_{8}$ | $K_{9}$ | 0 |
| 0 | 1 | 0 | 0 | 0 | 0 | 0 | 0 | 1 | $K_{1}$ | $K_{2}$ | $K_{3}$ | $K_{4}$ | $K_{5}$ | $K_{6}$ | $K_{7}$ | $K_{8}$ | $K_{9}$ | $K_{10}$ |
| 0 | 0 | 1 | 0 | 0 | 0 | 0 | 0 | 1 | $K_{1}$ | $K_{2}$ | $K_{3}$ | $K_{4}$ | $K_{5}$ | $K_{6}$ | $K_{7}$ | $K_{8}$ | $K_{9}$ | $K_{10}$ |
